# The Impact of the Siemens Trio to Prisma Upgrade and Volumetric Navigators on MRI Indices: A Reliability Study with Implications for Longitudinal Study Designs

**DOI:** 10.1101/2020.06.19.162420

**Authors:** Eric Plitman, Aurelie Bussy, Vanessa Valiquette, Alyssa Salaciak, Raihaan Patel, Marie-Lise Béland, Stephanie Tullo, Christine Tardif, M. Natasha Rajah, Jamie Near, Gabriel A. Devenyi, M. Mallar Chakravarty

## Abstract

Many magnetic resonance imaging (MRI) indices are being studied longitudinally to explore topics such as biomarker detection and clinical staging. A pertinent concern to longitudinal work is MRI scanner upgrades. When upgrades occur during the course of a longitudinal MRI neuroimaging investigation, there may be an impact on the compatibility of pre- and post-upgrade measures. Similarly, subject motion is another issue that may be detrimental to longitudinal MRI work; however, embedding volumetric navigators (vNavs) within acquisition sequences has emerged as a technique that allows for prospective motion correction. Our research group recently implemented an upgrade from a Siemens MAGNETOM 3T Trio system to a Siemens MAGNETOM 3T Prisma Fit system. The goals of the current work were to: 1) investigate the impact of this upgrade on commonly used structural imaging measures and proton magnetic resonance spectroscopy indices (“Prisma Upgrade protocol”) and 2) examine structural imaging measures in a sequence with vNavs alongside a standard acquisition sequence (“vNav protocol”). In both protocols, while high reliability was observed for most of the investigated MRI outputs, suboptimal reliability was observed for certain indices. Across the scanner upgrade, increases in frontal, temporal, and cingulate cortical thickness (CT) and thalamus volume, along with decreases in parietal CT, amygdala, globus pallidus, hippocampus, and striatum volumes were observed across the Prisma upgrade, and were linked to increases in signal-to-noise ratios. No significant impact of the upgrade was found in ^1^H-MRS analyses. Further, CT estimates were found to be larger in MPRAGE acquisitions compared to vNav-MPRAGE acquisitions mainly within temporal areas, while the opposite was found mostly in parietal brain regions. The results from this work should be considered in longitudinal study designs and comparable prospective motion correction investigations are warranted in cases of marked head movement.

## 1. Introduction

Magnetic resonance imaging (MRI)-based neurochemical, volumetric, and morphological (e.g. cortical thickness, surface area) indices are often used to explore research questions in neuroscience. For example, existing large-scale MRI studies examining patients with schizophrenia, Alzheimer’s disease, major depressive disorder, and autism spectrum disorder have previously identified neurochemical, volumetric, and morphological abnormalities, as compared to healthy controls (Schmaal et al., 2016; van Erp et al., 2018, 2016; van Rooij et al., 2018). Outside of the context of neuropsychiatric disorders, longitudinal MRI made a significant impact on studies of brain maturation (Raznahan et al., 2014; Reardon et al., 2018) and ageing (Fjell et al., 2015; Tullo et al., 2019; Voineskos et al., 2015). As such, various MRI indices are being investigated to assess their potential role as biomarkers for neuropsychiatric disorders and their utility in clinical staging, prognostication, prediction of illness onset, and the development of normative trajectories, as well as to further the general understanding of neuroscience.

Compared to cross-sectional approaches, longitudinal MRI studies are often considered a more effective strategy for biomarker investigations. Longitudinal brain imaging mitigates potential sources of confounding that may affect cross-sectional studies, such as participant heterogeneity and cohort effects. As a result, indices identified through longitudinal studies as relevant biomarkers in the pathophysiology of various neuropsychiatric disorders can be more readily assigned causality (Cannon et al., 2015; Chincarini et al., 2016; Jack et al., 2013; van Haren et al., 2011).

One concern that is pertinent to the completion of a longitudinal MRI study is the fact that MRI scanners often undergo hardware and software upgrades. It is conceivable that these upgrades may affect the measurement of various relevant MRI indices. When upgrades occur during the course of a longitudinal MRI investigation, there may be an impact on the compatibility of pre- and post-upgrade measures (Lee et al., 2019; Takao et al., 2013). Our research group recently implemented an upgrade for our 3T MRI scanner, from a Siemens MAGNETOM 3T Trio system to a Siemens MAGNETOM 3T Prisma Fit system. This included hardware upgrades to the gradient, radiofrequency, and shimming systems as well as software and applications upgrades (Siemens, 2020).

Another important issue for longitudinal MRI studies (although relevant to all MRI studies) is subject motion. Subject motion during a three-dimensional MRI acquisition is well-known to cause artifacts (e.g. ghosting, blurring) by making k-space sampling inconsistent, which may be detrimental to the quality and utility of the scan (Tisdall et al., 2012). This is an important consideration for longitudinal work, as it is common practice amongst research groups to remove poor quality scans from further analysis. This represents a loss of significant resources, which may often influence the utility of other data collected from that participant over the course of a longitudinal study design. In addition, removing high-motion scans can represent a source of bias within the dataset and unresolved motion has been demonstrated to impact both volumetric and morphological estimates (Bedford et al., 2020; Reuter et al., 2015). The impact of subject motion may be mitigated through multiple approaches, which may be classified as prospective and retrospective. One prospective technique that has become increasingly common involves embedding volumetric navigators (vNavs) (i.e. fast-acquisition low-resolution 3D EPI images) within longer acquisition sequences (Tisdall et al., 2016, 2012). vNavs undergo rapid rigid registration to estimate changes in head position, effectively tracking and updating subject motion during an acquisition. Among the advantages of this approach is its efficiency, as the vNavs are distributed throughout existing sequences within “dead-times”, thus providing benefit at a negligible cost of time, contrast, and intensity. Also, vNav sequences may permit the re-acquisition of repetition times (TRs) within which a large quantity of motion exists. Notably, vNavs have been shown to lead to a reduction in morphometry variation attributed to motion (Tisdall et al., 2016) and are currently being employed in large-scale initiatives where participant motion is known to be problematic (Alexander-Bloch et al., 2016; Pardoe et al., 2016) such as the Adolescent Brain Cognitive Development study (Casey et al., 2018).

The goals of the current work were to: 1) investigate the impact of upgrading from a Siemens 3T Trio system to a Siemens 3T MAGNETOM Prisma system on commonly used volumetric and morphometric structural imaging measures and proton magnetic resonance spectroscopy (^1^H-MRS) indices and 2) examine the reliability and similarity of volumetric and morphometric structural imaging measures in a sequence with vNavs alongside a standard T1-weighted acquisition sequence.

## 2. Methods

### 2.1. Participants

This study received approval by the Research Ethics Board of the Douglas Mental Health University Institute. Written informed consent was acquired from a total of nineteen individuals who completed the screening interview and were subsequently enrolled into the present study. Inclusion criteria were: capacity to consent; age 18-80; and fluency in English or French. Exclusion criteria included contraindications to MRI.

### 2.2. Study Design

This study was conducted from June 2018 to September 2019, and consisted of two protocols (Figure 1). The sample size within each investigation is detailed within Supplementary Table 1.

**Figure 1.**
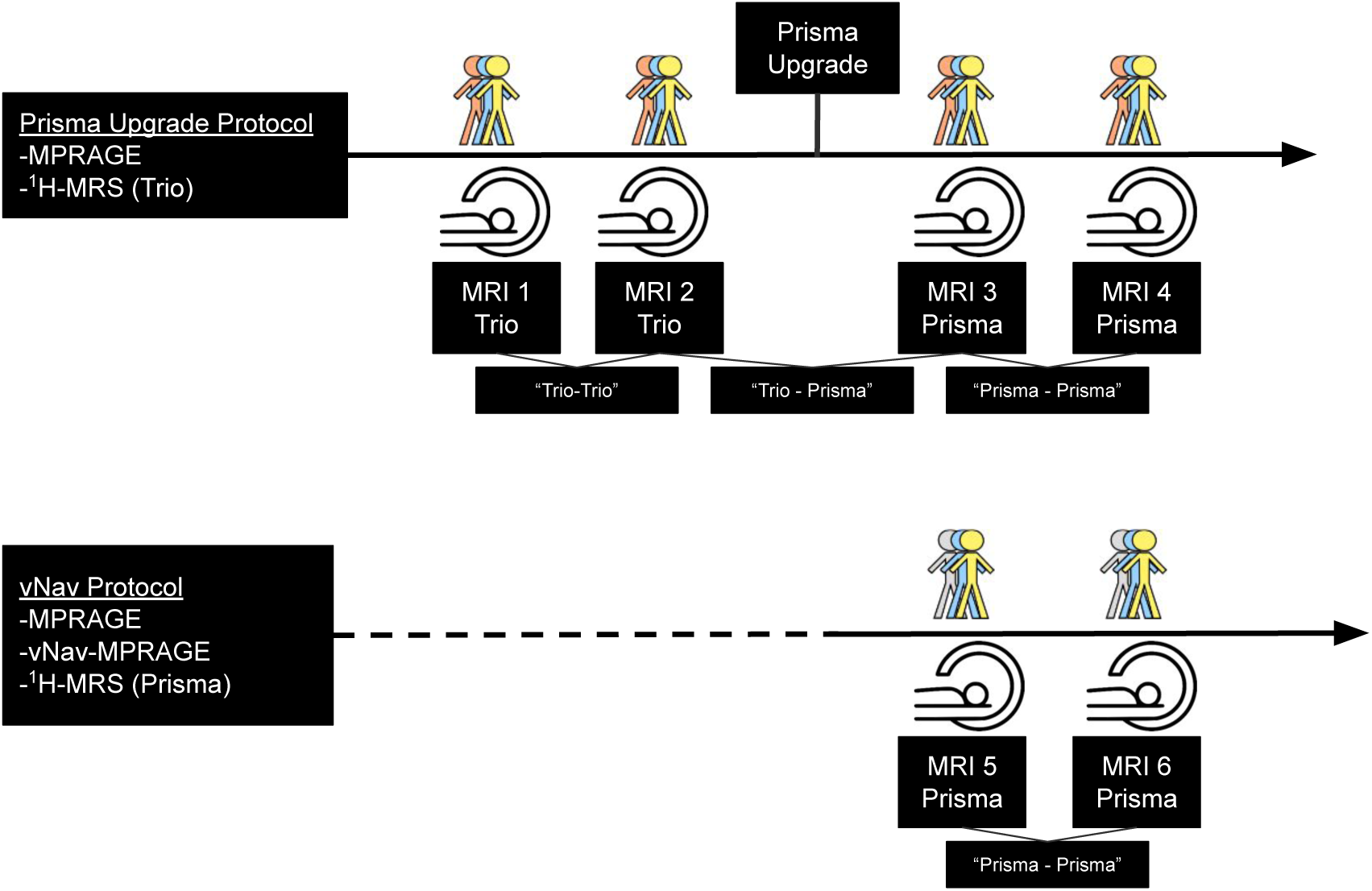
Study design.

#### 2.2.1. Prisma Upgrade Protocol

The first protocol was designed to assess the impact of the Prisma upgrade, and consisted of four MRI visits per subject: two using a Siemens MAGNETOM Trio MRI scanner and two using the Siemens MAGNETOM Prisma MRI Fit scanner following the completion of an upgrade. To merit inclusion of a subject, the participant must have undergone at least one pre-upgrade and one post-upgrade acquisition. In this study protocol, fourteen healthy individuals participated, although one participant failed to complete any post-upgrade scans. Thus, data from thirteen individuals (mean age: 28.62 ± 6.14 years, 6 females) were used for analysis. Nine subjects completed all four MRI visits, one subject completed two pre-upgrade visits and one post-upgrade visit, one subject completed one pre-upgrade visit and two post-upgrade visits, and two subjects completed one pre-upgrade visit and one post-upgrade visit. One participant who completed two pre-upgrade MPRAGE acquisitions only completed one pre-upgrade ^1^H-MRS acquisition.

#### 2.2.2. vNav Protocol

The second study protocol consisted of two MRI visits and was principally designed to: 1) investigate vNav structural sequences alongside standard structural acquisitions and 2) acquire post-upgrade ^1^H-MRS. Due to the latter purpose, we aimed to include as many subjects as possible from the Prisma Upgrade protocol in the vNav protocol. Thirteen individuals (mean age: 27.92 ± 6.63 years, 7 females) participated in the vNav protocol, eight of which also participated in the Prisma Upgrade protocol. MPRAGE acquisitions always occurred prior to vNav-MPRAGE acquisitions within a scanning session. Notably, MPRAGE data acquired within the vNav protocol was used to supplement post-upgrade (i.e. Prisma-Prisma) analyses within the Prisma Upgrade protocol.

### 2.3. Magnetic Resonance Imaging Acquisition

Participants were scanned at the Douglas Mental Health University Institute. In the Prisma Upgrade protocol, scanning procedures during MRI visits 1 and 2 were performed on a 3T Siemens Trio MRI machine using a 32-channel head coil. For MRI visits 3 and 4, scanning procedures were performed on a 3T Siemens MAGNETOM Prisma MRI machine using a 32-channel head coil. In each of these MRI visits, T1-weighted structural images were acquired (MPRAGE, TR=2300ms, TE=2.98ms (Trio) and 2.01ms (Prisma), TI=900ms, flip angle=9°, bandwidth=238Hz/Px (Trio) and 240Hz/Px (Prisma), voxel size=1.0 × 1.0 × 1.0mm, 192 × 240 × 256 matrix). In the vNav protocol, all scanning procedures were performed on a 3T Siemens MAGNETOM Prisma MRI machine. During both of the MRI visits, MPRAGE (parameters noted above) and vNav-MPRAGE (TR=2300ms, TE=2.90 ms, TI=1070ms, flip angle=9°, bandwidth=240Hz/Px, voxel size=1.0 × 1.0 × 1.0mm, 176 × 240 × 256 matrix) were acquired. vNav-MPRAGE sequences were provided by Massachusetts General Hospital.

### 2.4. Magnetic Resonance Spectroscopy Acquisition

In the Prisma Upgrade protocol, ^1^H-MRS was acquired within the left dorsolateral prefrontal cortex (DLPFC) during MRI visits 1 and 2. As part of the vNav protocol, ^1^H-MRS was acquired within the left DLPFC during both of the MRI visits. For ^1^H-MRS scans in both the Prisma Upgrade protocol and the vNav protocol, the MPRAGE anatomical image was used to guide placement of a 2.5 × 2.5 × 1.5 cm^3^ voxel in the left DLPFC. The voxel was first positioned such that when viewed in the sagittal plane, the centre of the MRS voxel in the A/P direction was aligned with the anterior extent of the genu of the corpus callosum. Next, when viewed in the coronal plane at the level of the genu of the corpus callosum, the voxel was positioned at the brain surface, approximately mid-way between the dorsomedial extent and the left-most lateral extent of the brain. To ensure maximal coverage of cortical grey matter in that region, the voxel was rotated about both the scanner y- and x-axes such that the largest face of the voxel (the 2.5 × 2.5 cm^2^ face) was parallel to the surface of the brain. Short echo-time ^1^H-MRS data were acquired using the SPECIAL sequence (Mekle et al., 2009) with TR=3500 ms (Trio) and 3200 ms (Prisma), TE=8.5 ms, 4096 spectral points, 4000 Hz spectral width. 144 (Trio) and 160 (Prisma) water-suppressed averages were acquired using the VAPOR water suppression scheme (Tkác et al., 1999), and an additional 8 non-water-suppressed averages were acquired for use in array coil reconstruction, lineshape correction, and optional signal referencing.

Prior to ^1^H-MRS acquisition, localized shimming was performed on the volume of interest. In the Prisma Upgrade protocol, shimming was performed using the projection-based FASTESTMAP method (Gruetter and Tkác, 2000), while in the vNav protocol, shimming was performed using the fieldmap-based GRE-SHIM method.

### 2.5. Preprocessing

Unless otherwise specified, structural imaging analysis was done in the minc format on a high-performance computing cluster (SciNet). T1-weighted structural images were preprocessed using the minc-bpipe-library pipeline (https://github.com/CobraLab/minc-bpipe-library), which included a signal intensity correction (i.e. N4-correction (Tustison et al., 2010)), and procedures to exclude the neck and skull. ^1^H-MRS analyses included the simulation of a site-specific basis set and data preprocessing performed using the FID-A toolkit (Simpson et al., 2017), the latter of which included removal of motion-corrupted averages, along with frequency and phase drift correction.

### 2.6. Morphological Analysis

Cortical thickness (CT) was estimated using the CIVET processing pipeline (Lerch and Evans, 2005) (version 2.1.0) on N4-corrected (Tustison et al., 2010) T1-weighted inputs (e.g. MPRAGE and vNav-MPRAGE). Images were aligned linearly to the ICBM 152 average template through a twelve-parameter transformation (Collins et al., 1994). Next, images were classified into grey matter, white matter, and cerebrospinal fluid (Zijdenbos et al., 2002). Hemispheres were then modeled as grey matter and white matter surfaces using a deformable model strategy, which generates 4 separate surfaces, each defined by 40,962 vertices (Kim et al., 2005). CT was determined in native space through nonlinear surface-based normalization that uses a midsurface between pial and white matter surfaces. Native-space thicknesses were used in analyses, considering that normalizing for head or brain volume has little relationship to CT and risks introducing noise (Sowell et al., 2007). The mean CT was estimated in 31 bilateral regions of interest (ROIs), as defined by the Desikan-Killiany-Tourville atlas (Klein and Tourville, 2012), using unsmoothed “tlaplace” outputs.

In addition, for complementary analyses, CT was estimated using the recon-all pipeline from FreeSurfer (version 6.0) (Dale et al., 1999; Fischl et al., 1999) on native T1-weighted images (e.g. MPRAGE and vNav-MPRAGE) in the DICOM file format. FreeSurfer methodology and documentation has been previously detailed and is available at: http://surfer.nmr.mgh.harvard.edu. Parcellation of reconstructed surfaces from FreeSurfer rendered CT estimates for each participant in ROIs defined by the Desikan-Killiany-Tourville atlas (Klein and Tourville, 2012). For consistency in nomenclature with CIVET, lateral occipital, pars opercularis, pars orbitalis, and pars triangularis ROIs are referred to as the inferior occipital cortex, lateral frontal opercularis, lateral frontal orbitalis, and lateral frontal triangularis, respectively.

### 2.7. Volumetric Analysis

Fully-automated segmentations of neuroanatomical structure volumes (SVs) (e.g. amygdala, globus pallidus, hippocampus, striatum, thalamus) were carried out using the Multiple Automatically Generated Templates (MAGeT-Brain) algorithm (Chakravarty et al., 2013; Pipitone et al., 2014). This technique is a modified multi-atlas segmentation technique designed to use a limited number of high-quality manually segmented atlases as input. Using 5 high-resolution brain images, different atlases were manually segmented (https://github.com/CoBrALab/atlases). Resultantly, globus pallidus, striatum, and thalamus volumes were examined using 5 subcortical atlases (Tullo et al., 2018) – based on a warped histology-based atlas (Chakravarty et al., 2006) – in addition to using previously used definitions of the amygdala (Treadway et al., 2015) and hippocampus (Amaral et al., 2018; Winterburn et al., 2013). In MAGeT Brain, a subset of the population under study is used as a template library through which the final segmentation is bootstrapped. Twenty-one templates were selected from the overall image pool to ensure optimal segmentation (https://github.com/CoBrALab/documentation/wiki/Best-Templates-for-MAGeT) and to achieve a representative template set. Each subject in the template library is segmented through nonlinear atlas-to-template registration followed by label propagation, yielding 5 unique definitions of the subdivisions for each of the templates. The bootstrapping of the final segmentations through the template library results in 105 candidate labels produced for each subject and labels are then fused using a majority vote to complete the segmentation process. Nonlinear registration was performed using a version of the Advanced Normalization Tools (ANTS) (Avants et al., 2008) (https://github.com/ANTsX/ANTs), a process which has been validated in several of our previous studies (Chakravarty et al., 2013; Makowski et al., 2018; Pipitone et al., 2014; Tullo et al., 2018). Skull-stripped atlas inputs were also used (https://github.com/CoBrALab/atlases) to match the outputs of the pre-processing.

As above, FreeSurfer was used, wherein automated segmentation rendered cortical and subcortical volumes (Fischl et al., 2002). Given the complementary nature of this analysis, only volumes from structures investigated with MAGeT-Brain (e.g. amygdala, globus pallidus [i.e. pallidum], hippocampus, striatum [i.e. sum of caudate and putamen], thalamus [i.e. thalamus proper]), were used.

For CT and SV analyses, outputs were not excluded from analysis on the basis of quality.

### 2.8 Signal-to-Noise Analysis

Signal-to-noise ratios (SNR) were estimated using noise_estimate (Coupé et al., 2009) from native MPRAGE acquisitions in the minc format that were included in the Prisma Upgrade protocol.

### 2.9. ^1^H-MRS Analysis

Glutamate (Glu), glutamine (Gln), Glu + Gln (Glx), myo-inositol (Ins), total choline (Cho, sum of glycerophosphocholine and phosphocholine), N-acetylaspartate (NAA), N-acetylaspartylglutamate (NAAG), total N-acetylaspartate (tNAA, sum of NAA and NAAG), gamma-Aminobutyric acid (GABA), glutathione (GSH), and lactate (Lac) were estimated from preprocessed ^1^H-MRS data using LCModel version 6.3-1H (Provencher, 2001).

Neurometabolite levels were referenced to total creatine (sum of creatine and phosphocreatine) levels. A basis set of neurometabolites, obtained via simulation in FID-A (Simpson et al., 2017) according to the *in vivo* scan parameters, was used. This basis set contained L-alanine, aspartate, creatine, Cr methylene group, GABA, glucose, Glu, Gln, GSH, glycerophosphocholine, glycine, β-hydroxybutyrate, L-lactate, Ins, NAA, NAAG, phosphocholine, phosphocreatine, phosphorylethanolamine, scyllo-inositol, serine, and taurine, as well as LCModel’s default model functions for mobile lipids (Lip) and macromolecules (MM): Lip09, Lip13a, Lip13b, Lip20, MM09, MM12, MM14, MM17, and MM20. In the current work, no cutoffs were employed for %SD, full-width at half maximum (FWHM), or SNR. Notably, FWHM and SNR were calculated using FID-A (Simpson et al., 2017). FWHM represents the FWHM of the water peak in the water unsuppressed scan and SNR is the signal of the NAA peak in the water suppressed scan divided by the standard deviation of the noise between 0 and 2 ppm in the same spectrum.

### 2.10. Statistical Analyses

Analyses were performed in R version 3.5.0 and can be further divided into “test-retest reliability analyses” and “similarity analyses”; the former sought to examine the test-retest reliability across two sessions using the same imaging sequence, whereas the latter sought to investigate the similarity between two different imaging sequences within the same study visit. Intraclass correlation coefficients (ICCs) were determined using the ICC function in the *psych* package. ICCs (3,1) (i.e. consistency) for which p<0.1 were interpreted according to previously established criteria (Cicchetti, 1994):

- “excellent”: 1.00-0.75
- “good”: 0.74-0.60
- “fair”: 0.59-0.40
- “poor”: 0.39-0.00

ICCs (2,1) (i.e. absolute agreement) are reported in Tables and the Supplementary Material but were not used for interpretation. Also, absolute percentage difference (APD) was calculated according to the following formula: [(timepoint 1 value - timepoint 2 value) / ((timepoint 1 value + timepoint 2 value) / 2)] × 100. Finally, linear mixed-effects models from the *lmerTest* package were used. Here, a Bonferroni correction for multiple comparisons was employed; thus, a significance level of p<0.00081 (0.05 / 62 ROIs) was employed for CT linear mixed-effects analyses, whereas a significance level of p<0.005 (0.05 / 10) was employed for both SV and ^1^H-MRS linear mixed-effects analyses. More detail regarding each study protocol is provided below.

#### 2.10.1. Test-Retest Reliability Analyses

For the Prisma Upgrade protocol, ICCs and APDs were calculated for the following pairs: visit 1 and visit 2 (i.e. Trio test-retest reliability, henceforth “Trio-Trio”), visit 2 and visit 3 (i.e. impact of Prisma upgrade, henceforth “Trio-Prisma”), and visit 3 and visit 4 (i.e. Prisma test-retest reliability, henceforth “Prisma-Prisma”). For the vNav protocol, ICCs were calculated between visit 5 and 6 (i.e. Prisma-Prisma). Of note, Trio-Prisma pairs were selected according to the criterion of proximity in time. The breakdown of data within each of the pairs noted above and the duration between acquisitions is in Supplementary Table 1. It deserves mention that MPRAGE reliabilities for CT and SV were recalculated for the first and second visits of the vNav protocol to serve as comparison for vNav reliabilities, using the same participant pool to do so. In addition, linear mixed-effects models were used to test differences between acquisitions in the pairs noted above, with visit as a fixed effect and subject as a random effect.

#### 2.10.2. Similarity Analyses

As part of the primary aim of the vNav protocol, similarity analyses compared standard structural sequences with vNav sequences within the same MRI session. Thus, similarity analyses tested the following pair: MPRAGE and vNav-MPRAGE. The breakdown of data within each pair is further displayed in Supplementary Table 1. For each participant, pairs were preferentially selected from the first visit of the vNav protocol. As above, ICCs and APDs were calculated for each pair, and linear mixed-effects models were used to test differences between acquisitions, with acquisition as a fixed effect and subject as a random effect.

#### 2.10.3. Signal-to-Noise Analyses

Using only the data from Trio-Prisma pairs noted above, a linear mixed-effects model was used to compare SNR between Trio and Prisma acquisitions, with scanner as a fixed effect and subject as a random effect. In addition, linear mixed-effects models were used to test the relationship between structural measures (i.e. ROI CT and SV) and SNR, with SNR as a fixed effect and subject as a random effect.

## 3. Results

### 3.1. Prisma Upgrade Protocol

#### 3.1.1. Cortical Thickness

ICCs, APDs, and linear mixed-effects analyses results reflecting test-retest reliability for CT measures estimated from CIVET and FreeSurfer across Trio-Trio, Trio-Prisma, and Prisma-Prisma investigations are shown in Table 1 (CIVET and Trio-Prisma) and Supplementary Tables 2 - 6. In the aforementioned analyses, reliability was found to be “excellent” or “good”, with the exception of those detailed below. For Trio-Trio investigations, reliability was “fair” within the right lateral orbitofrontal (CIVET) ROI. For Trio-Prisma investigations, reliability was “fair” within right lingual gyrus (CIVET), left transverse temporal (FreeSurfer), right rostral anterior cingulate (FreeSurfer), and right inferior occipital (FreeSurfer) ROIs, while reliability was “poor” within the right isthmus cingulate gyrus (CIVET) ROI. ICC results using CIVET and FreeSurfer are visualized in Figure 2 and Supplementary Figure 1, respectively.

**Figure 2.**
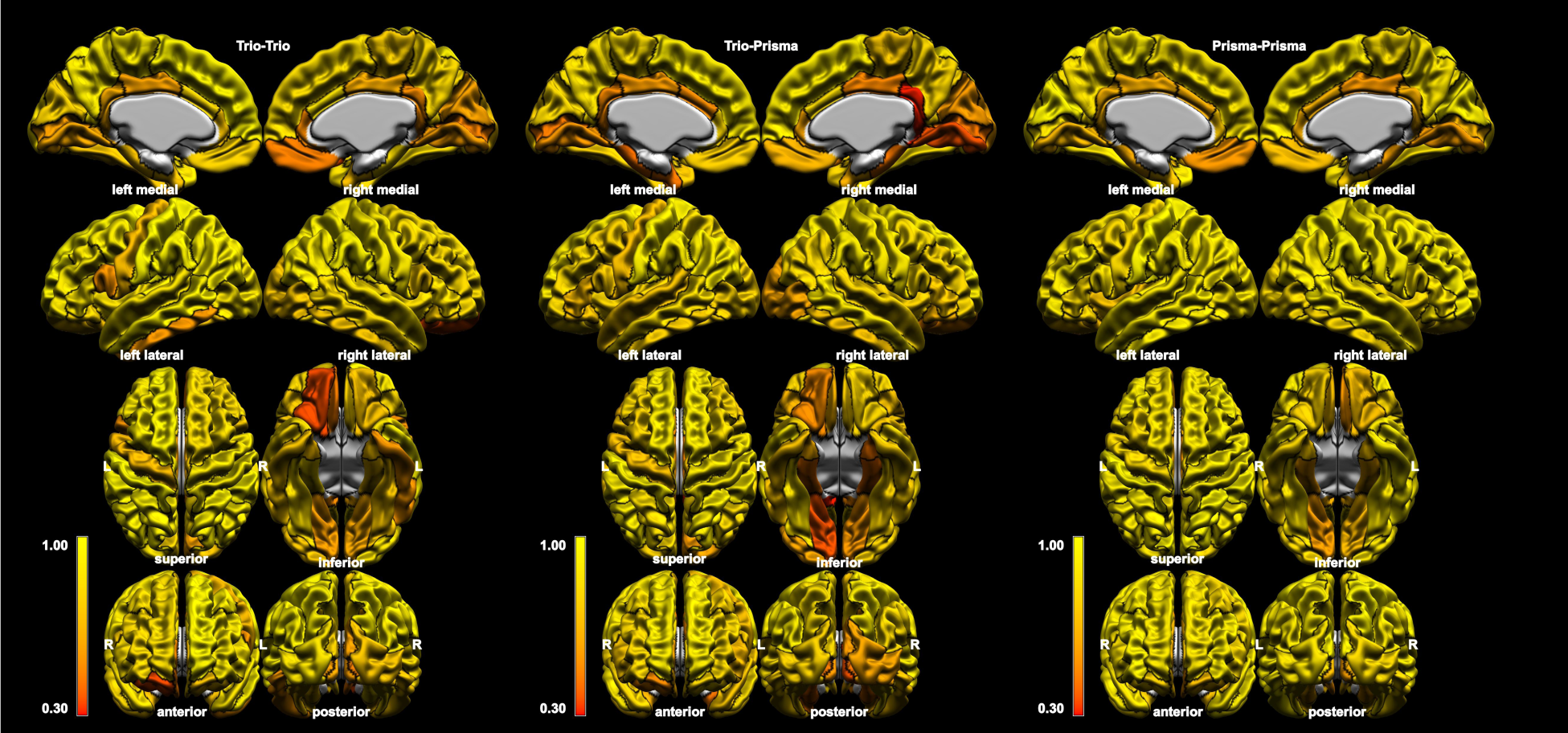
Intraclass correlation coefficients of cortical thickness estimated by the CIVET pipeline within regions of interest defined by the Desikan-Killiany-Tourville atlas (black boundaries). Results are shown for pairs consisting of two T1-weighted images from a Trio machine (left), Prisma machine (right), and one of each (middle).

**Table 1.** Trio-Prisma reliability measures for cortical thickness estimated by the CIVET pipeline.

| ROI | CIVET |  |  |  |  |  |  |  |  |  |  |  |  |  |  |  |
| --- | --- | --- | --- | --- | --- | --- | --- | --- | --- | --- | --- | --- | --- | --- | --- | --- |
|  | Left Hemisphere |  |  |  |  |  |  |  | Right Hemisphere |  |  |  |  |  |  |  |
|  | Timepoint 1<br>Mean (SD) | Timepoint 2<br>Mean (SD) | APD | ICC (2,1) | ICC (3,1) | p-value (ICC) | t-value (lmer) | p-value (lmer) | Timepoint 1<br>Mean (SD) | Timepoint 2<br>Mean (SD) | APD | ICC (2,1) | ICC (3,1) | p-value (ICC) | t-value (lmer) | p-value (lmer) |
| Caudal Anterior Cingulate | 2.82 (0.11) | 2.92 (0.13) | 3.80 | 0.53 | 0.70 | 0.003 | 3.90 | 0.002 | 2.71 (0.14) | 2.75 (0.13) | 1.89 | 0.89 | 0.92 | <0.001 | 2.59 | 0.023 |
| Caudal Middle Frontal | 3.06 (0.07) | 3.08 (0.07) | 0.91 | 0.87 | 0.89 | <0.001 | 1.87 | 0.087 | 3.01 (0.10) | 3.03 (0.11) | 1.24 | 0.90 | 0.92 | <0.001 | 2.10 | 0.057 |
| Cuneus | 2.53 (0.11) | 2.52 (0.12) | 1.60 | 0.90 | 0.91 | <0.001 | -1.11 | 0.287 | 2.61 (0.10) | 2.59 (0.11) | 2.64 | 0.74 | 0.74 | 0.001 | -1.12 | 0.284 |
| Entorhinal Cortex | 3.91 (0.26) | 3.91 (0.24) | 3.78 | 0.64 | 0.64 | 0.007 | 0.14 | 0.892 | 4.03 (0.18) | 4.00 (0.22) | 2.70 | 0.80 | 0.80 | <0.001 | -0.93 | 0.372 |
| Fusiform Gyrus | 3.15 (0.10) | 3.17 (0.11) | 1.22 | 0.89 | 0.89 | <0.001 | 1.31 | 0.213 | 3.17 (0.14) | 3.20 (0.13) | 1.47 | 0.91 | 0.93 | <0.001 | 2.41 | 0.033 |
| Inferior Occipital Cortex | 2.70 (0.07) | 2.67 (0.09) | 1.56 | 0.82 | 0.89 | <0.001 | -3.19 | 0.008 | 2.81 (0.07) | 2.82 (0.08) | 1.41 | 0.77 | 0.77 | <0.001 | 0.67 | 0.515 |
| Inferior Parietal | 2.96 (0.08) | 2.95 (0.10) | 0.70 | 0.96 | 0.96 | <0.001 | -1.66 | 0.123 | 3.02 (0.09) | 3.06 (0.11) | 1.72 | 0.83 | 0.91 | <0.001 | 4.02 | 0.002 |
| Inferior Temporal | 3.18 (0.12) | 3.23 (0.10) | 1.83 | 0.78 | 0.88 | <0.001 | 3.73 | 0.003 | 3.21 (0.12) | 3.31 (0.13) | 3.15 | 0.64 | 0.85 | <0.001 | 5.32 | <0.001* |
| Insula | 3.61 (0.19) | 3.65 (0.18) | 2.05 | 0.89 | 0.90 | <0.001 | 1.64 | 0.128 | 3.55 (0.16) | 3.66 (0.19) | 3.18 | 0.74 | 0.89 | <0.001 | 4.95 | <0.001* |
| Isthmus Cingulate Gyrus | 3.13 (0.19) | 3.19 (0.19) | 3.99 | 0.67 | 0.69 | 0.003 | 1.36 | 0.199 | 3.08 (0.14) | 3.21 (0.17) | 6.41 | 0.29 | 0.39 | 0.087 | 2.77 | 0.017 |
| Lateral Frontal Opercularis | 3.04 (0.07) | 3.06 (0.07) | 1.18 | 0.86 | 0.87 | <0.001 | 1.64 | 0.128 | 3.03 (0.08) | 3.05 (0.07) | 1.02 | 0.87 | 0.91 | <0.001 | 2.92 | 0.013 |
| Lateral Frontal Orbitalis | 3.10 (0.10) | 3.14 (0.09) | 1.78 | 0.78 | 0.85 | <0.001 | 2.84 | 0.015 | 3.12 (0.10) | 3.20 (0.09) | 2.75 | 0.61 | 0.81 | <0.001 | 4.83 | <0.001* |
| Lateral Frontal Triangularis | 2.96 (0.08) | 3.00 (0.10) | 1.56 | 0.77 | 0.83 | <0.001 | 2.72 | 0.019 | 2.93 (0.11) | 2.95 (0.10) | 1.50 | 0.85 | 0.86 | <0.001 | 1.34 | 0.204 |
| Lateral Orbitofrontal | 2.92 (0.09) | 3.00 (0.11) | 2.75 | 0.66 | 0.90 | <0.001 | 6.85 | <0.001* | 2.83 (0.07) | 2.95 (0.10) | 3.99 | 0.39 | 0.70 | 0.003 | 5.98 | <0.001* |
| Lingual Gyrus | 2.59 (0.10) | 2.64 (0.10) | 2.38 | 0.68 | 0.73 | 0.002 | 2.02 | 0.066 | 2.67 (0.09) | 2.71 (0.09) | 2.55 | 0.50 | 0.54 | 0.024 | 1.76 | 0.104 |
| Medial Orbitofrontal | 2.62 (0.10) | 2.73 (0.11) | 4.07 | 0.56 | 0.88 | <0.001 | 7.80 | <0.001* | 2.64 (0.11) | 2.73 (0.09) | 3.50 | 0.58 | 0.81 | <0.001 | 5.30 | <0.001* |
| Middle Temporal | 3.27 (0.08) | 3.28 (0.11) | 0.90 | 0.92 | 0.92 | <0.001 | 0.60 | 0.559 | 3.30 (0.11) | 3.36 (0.12) | 1.87 | 0.83 | 0.95 | <0.001 | 5.87 | <0.001* |
| Paracentral Gyrus | 2.97 (0.11) | 2.97 (0.12) | 1.11 | 0.92 | 0.92 | <0.001 | 0.15 | 0.886 | 3.01 (0.09) | 3.00 (0.10) | 1.36 | 0.86 | 0.86 | <0.001 | -0.55 | 0.592 |
| Parahippocampal | 2.74 (0.14) | 2.79 (0.14) | 3.77 | 0.60 | 0.62 | 0.010 | 1.41 | 0.184 | 2.75 (0.13) | 2.77 (0.13) | 3.18 | 0.64 | 0.64 | 0.007 | 0.81 | 0.435 |
| Pericalcarine | 2.45 (0.10) | 2.42 (0.09) | 3.00 | 0.62 | 0.65 | 0.006 | -1.53 | 0.152 | 2.50 (0.09) | 2.50 (0.08) | 2.29 | 0.64 | 0.64 | 0.007 | -0.07 | 0.943 |
| Postcentral Gyrus | 2.68 (0.12) | 2.65 (0.13) | 1.38 | 0.94 | 0.97 | <0.001 | -3.99 | 0.002 | 2.69 (0.10) | 2.68 (0.09) | 0.97 | 0.94 | 0.95 | <0.001 | -1.68 | 0.119 |
| Posterior Cingulate | 2.99 (0.10) | 2.97 (0.13) | 2.05 | 0.74 | 0.74 | 0.001 | -0.49 | 0.633 | 2.93 (0.07) | 2.94 (0.08) | 1.74 | 0.68 | 0.68 | 0.003 | 0.54 | 0.599 |
| Precentral Gyrus | 3.10 (0.06) | 3.08 (0.09) | 1.22 | 0.82 | 0.84 | <0.001 | -1.83 | 0.092 | 3.05 (0.10) | 3.06 (0.11) | 0.76 | 0.95 | 0.95 | <0.001 | 0.91 | 0.381 |
| Precuneus | 2.93 (0.08) | 2.91 (0.08) | 0.93 | 0.91 | 0.93 | <0.001 | -2.16 | 0.052 | 3.00 (0.08) | 3.01 (0.10) | 1.12 | 0.90 | 0.90 | <0.001 | 0.88 | 0.395 |
| Rostral Anterior Cingulate | 3.03 (0.14) | 3.11 (0.14) | 2.40 | 0.80 | 0.90 | <0.001 | 4.28 | 0.001 | 2.86 (0.12) | 2.94 (0.11) | 3.35 | 0.63 | 0.77 | <0.001 | 3.76 | 0.003 |
| Rostral Middle Frontal | 2.97 (0.12) | 3.01 (0.12) | 1.68 | 0.86 | 0.91 | <0.001 | 2.98 | 0.011 | 2.86 (0.11) | 2.91 (0.10) | 1.92 | 0.82 | 0.90 | <0.001 | 3.59 | 0.004 |
| Superior Frontal Gyrus | 3.11 (0.12) | 3.13 (0.12) | 0.97 | 0.95 | 0.96 | <0.001 | 2.14 | 0.054 | 3.05 (0.11) | 3.07 (0.10) | 0.96 | 0.95 | 0.96 | <0.001 | 2.28 | 0.042 |
| Superior Parietal | 2.73 (0.07) | 2.70 (0.08) | 1.41 | 0.84 | 0.92 | <0.001 | -4.35 | <0.001 | 2.74 (0.09) | 2.73 (0.10) | 0.84 | 0.96 | 0.96 | <0.001 | -1.17 | 0.266 |
| Superior Temporal | 3.28 (0.08) | 3.32 (0.11) | 1.54 | 0.81 | 0.86 | <0.001 | 2.70 | 0.019 | 3.30 (0.09) | 3.35 (0.10) | 1.75 | 0.78 | 0.90 | <0.001 | 4.64 | <0.001* |
| Supramarginal Gyrus | 3.03 (0.12) | 3.03 (0.14) | 0.91 | 0.96 | 0.96 | <0.001 | -0.14 | 0.887 | 3.04 (0.13) | 3.05 (0.14) | 0.99 | 0.96 | 0.96 | <0.001 | 0.62 | 0.548 |
| Transverse Temporal | 2.99 (0.12) | 3.04 (0.13) | 1.98 | 0.83 | 0.89 | <0.001 | 3.11 | 0.009 | 3.02 (0.13) | 3.07 (0.18) | 2.46 | 0.85 | 0.89 | <0.001 | 2.49 | 0.028 |

APD, absolute percentage difference; ICC, intraclass correlation coefficient; lmer, linear mixed-effects model; ROI, region of interest; SD, standard deviation. Note: positive t-values represent cortical thickness increases. Asterisk indicates significance after Bonferroni correction.

Results of linear mixed-effects analyses demonstrated differences between Trio and Prisma estimates. Using CIVET, a significant effect of the upgrade was found for the following ROIs: bilateral lateral orbitofrontal, bilateral medial orbitofrontal, right inferior temporal, right insula, right lateral frontal orbitalis, right middle temporal, and right superior temporal (Figure 3). Using FreeSurfer, a significant effect of the upgrade was found for the following ROIs: bilateral pericalcarine, left caudal anterior cingulate, left superior parietal, and right insula (Supplementary Figure 2).

**Figure 3.**
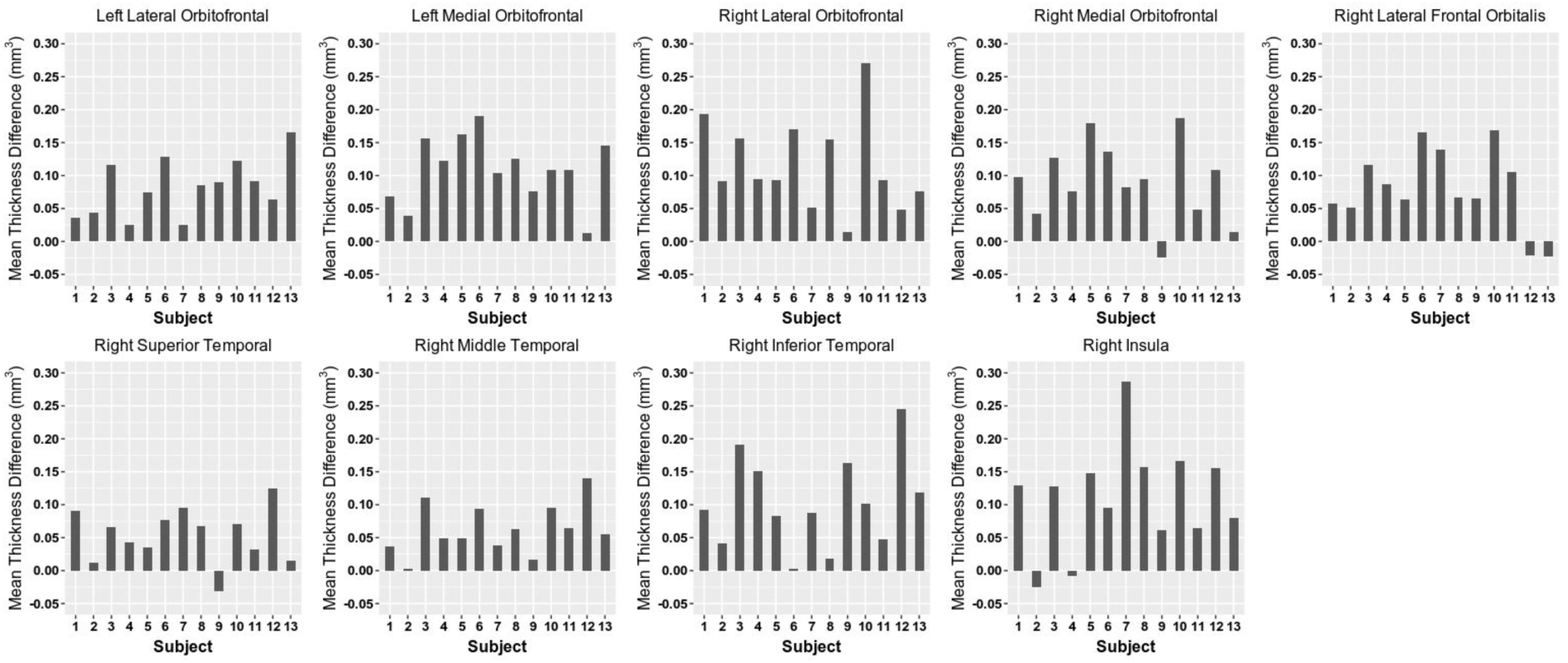
Cortical thickness estimated by the CIVET pipeline within regions of interest that achieved statistical significance in linear mixed-effects analyses. Mean thickness difference represents Prisma estimate minus Trio estimate.

#### 3.1.2. Structure Volume

ICCs, APDs, and linear mixed-effects analyses results reflecting test-retest reliability for SV measures estimated from MAGeT-Brain and FreeSurfer across Trio-Trio, Trio-Prisma, and Prisma-Prisma investigations are shown in Table 2 (MAGeT-Brain and Trio-Prisma) and Supplementary Tables 7 - 11. All MAGeT-Brain analyses rendered “excellent” reliability. Using FreeSurfer, reliability was found to be “excellent” or “good”, with the exception of those detailed below. Reliability with FreeSurfer across Trio-Prisma investigations was “fair” within the left amygdala and right thalamus, and did not reach statistical significance within the bilateral globus pallidus, left hippocampus, and left thalamus. Further, reliability with FreeSurfer across Prisma-Prisma investigations was “fair” within the left globus pallidus and “poor” within the right globus pallidus.

**Table 2.** Trio-Prisma reliability measures for structure volume estimated by the MAGeT-Brain pipeline.

|  | MAGeT-Brain |  |  |  |  |  |  |  |  |  |  |  |  |  |  |  |
| --- | --- | --- | --- | --- | --- | --- | --- | --- | --- | --- | --- | --- | --- | --- | --- | --- |
|  | Left Hemisphere |  |  |  |  |  |  |  | Right Hemisphere |  |  |  |  |  |  |  |
| ROI | Timepoint 1 Mean (SD) | Timepoint 2 Mean (SD) | APD | ICC (2,1) | ICC (3,1) | p-value (ICC) | t-value (lmer) | p-value (lmer) | Timepoint 1 Mean (SD) | Timepoint 2 Mean (SD) | APD | ICC (2,1) | ICC (3,1) | p-value (ICC) | t-value (lmer) | p-value (lmer) |
| Amygdala | 1245.85 (129.13) | 1218.00 (134.02) | 2.53 | 0.96 | 0.98 | <0.001 | -4.05 | 0.002* | 1191.00 (120.95) | 1182.69 (117.60) | 1.62 | 0.98 | 0.98 | <0.001 | -1.36 | 0.199 |
| Hippocampus | 2423.23 (243.48) | 2356.62 (202.27) | 2.67 | 0.92 | 0.96 | <0.001 | -3.61 | 0.004* | 2491.85 (277.44) | 2450.46 (223.06) | 2.67 | 0.94 | 0.95 | <0.001 | -1.94 | 0.077 |
| Globus Pallidus | 1616.85 (138.56) | 1569.39 (140.92) | 3.01 | 0.93 | 0.98 | <0.001 | -5.70 | <0.001* | 1440.23 (124.21) | 1403.62 (138.54) | 3.11 | 0.92 | 0.95 | <0.001 | -3.21 | 0.008 |
| Striatum | 9811.00 (1031.29) | 9861.69 (1054.92) | 0.65 | 1.00 | 1.00 | <0.001 | 2.45 | 0.030 | 10019.46 (1000.60) | 10003.62 (1004.25) | 0.56 | 1.00 | 1.00 | <0.001 | -0.88 | 0.396 |
| Thalamus | 6186.23 (469.56) | 6414.39 (516.63) | 3.61 | 0.88 | 0.97 | <0.001 | 7.32 | <0.001* | 6148.46 (490.02) | 6381.54 (516.14) | 3.76 | 0.88 | 0.98 | <0.001 | 7.91 | <0.001* |

APD, absolute percentage difference; ICC, intraclass correlation coefficient; lmer, linear mixed-effects model; ROI, region of interest; SD, standard deviation. Note: positive t-values represent structure volume increases. Asterisk indicates significance after Bonferroni correction.

Results of linear mixed-effects analyses demonstrated differences between Trio and Prisma estimates. Using MAGeT-Brain, a significant effect of the upgrade was found for the following ROIs: bilateral thalamus, left amygdala, left globus pallidus, and left hippocampus (Figure 4). Using FreeSurfer, a significant effect of the upgrade was found for the following ROIs: bilateral amygdala, bilateral globus pallidus, and bilateral striatum (Supplementary Figure 3).

**Figure 4.**
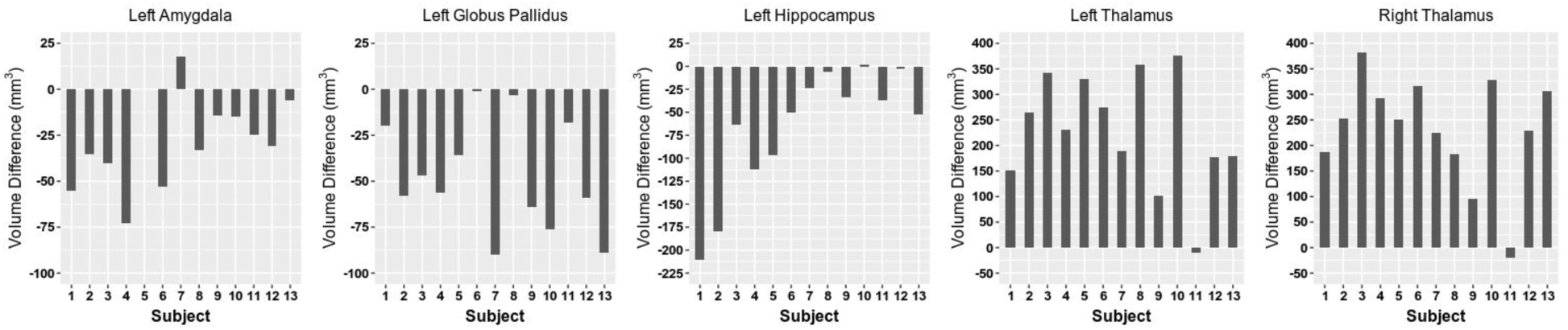
Structure volume estimated by the MAGeT-Brain pipeline within regions of interest that achieved statistical significance in linear mixed-effects analyses. Volume difference represents Prisma estimate minus Trio estimate.

#### 3.1.3. Proton Magnetic Resonance Spectroscopy

ICCs, APDs, and linear mixed-effects analyses results reflecting test-retest reliability for ^1^H-MRS measures across Trio-Trio, Trio-Prisma, and Prisma-Prisma investigations are shown in Table 3 (Trio-Prisma) and Supplementary Tables 12 and 13. All ^1^H-MRS metrics achieved “excellent” or “good” reliability with the exception of those detailed below. For Trio-Trio investigations, reliability was “fair” for Lac and SNR, and did not reach statistical significance for GABA, NAAG, and FWHM. For Trio-Prisma investigations, reliability did not reach statistical significance for GABA, Gln, Lac, NAAG, FWHM, and SNR. For Prisma-Prisma investigations, reliability was “fair” for GABA, NAAG and SNR, and did not reach statistical significance for GSH and Lac. Results of linear-mixed effects analyses were not statistically significant.

**Table 3.**
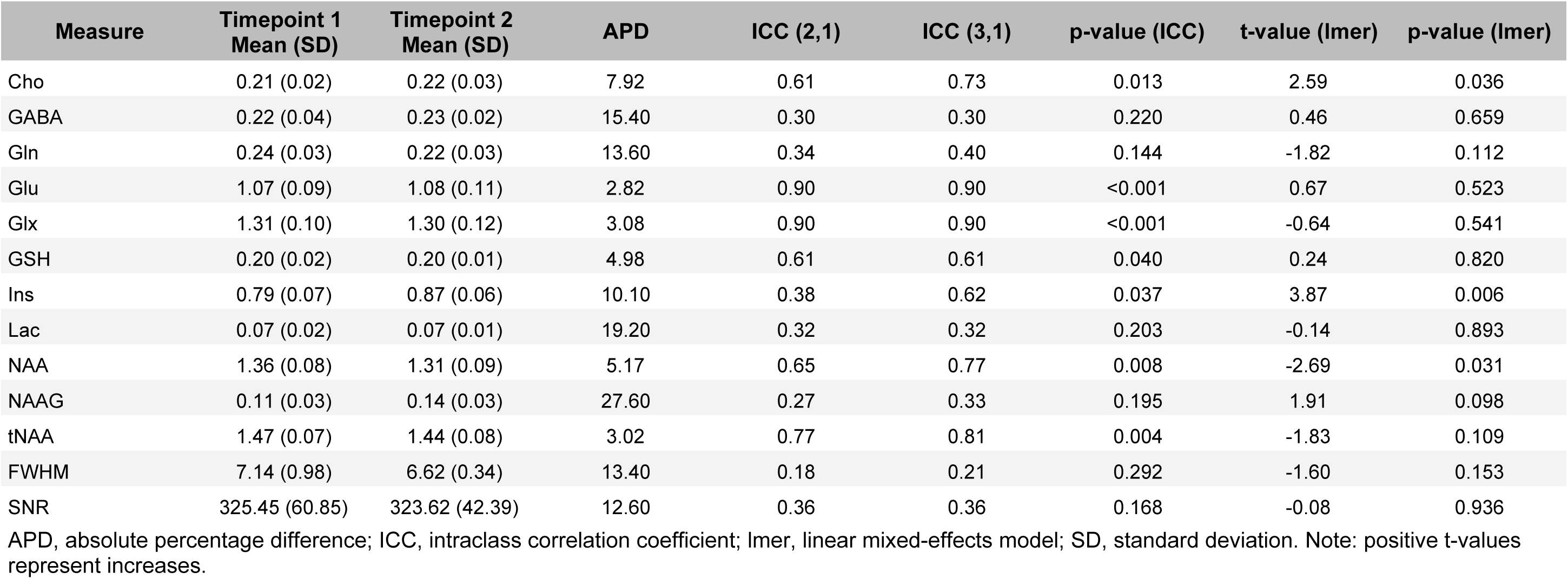
Trio-Prisma reliability measures for 1H-MRS indices.

| Measure | Timepoint 1<br>Mean (SD) | Timepoint 2<br>Mean (SD) | APD | ICC (2,1) | ICC (3,1) | p-value (ICC) | t-value (lmer) | p-value (lmer) |
| --- | --- | --- | --- | --- | --- | --- | --- | --- |
| Cho | 0.21 (0.02) | 0.22 (0.03) | 7.92 | 0.61 | 0.73 | 0.013 | 2.59 | 0.036 |
| GABA | 0.22 (0.04) | 0.23 (0.02) | 15.40 | 0.30 | 0.30 | 0.220 | 0.46 | 0.659 |
| Gln | 0.24 (0.03) | 0.22 (0.03) | 13.60 | 0.34 | 0.40 | 0.144 | -1.82 | 0.112 |
| Glu | 1.07 (0.09) | 1.08 (0.11) | 2.82 | 0.90 | 0.90 | <0.001 | 0.67 | 0.523 |
| Glx | 1.31 (0.10) | 1.30 (0.12) | 3.08 | 0.90 | 0.90 | <0.001 | -0.64 | 0.541 |
| GSH | 0.20 (0.02) | 0.20 (0.01) | 4.98 | 0.61 | 0.61 | 0.040 | 0.24 | 0.820 |
| Ins | 0.79 (0.07) | 0.87 (0.06) | 10.10 | 0.38 | 0.62 | 0.037 | 3.87 | 0.006 |
| Lac | 0.07 (0.02) | 0.07 (0.01) | 19.20 | 0.32 | 0.32 | 0.203 | -0.14 | 0.893 |
| NAA | 1.36 (0.08) | 1.31 (0.09) | 5.17 | 0.65 | 0.77 | 0.008 | -2.69 | 0.031 |
| NAAG | 0.11 (0.03) | 0.14 (0.03) | 27.60 | 0.27 | 0.33 | 0.195 | 1.91 | 0.098 |
| tNAA | 1.47 (0.07) | 1.44 (0.08) | 3.02 | 0.77 | 0.81 | 0.004 | -1.83 | 0.109 |
| FWHM | 7.14 (0.98) | 6.62 (0.34) | 13.40 | 0.18 | 0.21 | 0.292 | -1.60 | 0.153 |
| SNR | 325.45 (60.85) | 323.62 (42.39) | 12.60 | 0.36 | 0.36 | 0.168 | -0.08 | 0.936 |

APD, absolute percentage difference; ICC, intraclass correlation coefficient; lmer, linear mixed-effects model; SD, standard deviation. Note: positive t-values represent increases.

### 3.2. vNav Protocol

#### 3.2.1. Cortical Thickness

##### 3.2.1.1. Test-retest reliability

ICCs, APDs, and linear mixed-effects analyses results reflecting test-retest reliability for CT measures estimated from CIVET and FreeSurfer in MPRAGE and vNav-MPRAGE sequences are shown in Supplementary Tables 14 - 17. Using CIVET, reliability was found to be “excellent” or “good”, with the exception of those detailed below. For MPRAGE investigations, reliability was “fair” within the left lateral orbitofrontal ROI. For vNav-MPRAGE investigations, reliability was “fair” within the left lateral frontal orbitalis, right isthmus cingulate, right pericalcarine, and right posterior cingulate ROIs. FreeSurfer analyses rendered “excellent” or “good” reliability, with the exception of the right rostral anterior cingulate ROI in vNav-MPRAGE sequences, which was found to be “fair”. ICC results using CIVET and FreeSurfer are illustrated in Figure 5 and Supplementary Figure 4, respectively. Results of linear-mixed effects analyses were not statistically significant.

**Figure 5.**
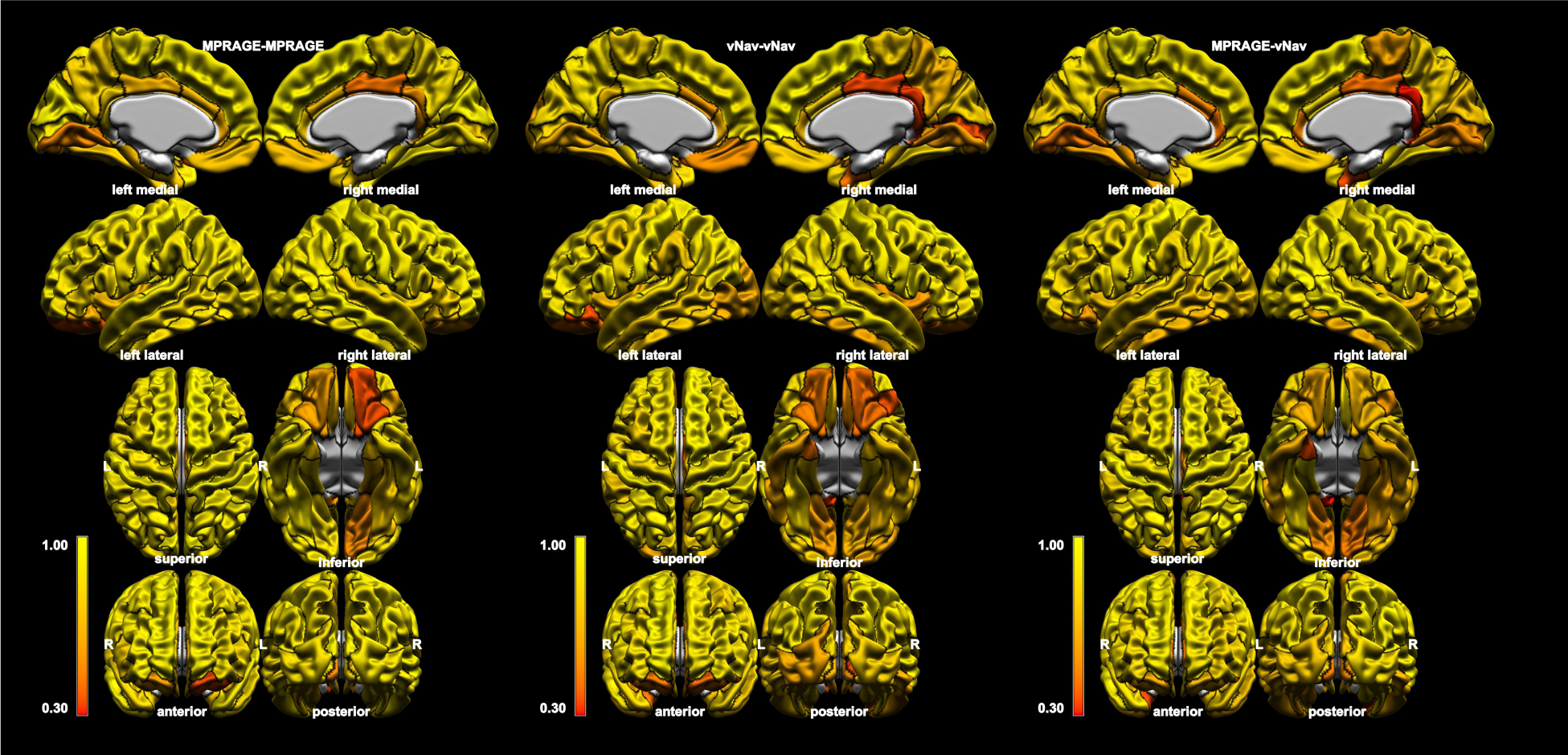
Intraclass correlation coefficients of cortical thickness estimated by the CIVET pipeline within regions of interest defined by the Desikan-Killiany-Tourville atlas (black boundaries). Results are shown for pairs consisting of two T1-weighted images (left), two volumetric navigator T1-weighted images (middle), and one of each (right).

##### 3.2.1.2. Similarity analyses

ICCs, APDs, and linear mixed-effects analyses results reflecting similarity of CT measures acquired from MPRAGE and vNav-MPRAGE sequences, as estimated by CIVET and FreeSurfer, are shown in Table 4 and Supplementary Table 18, respectively. In the aforementioned analyses, reliability was found to be “excellent” or “good”, with the exception of those detailed below. Reliability was “fair” within the right entorhinal cortex (CIVET), “poor” within the right isthmus cingulate (CIVET), and did not reach statistical significance within the right entorhinal cortex (FreeSurfer). ICC results using CIVET and FreeSurfer are illustrated in Figure 5 and Supplementary Figure 4, respectively.

**Table 4.** MPRAGE and vNav-MPRAGE similarity measures for cortical thickness estimated by the CIVET pipeline.

| ROI | CIVET |  |  |  |  |  |  |  |  |  |  |  |  |  |  |  |
| --- | --- | --- | --- | --- | --- | --- | --- | --- | --- | --- | --- | --- | --- | --- | --- | --- |
|  | Left Hemisphere |  |  |  |  |  |  |  | Right Hemisphere |  |  |  |  |  |  |  |
|  | MPRAGE Mean (SD) | vNav-MPRAGE Mean (SD) | APD | ICC (2,1) | ICC (3,1) | p-value (ICC) | t-value (lmer) | p-value (lmer) | MPRAGE Mean (SD) | vNav-MPRAGE Mean (SD) | APD | ICC (2,1) | ICC (3,1) | p-value (ICC) | t-value (lmer) | p-value (lmer) |
| Caudal Anterior Cingulate | 2.86 (0.14) | 2.80 (0.11) | 3.42 | 0.62 | 0.69 | 0.003 | -2.37 | 0.036 | 2.78 (0.13) | 2.74 (0.14) | 2.06 | 0.89 | 0.93 | <0.001 | -2.87 | 0.014 |
| Caudal Middle Frontal | 3.09 (0.07) | 3.06 (0.07) | 0.93 | 0.89 | 0.92 | <0.001 | -2.60 | 0.023 | 3.07 (0.09) | 3.05 (0.08) | 0.88 | 0.92 | 0.95 | <0.001 | -3.06 | 0.010 |
| Cuneus | 2.55 (0.10) | 2.55 (0.11) | 1.52 | 0.90 | 0.90 | <0.001 | 0.02 | 0.984 | 2.59 (0.12) | 2.58 (0.10) | 1.79 | 0.86 | 0.86 | <0.001 | -0.95 | 0.362 |
| Entorhinal Cortex | 4.00 (0.29) | 3.83 (0.27) | 4.57 | 0.75 | 0.88 | <0.001 | -4.35 | <0.001 | 4.07 (0.22) | 3.98 (0.19) | 4.13 | 0.50 | 0.53 | 0.025 | -1.72 | 0.112 |
| Fusiform Gyrus | 3.14 (0.09) | 3.11 (0.08) | 1.55 | 0.76 | 0.78 | <0.001 | -1.57 | 0.143 | 3.19 (0.11) | 3.14 (0.10) | 1.67 | 0.86 | 0.93 | <0.001 | -3.91 | 0.002 |
| Inferior Occipital Cortex | 2.69 (0.07) | 2.68 (0.07) | 1.08 | 0.87 | 0.87 | <0.001 | -0.97 | 0.351 | 2.81 (0.09) | 2.78 (0.08) | 1.31 | 0.89 | 0.93 | <0.001 | -2.92 | 0.013 |
| Inferior Parietal | 2.95 (0.10) | 2.96 (0.09) | 0.93 | 0.94 | 0.94 | <0.001 | 0.90 | 0.383 | 3.07 (0.10) | 3.02 (0.09) | 1.58 | 0.86 | 0.97 | <0.001 | -8.39 | <0.001* |
| Inferior Temporal | 3.19 (0.12) | 3.20 (0.10) | 1.47 | 0.82 | 0.82 | <0.001 | 0.54 | 0.598 | 3.28 (0.12) | 3.23 (0.08) | 2.03 | 0.74 | 0.81 | <0.001 | -2.72 | 0.019 |
| Insula | 3.67 (0.15) | 3.61 (0.15) | 2.44 | 0.78 | 0.85 | <0.001 | -3.00 | 0.011 | 3.67 (0.17) | 3.60 (0.13) | 2.29 | 0.78 | 0.84 | <0.001 | -2.56 | 0.025 |
| Isthmus Cingulate Gyrus | 3.16 (0.18) | 3.04 (0.23) | 3.90 | 0.73 | 0.83 | <0.001 | -3.29 | 0.006 | 3.10 (0.13) | 2.97 (0.17) | 5.73 | 0.27 | 0.37 | 0.097 | -2.92 | 0.013 |
| Lateral Frontal Opercularis | 3.07 (0.10) | 3.06 (0.08) | 0.96 | 0.93 | 0.93 | <0.001 | -0.64 | 0.534 | 3.06 (0.11) | 3.04 (0.10) | 1.03 | 0.94 | 0.94 | <0.001 | -1.26 | 0.232 |
| Lateral Frontal Orbitalis | 3.15 (0.08) | 3.13 (0.09) | 1.45 | 0.74 | 0.74 | 0.001 | -1.22 | 0.246 | 3.16 (0.14) | 3.12 (0.12) | 2.02 | 0.83 | 0.85 | <0.001 | -1.96 | 0.073 |
| Lateral Frontal Triangularis | 2.97 (0.08) | 2.97 (0.08) | 1.05 | 0.91 | 0.91 | <0.001 | -0.29 | 0.776 | 2.97 (0.10) | 2.96 (0.09) | 0.97 | 0.94 | 0.94 | <0.001 | -0.51 | 0.621 |
| Lateral Orbitofrontal | 3.02 (0.07) | 2.98 (0.07) | 1.69 | 0.71 | 0.83 | <0.001 | -3.77 | 0.003 | 2.95 (0.08) | 2.92 (0.09) | 1.56 | 0.77 | 0.80 | <0.001 | -1.98 | 0.072 |
| Lingual Gyrus | 2.63 (0.09) | 2.59 (0.08) | 2.50 | 0.60 | 0.66 | 0.005 | -2.21 | 0.047 | 2.66 (0.09) | 2.61 (0.06) | 2.75 | 0.59 | 0.71 | 0.002 | -3.14 | 0.008 |
| Medial Orbitofrontal | 2.73 (0.12) | 2.78 (0.11) | 1.87 | 0.86 | 0.94 | <0.001 | 4.51 | <0.001* | 2.72 (0.11) | 2.77 (0.11) | 2.55 | 0.78 | 0.88 | <0.001 | 3.80 | 0.003 |
| Middle Temporal | 3.28 (0.11) | 3.26 (0.10) | 1.56 | 0.85 | 0.86 | <0.001 | -1.34 | 0.206 | 3.37 (0.12) | 3.32 (0.11) | 1.61 | 0.88 | 0.97 | <0.001 | -6.16 | <0.001* |
| Paracentral Gyrus | 2.96 (0.13) | 2.96 (0.14) | 0.73 | 0.98 | 0.98 | <0.001 | -0.77 | 0.454 | 3.00 (0.10) | 2.93 (0.10) | 2.47 | 0.63 | 0.77 | <0.001 | -3.65 | 0.003 |
| Parahippocampal | 2.83 (0.11) | 2.72 (0.14) | 4.39 | 0.56 | 0.77 | <0.001 | -4.75 | <0.001* | 2.83 (0.15) | 2.71 (0.12) | 4.13 | 0.70 | 0.95 | <0.001 | -10.09 | <0.001* |
| Pericalcarine | 2.40 (0.12) | 2.42 (0.11) | 1.73 | 0.90 | 0.90 | <0.001 | 1.26 | 0.231 | 2.49 (0.10) | 2.48 (0.08) | 1.92 | 0.73 | 0.73 | 0.001 | -0.61 | 0.556 |
| Postcentral Gyrus | 2.69 (0.11) | 2.71 (0.11) | 1.10 | 0.95 | 0.98 | <0.001 | 4.58 | <0.001* | 2.69 (0.08) | 2.71 (0.09) | 0.98 | 0.93 | 0.94 | <0.001 | 1.68 | 0.118 |
| Posterior Cingulate | 2.96 (0.12) | 2.94 (0.11) | 1.38 | 0.93 | 0.94 | <0.001 | -1.82 | 0.093 | 2.96 (0.07) | 2.89 (0.06) | 2.42 | 0.41 | 0.63 | 0.008 | -4.42 | <0.001 |
| Precentral Gyrus | 3.08 (0.08) | 3.05 (0.07) | 0.87 | 0.90 | 0.94 | <0.001 | -3.16 | 0.008 | 3.09 (0.08) | 3.05 (0.08) | 1.13 | 0.89 | 0.97 | <0.001 | -6.56 | <0.001* |
| Precuneus | 2.93 (0.07) | 2.92 (0.07) | 0.82 | 0.93 | 0.93 | <0.001 | -1.42 | 0.180 | 3.02 (0.07) | 2.96 (0.06) | 1.97 | 0.66 | 0.90 | <0.001 | -6.93 | <0.001* |
| Rostral Anterior Cingulate | 3.06 (0.23) | 3.05 (0.17) | 4.58 | 0.64 | 0.64 | 0.007 | -0.27 | 0.792 | 2.95 (0.15) | 2.91 (0.15) | 3.57 | 0.68 | 0.69 | 0.003 | -1.26 | 0.231 |
| Rostral Middle Frontal | 3.02 (0.10) | 3.01 (0.10) | 1.14 | 0.91 | 0.91 | <0.001 | -0.97 | 0.351 | 2.95 (0.09) | 2.97 (0.08) | 1.00 | 0.89 | 0.92 | <0.001 | 2.42 | 0.032 |
| Superior Frontal Gyrus | 3.16 (0.11) | 3.17 (0.11) | 0.85 | 0.95 | 0.95 | <0.001 | 0.83 | 0.422 | 3.09 (0.12) | 3.12 (0.10) | 0.85 | 0.95 | 0.97 | <0.001 | 3.38 | 0.005 |
| Superior Parietal | 2.72 (0.09) | 2.72 (0.08) | 0.76 | 0.95 | 0.95 | <0.001 | -0.30 | 0.768 | 2.77 (0.08) | 2.76 (0.07) | 0.64 | 0.96 | 0.96 | <0.001 | -2.15 | 0.053 |
| Superior Temporal | 3.30 (0.11) | 3.26 (0.11) | 1.44 | 0.87 | 0.90 | <0.001 | -2.45 | 0.031 | 3.35 (0.10) | 3.30 (0.09) | 1.57 | 0.81 | 0.93 | <0.001 | -5.32 | <0.001* |
| Supramarginal Gyrus | 3.06 (0.11) | 3.04 (0.11) | 1.20 | 0.91 | 0.91 | <0.001 | -0.85 | 0.413 | 3.09 (0.12) | 3.05 (0.10) | 1.16 | 0.92 | 0.96 | <0.001 | -3.76 | 0.003 |
| Transverse Temporal | 3.00 (0.15) | 2.99 (0.13) | 1.59 | 0.92 | 0.92 | <0.001 | -0.29 | 0.775 | 3.06 (0.15) | 3.05 (0.13) | 1.18 | 0.96 | 0.96 | <0.001 | -0.70 | 0.499 |

APD, absolute percentage difference; ICC, intraclass correlation coefficient; lmer, linear mixed-effects model; ROI, region of interest; SD, standard deviation. Note: positive t-values represent cortical thickness increases in vNav-MPRAGE. Asterisk indicates significance after Bonferroni correction.

Results of linear mixed-effects analyses rendered differences between CT estimates from MPRAGE and vNav-MPRAGE sequences. Using CIVET, a significant effect of sequence type was found for the following ROIs: bilateral parahippocampal, left medial orbitofrontal, left postcentral, right inferior parietal, right middle temporal, right precentral, right precuneus, and right superior temporal (Figure 6). Using FreeSurfer, a significant effect of sequence type was found for the following ROIs: bilateral postcentral, left inferior parietal, left superior parietal, left transverse temporal, right inferior occipital, right lingual, right lateral frontal triangularis, right pericalcarine, right rostral middle frontal, and right superior frontal (Supplementary Figure 5).

**Figure 6.**
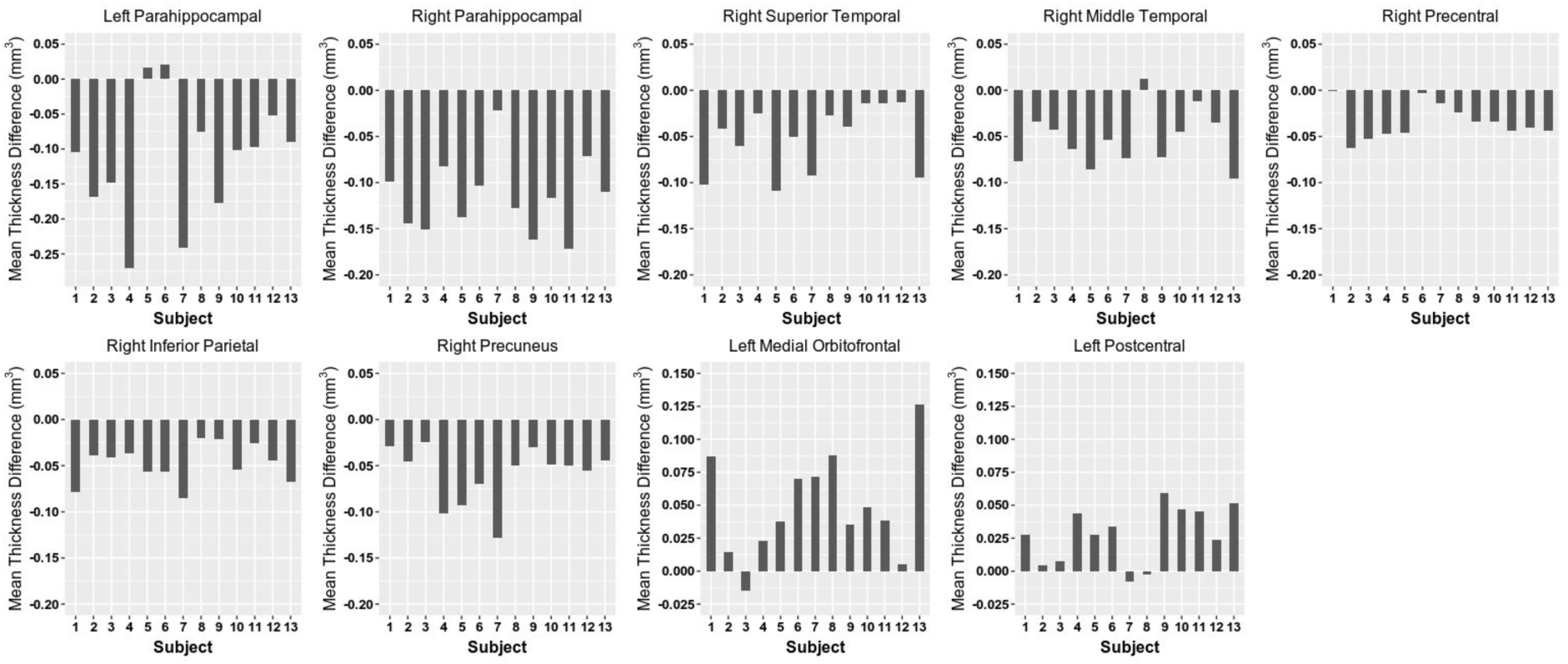
Cortical thickness estimated by the CIVET pipeline within regions of interest that achieved statistical significance in linear mixed-effects analyses. Mean thickness difference represents vNav-MPRAGE estimate minus MPRAGE estimate.

#### 3.2.2. Structure Volume

##### 3.2.2.1. Test-retest reliability

ICCs, APDs, and linear mixed-effects analyses results reflecting test-retest reliability for SV measures estimated from MAGeT-Brain and FreeSurfer in MPRAGE and vNav-MPRAGE sequences are shown in Supplementary Tables 19 - 22. All analyses rendered “excellent” or “good” reliability. Using MAGeT-Brain, results of linear-mixed effects analyses were not statistically significant. Using FreeSurfer, a significant effect of visit was found in the right amygdala for vNav-MPRAGE sequences.

##### 3.2.2.2. Similarity analyses

ICCs, APDs, and linear mixed-effects analyses results reflecting similarity of SV measures acquired from MPRAGE and vNav-MPRAGE sequences, as estimated by MAGeT-Brain and FreeSurfer, are shown in Table 5 and Supplementary Table 23, respectively. All analyses rendered “excellent” reliability. Using MAGeT-Brain, results of linear-mixed effects analyses were not statistically significant. Using FreeSurfer, a significant effect of sequence was found in the left hippocampus and right thalamus (Supplementary Figure 6).

**Table 5.**
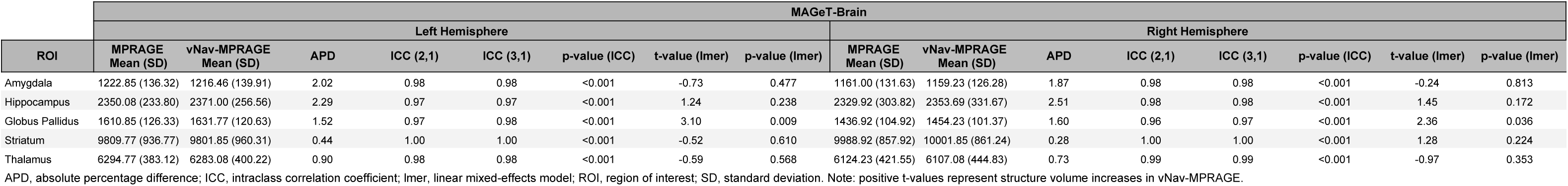
MPRAGE and vNav-MPRAGE similarity measures for structure volume estimated by the MAGeT-Brain pipeline.

| ROI | MAGEt-Brain |  |  |  |  |  |  |  |  |  |  |  |  |  |  |  |
| --- | --- | --- | --- | --- | --- | --- | --- | --- | --- | --- | --- | --- | --- | --- | --- | --- |
|  | Left Hemisphere |  |  |  |  |  |  |  | Right Hemisphere |  |  |  |  |  |  |  |
|  | MPRAGE<br>Mean (SD) | vNav-MPRAGE<br>Mean (SD) | APD | ICC (2,1) | ICC (3,1) | p-value (ICC) | t-value (lmer) | p-value (lmer) | MPRAGE<br>Mean (SD) | vNav-MPRAGE<br>Mean (SD) | APD | ICC (2,1) | ICC (3,1) | p-value (ICC) | t-value (lmer) | p-value (lmer) |
| Amygdala | 1222.85 (136.32) | 1216.46 (139.91) | 2.02 | 0.98 | 0.98 | <0.001 | -0.73 | 0.477 | 1161.00 (131.63) | 1159.23 (126.28) | 1.87 | 0.98 | 0.98 | <0.001 | -0.24 | 0.813 |
| Hippocampus | 2350.08 (233.80) | 2371.00 (256.56) | 2.29 | 0.97 | 0.97 | <0.001 | 1.24 | 0.238 | 2329.92 (303.82) | 2353.69 (331.67) | 2.51 | 0.98 | 0.98 | <0.001 | 1.45 | 0.172 |
| Globus Pallidus | 1610.85 (126.33) | 1631.77 (120.63) | 1.52 | 0.97 | 0.98 | <0.001 | 3.10 | 0.009 | 1436.92 (104.92) | 1454.23 (101.37) | 1.60 | 0.96 | 0.97 | <0.001 | 2.36 | 0.036 |
| Striatum | 9809.77 (936.77) | 9801.85 (960.31) | 0.44 | 1.00 | 1.00 | <0.001 | -0.52 | 0.610 | 9988.92 (857.92) | 10001.85 (861.24) | 0.28 | 1.00 | 1.00 | <0.001 | 1.28 | 0.224 |
| Thalamus | 6294.77 (383.12) | 6283.08 (400.22) | 0.90 | 0.98 | 0.98 | <0.001 | -0.59 | 0.568 | 6124.23 (421.55) | 6107.08 (444.83) | 0.73 | 0.99 | 0.99 | <0.001 | -0.97 | 0.353 |

APD, absolute percentage difference; ICC, intraclass correlation coefficient; lmer, linear mixed-effects model; ROI, region of interest; SD, standard deviation. Note: positive t-values represent structure volume increases in vNav-MPRAGE.

### 3.3. Signal-to-Noise Analyses

A significant increase in SNR was identified following the scanner upgrade (Trio: 20.07 ± 0.66, Prisma: 24.30 ± 1.32, t=13.98, p<0.001) (Figure 7).

**Figure 7.**
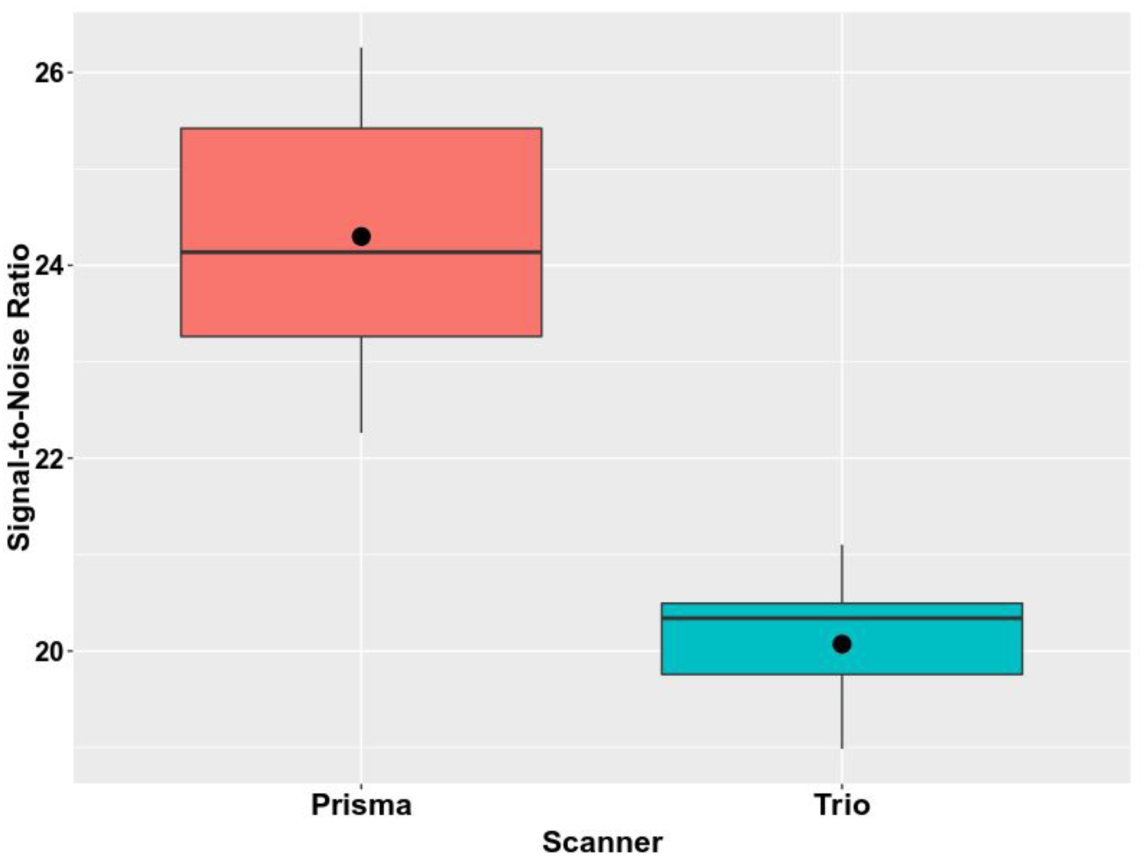
Signal-to-Noise Ratios in Siemens Prisma and Trio scanners. Center circle reflects mean value.

Linear mixed-effects analyses results reflecting the relationship between SNR and CT are shown in Supplementary Table 24. Using CIVET, a significant association between CT and SNR was identified within bilateral lateral orbitofrontal, bilateral medial orbitofrontal, bilateral rostral anterior cingulate, left caudal anterior cingulate, right inferior parietal, right inferior temporal, right insula, right middle temporal, and right superior temporal ROIs. Using FreeSurfer, a significant association between CT and SNR was identified within left caudal anterior cingulate, left superior parietal, and right insula ROIs.

Linear mixed-effects analyses results reflecting the relationship between SNR and SV are shown in Supplementary Table 25. Using MAGeT-Brain, a significant association between SV and SNR was identified within bilateral globus pallidus, bilateral thalamus, and left amygdala ROIs. Using FreeSurfer, a significant association between SV and SNR was identified within bilateral amygdala, bilateral globus pallidus, and bilateral striatum ROIs.

## 4. Discussion

The present study aimed to: 1) investigate the impact of an upgrade from a Siemens Trio to a Siemens Prisma MRI scanner and 2) compare a sequence with vNavs against a standard T1-weighted acquisition sequence. To the best of our knowledge, the current work represents the largest sample used to assess the impact of the Prisma upgrade on volumetric and morphological indices, the first to use CIVET and MAGeT-Brain to investigate this research question, and the first to examine neurochemical indices. Our results suggest high reliability in most investigated MRI outputs across the Prisma upgrade, although suboptimal reliability was observed in a minority of ROIs. Further, in certain ROIs, despite high reliability, CT and SV estimates differed across the Prisma upgrade. In addition, the present study is the first to explore vNav reliability and similarity analyses across multiple MRI visits, and use CIVET and MAGeT-Brain to address this research question, albeit in a sample that may not be representative of the clinical population in whom prospective motion correction would be most beneficial. The current work identified certain ROIs within which MPRAGE and vNav-MPRAGE sequences were most dissimilar.

The identification of particular ROIs within which the Prisma upgrade impacted reliability is relevant towards longitudinal MRI investigations. In the present work, we observed “poor” reliability in right isthmus cingulate gyrus CT and “fair” reliability in right lingual gyrus CT, as estimated by CIVET, and “fair” reliability of CT within the left transverse temporal, right rostral anterior cingulate, and right inferior occipital ROIs, as estimated by FreeSurfer. Also, SV acquired using FreeSurfer in the left amygdala and right thalamus rendered “fair” reliability, and ICCs within other structures did not reach statistical significance. For future longitudinal MRI investigations, interpretation of estimates of CT and SV within these ROIs merits caution, as they may be impacted in an inconsistent manner by the Prisma upgrade. Notably, it is possible that low reliability observed for CT and SV estimates across the Prisma upgrade in these ROIs may be in part attributable to substandard reliability of these measures in general. To this effect, the examination of Trio-Trio and Prisma-Prisma reliability complements the interpretation of the Trio-Prisma results. For example, in Trio-Trio investigations, CIVET-acquired right lateral orbitofrontal CT had “fair” reliability, and in Prisma-Prisma investigations, FreeSurfer-acquired bilateral globus pallidus volume had “fair” or “poor” reliability. Interestingly, cortical ROIs with lower ICCs include those believed to be affected by distortions (e.g. orbitofrontal areas) as well as those with a thin cortex that may contribute to poorer segmentation (e.g. V1). Thus, low reliability observed in these regions within Trio-Prisma pairs may not solely be a result of the upgrade.

Despite demonstrating “excellent” or “good” reliability across the Prisma upgrade, CT increases, as estimated by CIVET, were identified in 5 frontal ROIs (i.e. bilateral lateral orbitofrontal, bilateral medial orbitofrontal, right lateral frontal orbitalis), 3 right temporal ROIs (i.e. inferior temporal, middle temporal, superior temporal), and the right insula. Likewise, CT increases, as estimated by FreeSurfer, were identified within the left caudal anterior cingulate and the right insula; however, CT decreases were identified using FreeSurfer within 3 parietal ROIs (i.e. bilateral pericalcarine, left superior parietal). That being said, it is noteworthy that CT increases within frontal brain regions such as the medial orbitofrontal and lateral orbitofrontal ROIs were also observed using FreeSurfer, yet failed to achieve statistical significance after correction for multiple comparisons. Furthermore, analyses from the current work suggest that the frontal and temporal CT increases identified by CIVET, along with the parietal CT decreases and temporal and cingulate CT increases identified by FreeSurfer, are related to SNR increases that resulted from the Trio to Prisma upgrade. This phenomenon is in line with previous studies that have reported upon volume increases attributed to SNR increases following MRI scanner upgrades (Lee et al., 2019; Potvin et al., 2019; Shuter et al., 2008). Notably, orbitofrontal regions in particular are influenced by magnetic susceptibility differences in the air-tissue interface that exists between the nasal sinus cavity and the lower frontal cortex; this phenomenon similarly applies to the temporal cortices and is supported by magnetic field B_0_ mapping (de Graaf, 2019).

The findings of increased CT within frontal and cingulate ROIs and decreased CT within parietal ROIs following the Prisma upgrade is consistent with the current literature. In particular, Potvin and colleagues similarly investigated an upgrade from a Siemens Trio to the Prisma on three healthy male volunteers at three separate sites (Potvin et al., 2019). The authors analyzed T1-weighted images using FreeSurfer and, akin to the present study, parcellated ROIs using the DKT atlas. Increased thickness estimates were observed in several ROIs, predominantly within frontal regions, in addition to cingulate ROIs, and decreased thickness estimates were observed in left superior parietal and left entorhinal ROIs.

Moreover, in the present study, as assessed by MAGeT-Brain, decreases in left amygdala, left globus pallidus, and left hippocampus volumes, along with increases in bilateral thalamus volumes, were identified across the Prisma upgrade. Similarly, decreases in bilateral amygdala, bilateral globus pallidus, and bilateral striatum volumes were identified across the Prisma upgrade using FreeSurfer. These findings merit comparison to those of Potvin and colleagues, who identified increases in right amygdala, right caudate, left pallidum, and right putamen volumes (Potvin et al., 2019). Interestingly, the authors identified a negative correlation between subcortical contrast-to-noise ratio and subcortical volume, akin to the results of the current work.

Furthermore, in the vNav protocol of the present study, the comparison of vNav-MPRAGE and MPRAGE sequences was not designed to assess the effectiveness of vNav-MPRAGE sequences as a motion correction technique, but rather to help understand whether their use should be expected to impact estimates of volumetric and morphological indices, in a study with the expectation of a lack of substantial participant motion. Overall, the current work suggests high reliability (i.e. similarity) in most volumetric and morphological indices between vNav-MPRAGE and MPRAGE sequences. However, similarity analyses comparing vNav-MPRAGE and MPRAGE sequences demonstrated reduced reliability in CT estimates within the right entorhinal cortex (CIVET and FreeSurfer) and the right isthmus cingulate (CIVET). Notably, the finding of low reliability within the right cingulate ROIs between these sequences may be in part explained by the results of test-retest reliability analyses of the vNav-MPRAGE sequences, which similarly identified low reliability within cingulate areas. This may be attributed to poor midline classification, which may be a consequence of an asymmetric model not having been used for CT analyses. These findings are applicable to studies that are composed of one participant subgroup that is more likely to move during an MRI scan and one participant subgroup that is more likely to remain still (e.g. case-control studies comparing patients with autism spectrum disorder and healthy controls).

Further, CT estimated using CIVET was lower in vNav-MPRAGE compared to MPRAGE sequences within 4 temporal ROIs (i.e. bilateral parahippocampal, right superior temporal, right middle temporal), and right precentral, right inferior parietal, and right precuneus ROIs, yet was higher in vNav-MPRAGE compared to MPRAGE sequences within left medial orbitofrontal and left postcentral ROIs. In contrast, CT estimated using FreeSurfer was higher in vNav-MPRAGE compared to MPRAGE sequences within 5 parietal ROIs (i.e. bilateral postcentral, left superior parietal, left inferior parietal, right pericalcarine), 3 frontal ROIs (i.e. right superior frontal, right rostral middle frontal, right lateral frontal triangularis), and the left transverse temporal ROI, yet was lower in vNav-MPRAGE sequences in the right inferior occipital ROI. It is noteworthy that this comparison may be in part affected by TI differences.

In reference to previous literature investigating vNav sequences, their improvement in cases with deliberate motion is evident (Andersen et al., 2019). Seminal work by Tisdall *et al.* reported that reductions in dice overlap ratios in FreeSurfer-acquired cortical and subcortical volumes between no-motion and motion acquisitions improved when vNav sequences were used for the latter (Tisdall et al., 2012). More recently, Ai *et al.* collected two MPRAGE and two vNav acquisitions within the same MRI session (Ai et al., 2019). Using ICCs, the authors found that reliability in median CT across FreeSurfer’s 62 cortical regions, as assessed by MindBoggle, was highest within vNav sequences and lowest in MPRAGE sequences. Also, ICCs were not as sensitive to motion in vNav sequences, a phenomenon that was identified to be particularly relevant for CT, which typically rendered lower ICCs than area and volume. Finally, the authors demonstrated that the reproducibility (termed similarity in the current study) between MPRAGE and vNav-MPRAGE sequences was good (ICCs on average > 0.6) and exceeded the reproducibility between MPRAGE and MPRAGE pairs. Thus, the authors concluded that morphometric results obtained from MPRAGE sequences should be replicated using vNav-MPRAGE sequences.

The ^1^H-MRS component of the current study demonstrated “excellent” or “good” reliability across the neurometabolites most commonly investigated within the literature, including Glu, Glx, Ins, Cho, NAA, and tNAA. Importantly, such findings were largely consistent across each segment of the Prisma Upgrade protocol (e.g. Trio-Trio, Trio-Prisma, Prisma-Prisma). That being said, amongst these neurometabolites, it deserves mention that the reliability of Cho levels increased in the Prisma-Prisma segment and the reliability of Ins levels decreased in the Trio-Prisma segment. Furthermore, the other ^1^H-MRS indices investigated in the current study included GABA, Gln, GSH, Lac, and NAAG levels (neurometabolites often excluded from 3T investigations not employing editing sequences (Wijtenburg et al., 2015)) as well as FWHM and SNR. While the failure to achieve statistical significance within several of the analyses concerning these indices renders their interpretation to be somewhat speculative, findings from the current work appear to suggest that the reliabilities of GABA, NAAG, and FWHM were highest in the Prisma-Prisma component, the reliabilities of Gln levels and SNR decreased in the Trio-Prisma component, and the reliabilities of GSH and Lac levels were highest in the Trio-Trio component.

The present work should be considered in light of several limitations. First, it should be mentioned that the present work aimed to standardize pre- and post-scans and thus does not comprehensively assess the full benefits of the PRISMA upgrade. Second, the potential influence of differences between imaging parameters between pre- and post-scans as well as between MPRAGE and vNav-MPRAGE sequences cannot be discounted. Third, it is notable that the advantages of vNavs may be more apparent in instances with marked movement, such as in populations that are more prone to movement or cases of forced movement. Indeed, prospective motion correction using vNavs may have a negative impact on image quality in cases where participants are relatively still. In the current work, motion was not directly assessed. Overall, the current work examines a participant sample that may be unrepresentative of the clinical population in whom prospective motion correction is beneficial; rather, it is likely representative of the control sample in large-scale case-control studies. Finally, given the test-retest reliability nature of this investigation, exclusion of outputs from analysis on the basis of quality was not performed in an effort not to bias comparisons.

## 5. Conclusions

Taken together, as part of its Prisma Upgrade protocol, the current study investigated the impact that the Prisma upgrade may have on common MRI outputs (e.g. CT, SV, ^1^H-MRS indices). High reliability was identified in most of the MRI outputs that were investigated across the Prisma upgrade. However, suboptimal reliability and differences were observed for CT or SV estimates within certain ROIs across the Prisma upgrade. Further, as assessed by CIVET and FreeSurfer, CT increases in frontal, temporal, and cingulate ROIs, along with CT decreases in parietal ROIs may ensue across a Prisma upgrade, which appear to be linked to increases in SNR. Additionally, as assessed by MAGeT-Brain and FreeSurfer, volume decreases in the amygdala, globus pallidus, hippocampus, and striatum may result from a Prisma upgrade, along with volume increases in the thalamus; these alterations were also found to be linked to SNR. These findings should be particularly considered in the design of longitudinal study designs integrating measures from before and after the upgrade, and are also highly relevant to large-scale initiatives incorporating data from multiple sites. It would be the authors’ suggestion for scanner upgrade to be considered as a covariate (i.e. random effect) in statistical analyses.

In addition, as part of its vNav Protocol, the current study reported upon ROIs within which CT and SV estimates had suboptimal reliability or differed between MPRAGE and vNav-MPRAGE sequences. As assessed by CIVET, several ROIs within which MPRAGE estimates exceeded vNav-MPRAGE estimates were identified, predominantly involving temporal areas. In contrast, as assessed by FreeSurfer, higher CT estimates in vNav-MPRAGE acquisitions were identified, including parietal, frontal, and temporal ROIs. It is worth mentioning that the key advantages of vNav sequences may be more apparent in instances with marked head movement, such as populations that are more prone to head movement (e.g. children, certain patient populations) or cases of forced head movement. Thus, such investigations necessitate further examination.

## Acknowledgements

Support from the CIC and Douglas Research Centre and its staff who supported this work as well as the FRQS PSRv2 Grant and HBHL Platform Grant that supported the Prisma-Fit upgrade at the Douglas CIC. Data processing was partially performed using the gpc resources of the SciNet HPC Consortium.

---

**Supplementary Figure 1.**
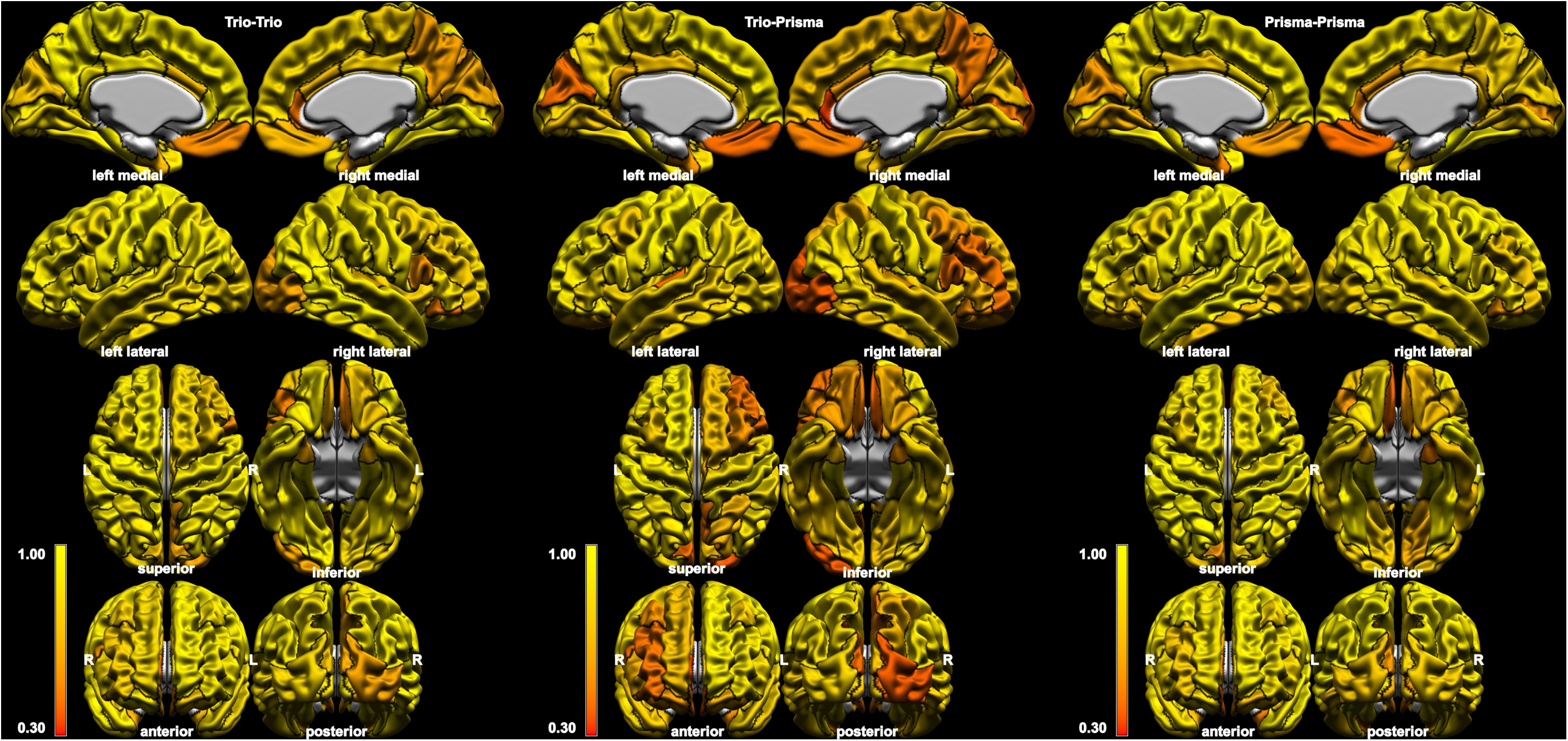
Intraclass correlation coefficients of cortical thickness estimated by FreeSurfer within regions of interest defined by the Desikan-Killiany-Tourville atlas (black boundaries). Results are shown for pairs consisting of two T1-weighted images from a Trio machine (left), Prisma machine (right), and one of each (middle).

**Supplementary Figure 2.**
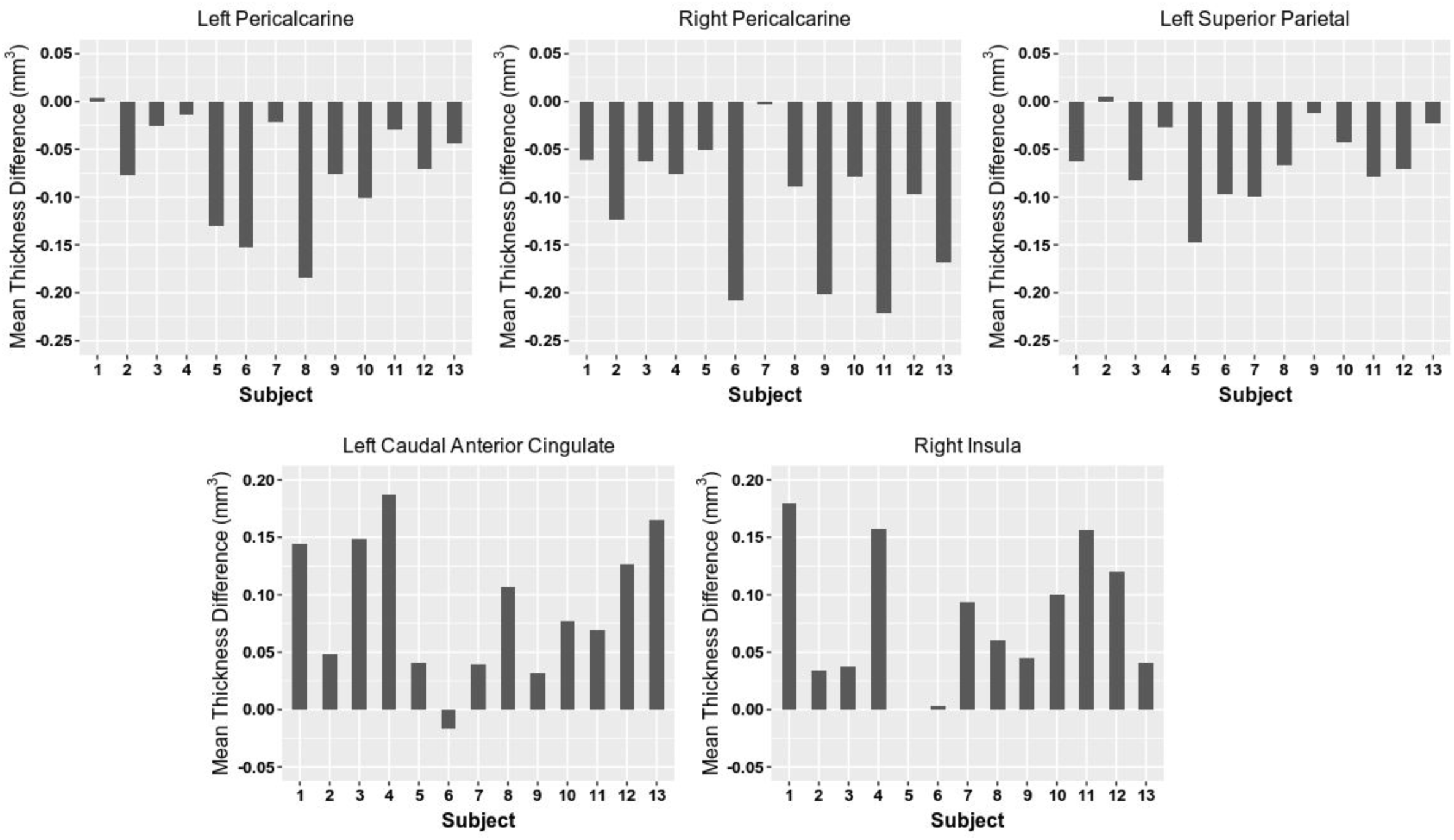
Cortical thickness estimated by FreeSurfer within regions of interest that achieved statistical significance in linear mixed-effects analyses. Mean thickness difference represents Prisma estimate minus Trio estimate.

**Supplementary Figure 3.**
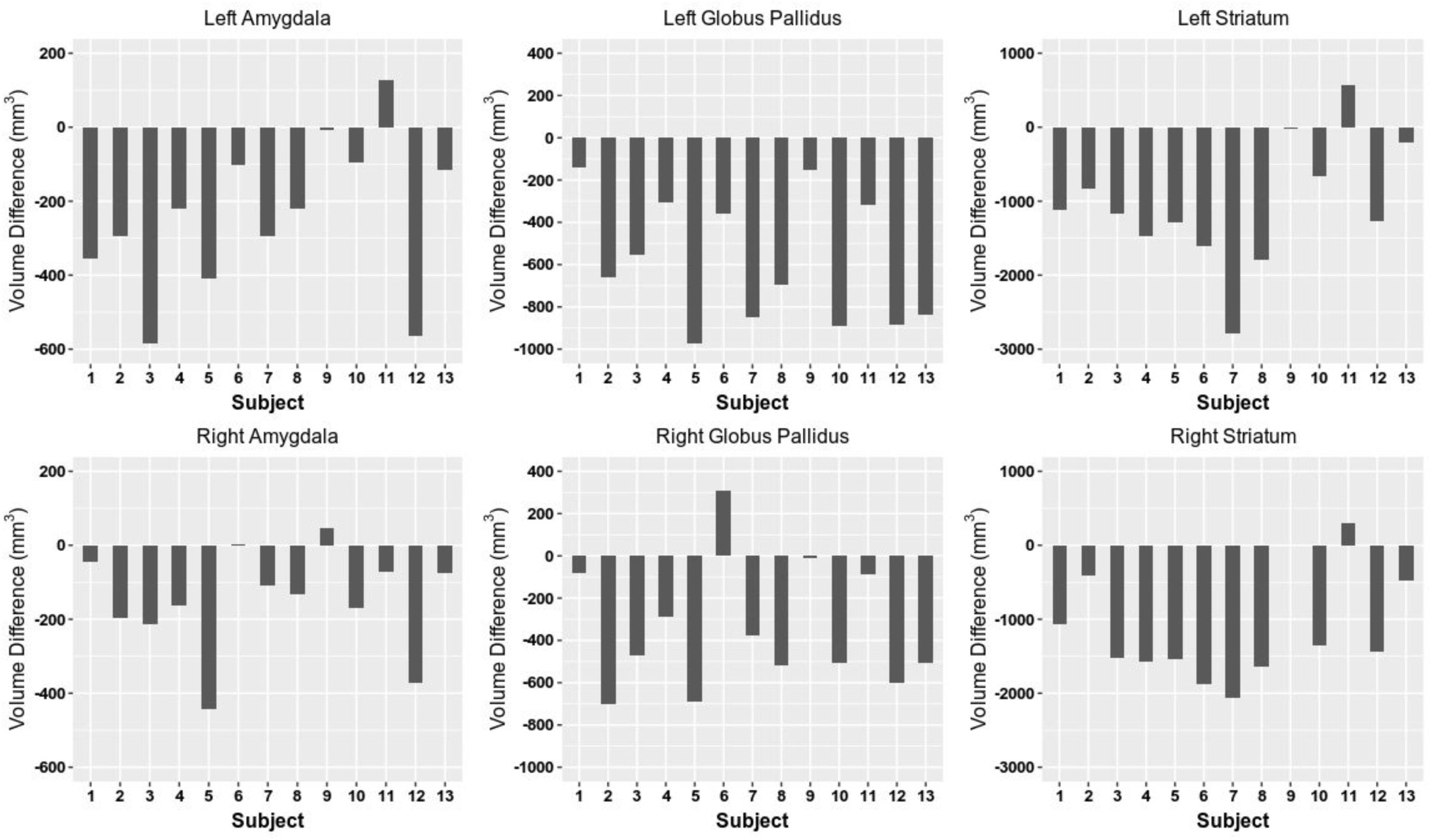
Structure volume estimated by FreeSurfer within regions of interest that achieved statistical significance in linear mixed-effects analyses. Volume difference represents Prisma estimate minus Trio estimate.

**Supplementary Figure 4.**
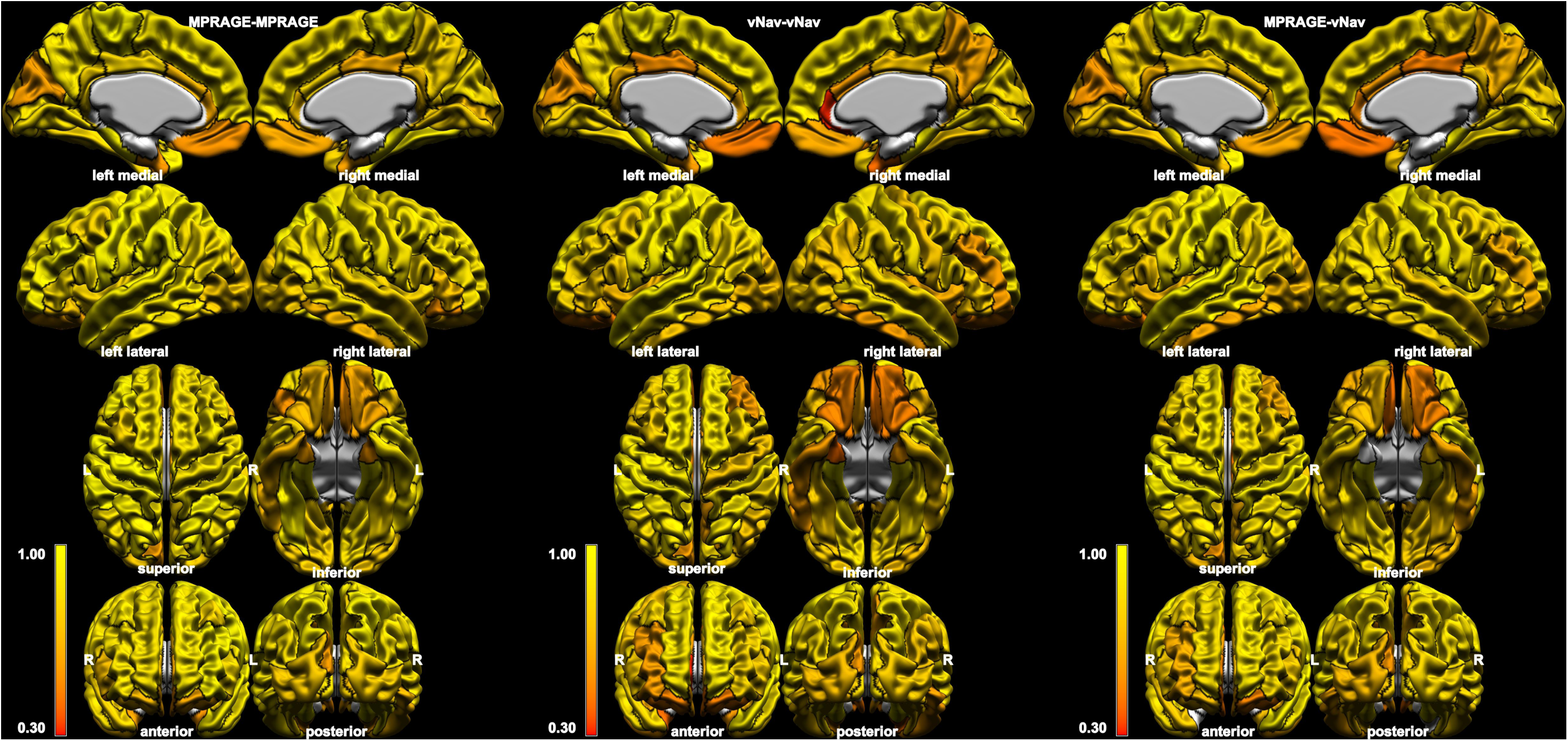
Intraclass correlation coefficients of cortical thickness estimated by FreeSurfer within regions of interest defined by the Desikan-Killiany-Tourville atlas (black boundaries). Results are shown for pairs consisting of two T1-weighted images (left), two volumetric navigator T1-weighted images (middle), and one of each (right). Note: right entorhinal cortex not included in MPRAGE-vNav, as ICC did not reach statistical significance (ICC=0.28, p=0.163).

**Supplementary Figure 5.**
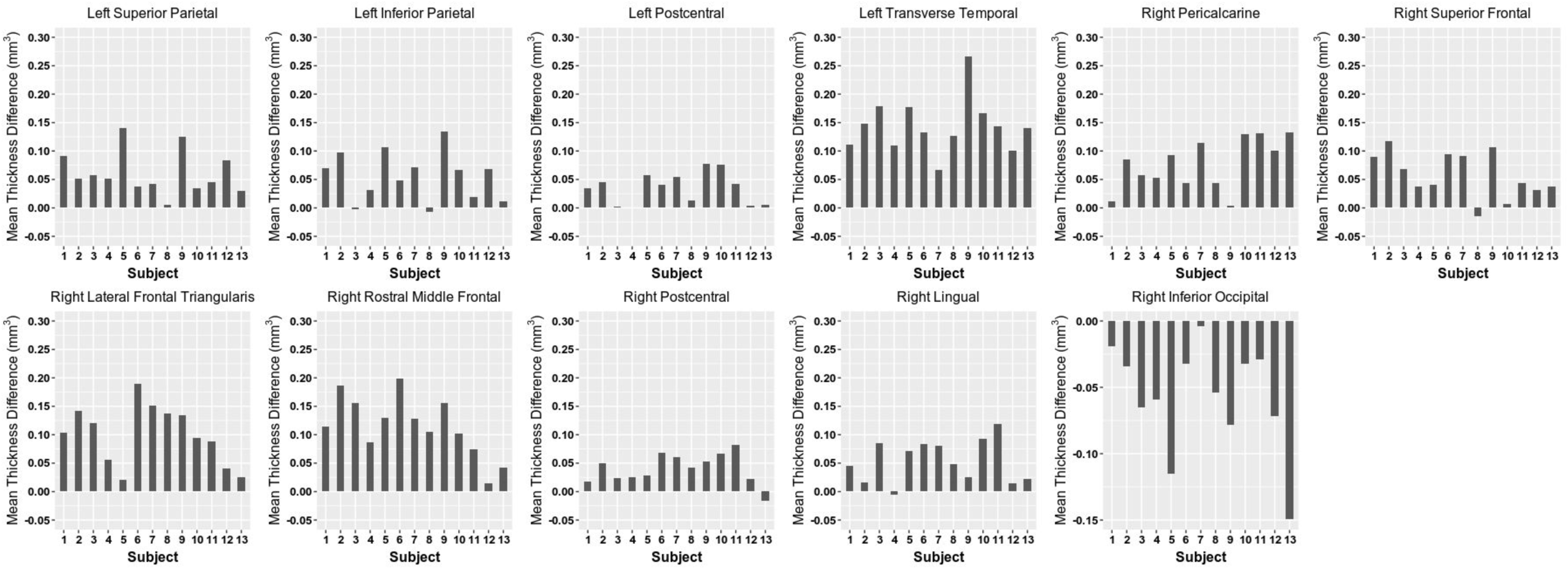
Cortical thickness estimated by the FreeSurfer pipeline within regions of interest that achieved statistical significance in linear mixed-effects analyses. Mean thickness difference represents vNav-MPRAGE estimate minus MPRAGE estimate.

**Supplementary Figure 6.**
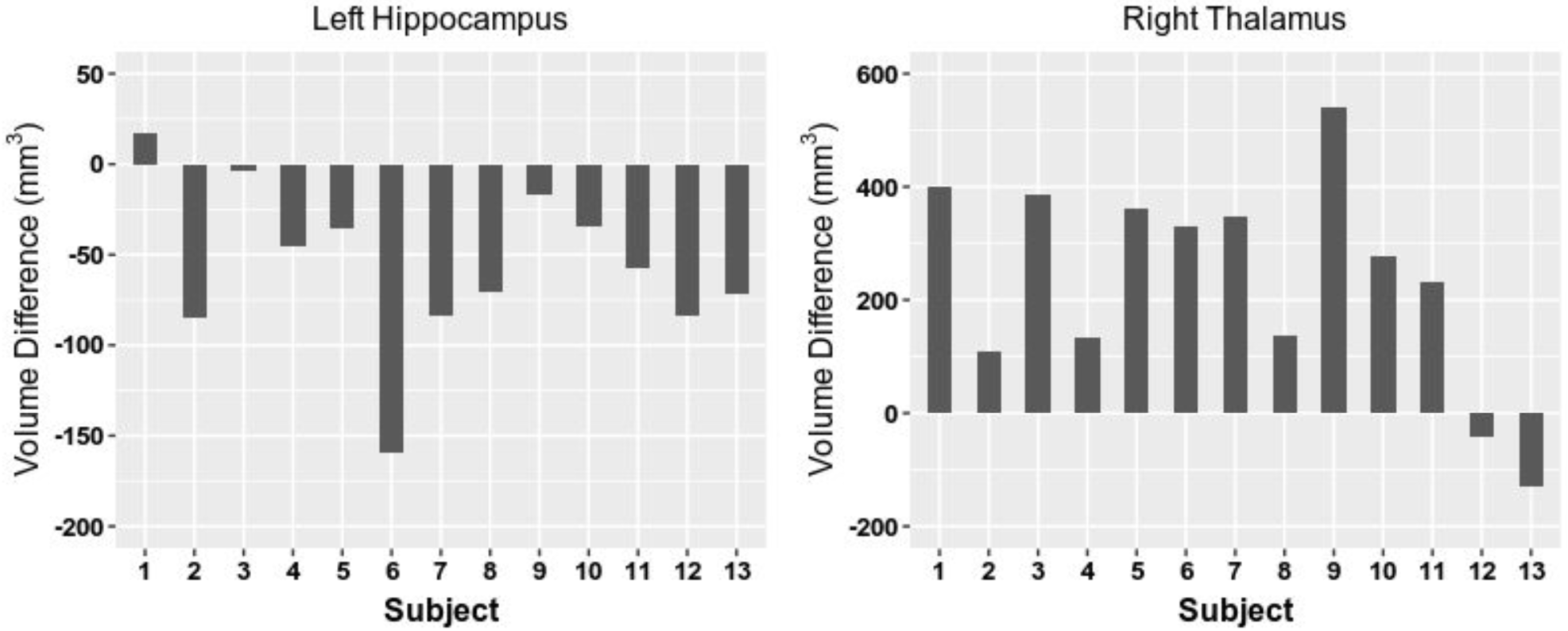
Structure volume estimated by the FreeSurfer pipeline within regions of interest that achieved statistical significance in linear mixed-effects analyses. Mean thickness difference represents vNav-MPRAGE estimate minus MPRAGE estimate.

## Supplementary Table Outline and Organization

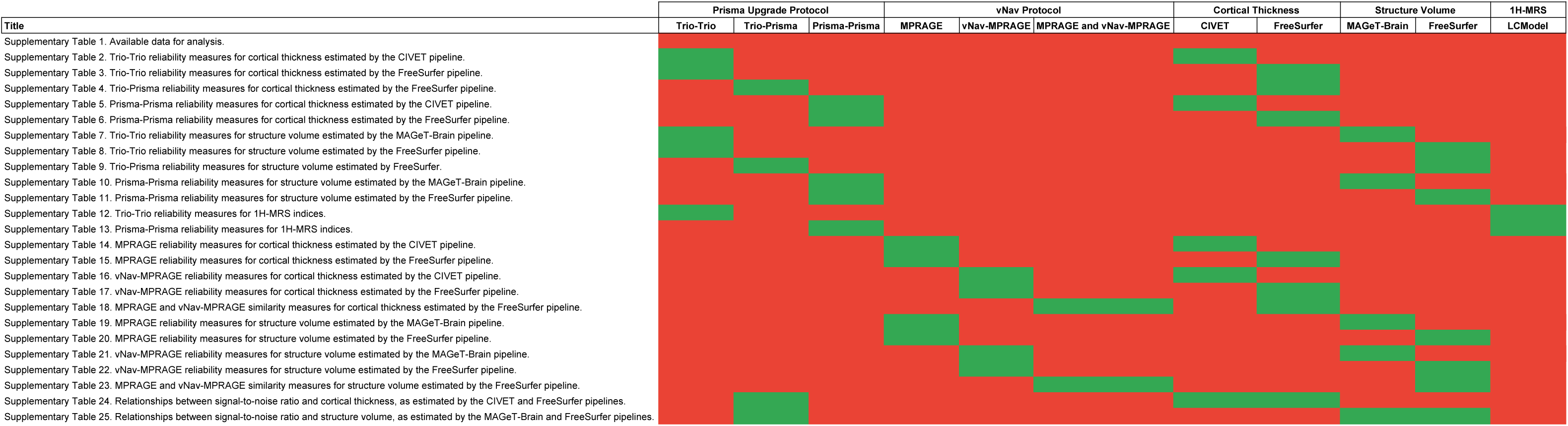

**Supplementary Table 1.** Available data for analysis.

| Visits | Sequence | MRI Measure | Analysis Pipeline | Number of Pairs | Mean Days<br>between<br>Acquisitions (SD)<br>[range] | Clarification Regarding Number of Pairs |
| --- | --- | --- | --- | --- | --- | --- |
| Test-retest reliability analyses - Prisma Upgrade Protocol |  |  |  |  |  |  |
| Trio - Trio | T1w (MPRAGE) | CT | CIVET | 10 | 5.2 (4.1) [1-15] | Of 13 subjects included in the Prisma Upgrade Protocol, 10 completed 2 Trio visits |
|  |  |  | FS | 10 |  |  |
|  |  | SV | MAGeT | 10 |  |  |
|  |  |  | FS | 10 |  |  |
|  | 1H-MRS | Met | LCModel | 9 | 5.6 (4.2) [1-15] | Of 13 subjects included in the Prisma Upgrade Protocol, 9 completed 2 Trio visits |
| Trio - Prisma | T1w (MPRAGE) | CT | CIVET | 13 | 114.9 (12.8) [91-133] | Thirteen subjects were included in the Prisma Upgrade Protocol, each of which had at least 1 Trio visit and 1 Prisma visit |
|  |  |  | FS | 13 |  |  |
|  |  | SV | MAGeT | 13 |  |  |
|  |  |  | FS | 13 |  |  |
|  | 1H-MRS | Met | LCModel | 8 | 382.5 (23.2) [364-434] | Of 13 subjects included in the Prisma Upgrade Protocol, 8 had both at least 1 Trio visit and 1 Prisma visit. Notably, the Prisma visit was acquired during the vNav Protocol |
| Prisma - Prisma | T1w (MPRAGE) | CT | CIVET | 16 | 10.06 (8.4) [1-34] | Of the 13 subjects included in the Prisma Upgrade Protocol, 10 completed 2 Prisma visits. One subject who failed to complete 2 Prisma visits in the Prisma Upgrade Protocol did so within the vNav Protocol. Five new subjects completed 2 Prisma visits within the vNav Protocol. |
|  |  |  | FS | 16 |  |  |
|  |  | SV | MAGeT | 16 |  |  |
|  |  |  | FS | 16 |  |  |
|  | 1H-MRS | Met | LCModel | 13 | 4.3 (3.4) [1-12] | Of the 13 subjects included in the vNav Protocol, all completed 2 Prisma visits |
| Test-retest reliability analyses - vNav Protocol |  |  |  |  |  |  |
| Prisma - Prisma | T1w (MPRAGE) | CT | CIVET | 13 | 4.3 (3.5) [1-12] | Thirteen subjects were included in the vNav Protocol, each of which completed 2 MPRAGE and 2 vNav-MPRAGE acquisitions |
|  |  |  | FS | 13 |  |  |
|  |  | SV | MAGeT | 13 |  |  |
|  |  |  | FS | 13 |  |  |
|  | vNav T1w (vNav-MPRAGE) | CT | CIVET | 13 | 4.3 (3.5) [1-12] |  |
|  |  |  | FS | 13 |  |  |
|  |  | SV | MAGeT | 13 |  |  |
|  |  |  | FS | 13 |  |  |
| Similarity analyses - vNav Protocol |  |  |  |  |  |  |
| Prisma - Prisma | T1w (MPRAGE) and vNav T1w (vNav-MPRAGE) | CT | CIVET | 13 | N/A |  |
|  |  |  | FS | 13 |  |  |
|  |  | SV | MAGeT | 13 |  |  |
|  |  |  | FS | 13 |  |  |

**Supplementary Table 2.** Trio-Trio reliability measures for cortical thickness estimated by the CIVET pipeline.

| ROI | CIVET |  |  |  |  |  |  |  |  |  |  |  |  |  |  |  |
| --- | --- | --- | --- | --- | --- | --- | --- | --- | --- | --- | --- | --- | --- | --- | --- | --- |
|  | Left Hemisphere |  |  |  |  |  |  |  | Right Hemisphere |  |  |  |  |  |  |  |
|  | Timepoint 1<br>Mean (SD) | Timepoint 2<br>Mean (SD) | APD | ICC (2,1) | ICC (3,1) | p-value (ICC) | t-value (lmer) | p-value (lmer) | Timepoint 1<br>Mean (SD) | Timepoint 2<br>Mean (SD) | APD | ICC (2,1) | ICC (3,1) | p-value (ICC) | t-value (lmer) | p-value (lmer) |
| Caudal Anterior Cingulate | 2.78 (0.13) | 2.81 (0.10) | 1.74 | 0.89 | 0.91 | <0.001 | 1.99 | 0.077 | 2.70 (0.14) | 2.73 (0.15) | 1.71 | 0.93 | 0.94 | <0.001 | 1.66 | 0.130 |
| Caudal Middle Frontal | 3.06 (0.09) | 3.06 (0.08) | 0.71 | 0.96 | 0.96 | <0.001 | 0.32 | 0.756 | 3.02 (0.09) | 3.03 (0.10) | 0.97 | 0.92 | 0.92 | <0.001 | 0.81 | 0.437 |
| Cuneus | 2.51 (0.09) | 2.51 (0.11) | 1.43 | 0.89 | 0.89 | <0.001 | 0.02 | 0.988 | 2.60 (0.06) | 2.59 (0.08) | 1.57 | 0.76 | 0.76 | 0.003 | -0.98 | 0.350 |
| Entorhinal Cortex | 3.83 (0.33) | 3.85 (0.26) | 3.61 | 0.85 | 0.85 | <0.001 | 0.53 | 0.609 | 4.00 (0.31) | 3.99 (0.20) | 3.00 | 0.84 | 0.84 | <0.001 | -0.17 | 0.867 |
| Fusiform Gyrus | 3.13 (0.13) | 3.15 (0.09) | 1.63 | 0.86 | 0.86 | <0.001 | 0.87 | 0.405 | 3.18 (0.15) | 3.17 (0.16) | 1.44 | 0.92 | 0.92 | <0.001 | -0.27 | 0.792 |
| Inferior Occipital Cortex | 2.67 (0.08) | 2.68 (0.07) | 0.89 | 0.94 | 0.94 | <0.001 | 1.41 | 0.192 | 2.82 (0.06) | 2.80 (0.08) | 1.20 | 0.84 | 0.86 | <0.001 | -1.74 | 0.117 |
| Inferior Parietal | 2.96 (0.11) | 2.96 (0.09) | 0.71 | 0.97 | 0.97 | <0.001 | 0.25 | 0.807 | 3.04 (0.10) | 3.03 (0.10) | 0.66 | 0.96 | 0.96 | <0.001 | -1.53 | 0.160 |
| Inferior Temporal | 3.12 (0.07) | 3.15 (0.12) | 1.76 | 0.71 | 0.73 | 0.006 | 1.34 | 0.214 | 3.26 (0.13) | 3.23 (0.13) | 1.60 | 0.85 | 0.86 | <0.001 | -1.54 | 0.158 |
| Insula | 3.59 (0.19) | 3.59 (0.21) | 1.72 | 0.93 | 0.93 | <0.001 | -0.11 | 0.911 | 3.58 (0.21) | 3.53 (0.17) | 2.02 | 0.90 | 0.92 | <0.001 | -1.86 | 0.096 |
| Isthmus Cingulate Gyrus | 3.07 (0.24) | 3.10 (0.21) | 3.71 | 0.79 | 0.79 | 0.002 | 0.67 | 0.519 | 3.02 (0.16) | 3.05 (0.12) | 3.04 | 0.70 | 0.70 | 0.008 | 0.69 | 0.508 |
| Lateral Frontal Opercularis | 3.06 (0.06) | 3.07 (0.06) | 1.03 | 0.72 | 0.72 | 0.006 | 0.92 | 0.382 | 3.04 (0.08) | 3.04 (0.08) | 1.01 | 0.87 | 0.87 | <0.001 | -0.15 | 0.884 |
| Lateral Frontal Orbitalis | 3.09 (0.11) | 3.10 (0.11) | 1.18 | 0.90 | 0.90 | <0.001 | 0.80 | 0.446 | 3.16 (0.11) | 3.15 (0.10) | 1.51 | 0.86 | 0.87 | <0.001 | -1.04 | 0.327 |
| Lateral Frontal Triangularis | 2.95 (0.11) | 2.96 (0.09) | 0.89 | 0.94 | 0.94 | <0.001 | 1.06 | 0.317 | 2.94 (0.11) | 2.93 (0.12) | 1.29 | 0.92 | 0.92 | <0.001 | -0.45 | 0.665 |
| Lateral Orbitofrontal | 2.93 (0.08) | 2.93 (0.09) | 1.32 | 0.85 | 0.85 | <0.001 | 0.09 | 0.931 | 2.87 (0.06) | 2.84 (0.06) | 2.10 | 0.47 | 0.52 | 0.050 | -1.74 | 0.115 |
| Lingual Gyrus | 2.59 (0.10) | 2.58 (0.11) | 2.37 | 0.78 | 0.78 | 0.003 | -0.13 | 0.901 | 2.70 (0.07) | 2.66 (0.09) | 2.09 | 0.70 | 0.75 | 0.004 | -1.99 | 0.077 |
| Medial Orbitofrontal | 2.63 (0.10) | 2.63 (0.11) | 1.52 | 0.89 | 0.89 | <0.001 | 0.36 | 0.724 | 2.64 (0.12) | 2.65 (0.12) | 2.88 | 0.66 | 0.66 | 0.013 | 0.32 | 0.756 |
| Middle Temporal | 3.26 (0.11) | 3.27 (0.09) | 0.65 | 0.97 | 0.97 | <0.001 | 1.65 | 0.133 | 3.33 (0.10) | 3.31 (0.12) | 1.27 | 0.91 | 0.91 | <0.001 | -1.09 | 0.305 |
| Paracentral Gyrus | 2.98 (0.11) | 2.98 (0.12) | 0.92 | 0.96 | 0.96 | <0.001 | -0.59 | 0.570 | 2.99 (0.06) | 2.99 (0.09) | 1.18 | 0.88 | 0.88 | <0.001 | 0.18 | 0.863 |
| Parahippocampal | 2.73 (0.10) | 2.77 (0.12) | 1.99 | 0.81 | 0.85 | <0.001 | 2.10 | 0.065 | 2.78 (0.13) | 2.76 (0.13) | 1.77 | 0.91 | 0.92 | <0.001 | -1.19 | 0.266 |
| Pericalcarine | 2.42 (0.12) | 2.44 (0.10) | 2.52 | 0.75 | 0.75 | 0.004 | 0.77 | 0.459 | 2.52 (0.10) | 2.47 (0.07) | 3.23 | 0.58 | 0.67 | 0.012 | -2.33 | 0.045 |
| Postcentral Gyrus | 2.68 (0.13) | 2.68 (0.14) | 0.60 | 0.99 | 0.99 | <0.001 | 0.70 | 0.500 | 2.70 (0.09) | 2.69 (0.11) | 0.89 | 0.96 | 0.96 | <0.001 | -0.96 | 0.360 |
| Posterior Cingulate | 2.94 (0.11) | 2.96 (0.09) | 1.82 | 0.76 | 0.76 | 0.003 | 0.91 | 0.384 | 2.92 (0.07) | 2.91 (0.06) | 1.18 | 0.78 | 0.79 | 0.002 | -1.32 | 0.218 |
| Precentral Gyrus | 3.09 (0.08) | 3.11 (0.07) | 1.05 | 0.82 | 0.83 | <0.001 | 1.27 | 0.236 | 3.06 (0.11) | 3.06 (0.11) | 0.92 | 0.95 | 0.95 | <0.001 | 0.31 | 0.762 |
| Precuneus | 2.92 (0.08) | 2.93 (0.09) | 0.72 | 0.96 | 0.96 | <0.001 | 1.09 | 0.306 | 3.00 (0.10) | 3.00 (0.09) | 1.13 | 0.91 | 0.91 | <0.001 | -0.20 | 0.845 |
| Rostral Anterior Cingulate | 3.00 (0.22) | 3.03 (0.16) | 3.09 | 0.85 | 0.85 | <0.001 | 1.04 | 0.327 | 2.85 (0.11) | 2.86 (0.13) | 2.87 | 0.69 | 0.69 | 0.010 | 0.44 | 0.672 |
| Rostral Middle Frontal | 2.97 (0.12) | 2.99 (0.12) | 1.21 | 0.93 | 0.95 | <0.001 | 2.19 | 0.056 | 2.89 (0.12) | 2.87 (0.12) | 1.45 | 0.91 | 0.91 | <0.001 | -0.69 | 0.506 |
| Superior Frontal Gyrus | 3.11 (0.13) | 3.13 (0.13) | 0.85 | 0.97 | 0.98 | <0.001 | 1.80 | 0.106 | 3.05 (0.11) | 3.06 (0.12) | 1.27 | 0.92 | 0.92 | <0.001 | 0.71 | 0.496 |
| Superior Parietal | 2.71 (0.08) | 2.73 (0.07) | 0.77 | 0.95 | 0.97 | <0.001 | 2.95 | 0.016 | 2.73 (0.10) | 2.73 (0.11) | 0.73 | 0.97 | 0.97 | <0.001 | -0.43 | 0.679 |
| Superior Temporal | 3.27 (0.10) | 3.28 (0.09) | 0.93 | 0.94 | 0.94 | <0.001 | 0.63 | 0.542 | 3.31 (0.10) | 3.30 (0.10) | 0.88 | 0.92 | 0.93 | <0.001 | -1.25 | 0.244 |
| Supramarginal Gyrus | 3.02 (0.14) | 3.03 (0.14) | 0.79 | 0.98 | 0.98 | <0.001 | 1.58 | 0.149 | 3.06 (0.15) | 3.05 (0.15) | 0.93 | 0.97 | 0.98 | <0.001 | -1.16 | 0.274 |
| Transverse Temporal | 3.00 (0.13) | 2.99 (0.12) | 1.06 | 0.95 | 0.96 | <0.001 | -1.53 | 0.161 | 3.05 (0.17) | 3.02 (0.14) | 1.82 | 0.91 | 0.92 | <0.001 | -1.60 | 0.143 |

APD, absolute percentage difference; ICC, intraclass correlation coefficient; lmer, linear mixed-effects model; ROI, region of interest; SD, standard deviation. Note: positive t-values represent cortical thickness increases.

**Supplementary Table 3.**
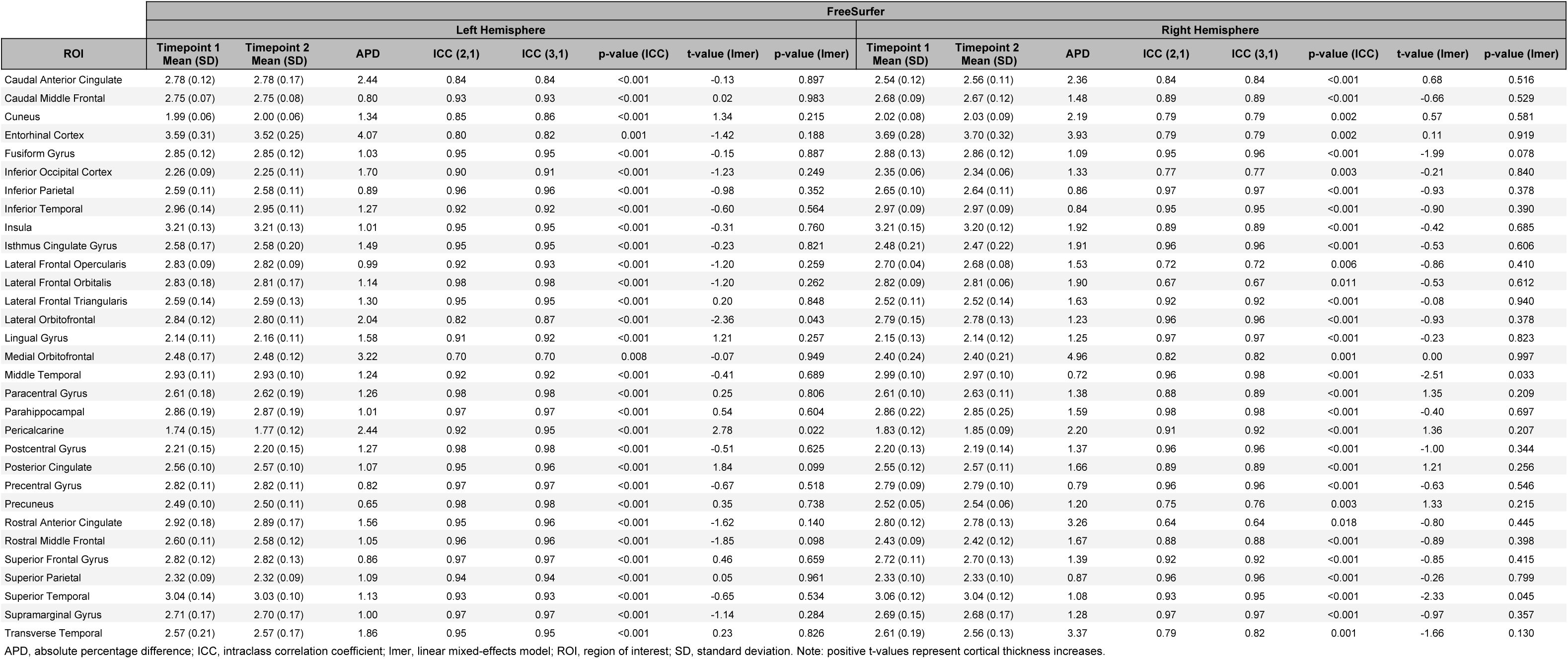
Trio-Trio reliability measures for cortical thickness estimated by the FreeSurfer pipeline.

| ROI | FreeSurfer |  |  |  |  |  |  |  |  |  |  |  |  |  |  |  |
| --- | --- | --- | --- | --- | --- | --- | --- | --- | --- | --- | --- | --- | --- | --- | --- | --- |
|  | Left Hemisphere |  |  |  |  |  |  |  | Right Hemisphere |  |  |  |  |  |  |  |
|  | Timepoint 1<br>Mean (SD) | Timepoint 2<br>Mean (SD) | APD | ICC (2,1) | ICC (3,1) | p-value (ICC) | t-value (lmer) | p-value (lmer) | Timepoint 1<br>Mean (SD) | Timepoint 2<br>Mean (SD) | APD | ICC (2,1) | ICC (3,1) | p-value (ICC) | t-value (lmer) | p-value (lmer) |
| Caudal Anterior Cingulate | 2.78 (0.12) | 2.78 (0.17) | 2.44 | 0.84 | 0.84 | <0.001 | -0.13 | 0.897 | 2.54 (0.12) | 2.56 (0.11) | 2.36 | 0.84 | 0.84 | <0.001 | 0.68 | 0.516 |
| Caudal Middle Frontal | 2.75 (0.07) | 2.75 (0.08) | 0.80 | 0.93 | 0.93 | <0.001 | 0.02 | 0.983 | 2.68 (0.09) | 2.67 (0.12) | 1.48 | 0.89 | 0.89 | <0.001 | -0.66 | 0.529 |
| Cuneus | 1.99 (0.06) | 2.00 (0.06) | 1.34 | 0.85 | 0.86 | <0.001 | 1.34 | 0.215 | 2.02 (0.08) | 2.03 (0.09) | 2.19 | 0.79 | 0.79 | 0.002 | 0.57 | 0.581 |
| Entorhinal Cortex | 3.59 (0.31) | 3.52 (0.25) | 4.07 | 0.80 | 0.82 | 0.001 | -1.42 | 0.188 | 3.69 (0.28) | 3.70 (0.32) | 3.93 | 0.79 | 0.79 | 0.002 | 0.11 | 0.919 |
| Fusiform Gyrus | 2.85 (0.12) | 2.85 (0.12) | 1.03 | 0.95 | 0.95 | <0.001 | -0.15 | 0.887 | 2.88 (0.13) | 2.86 (0.12) | 1.09 | 0.95 | 0.96 | <0.001 | -1.99 | 0.078 |
| Inferior Occipital Cortex | 2.26 (0.09) | 2.25 (0.11) | 1.70 | 0.90 | 0.91 | <0.001 | -1.23 | 0.249 | 2.35 (0.06) | 2.34 (0.06) | 1.33 | 0.77 | 0.77 | 0.003 | -0.21 | 0.840 |
| Inferior Parietal | 2.59 (0.11) | 2.58 (0.11) | 0.89 | 0.96 | 0.96 | <0.001 | -0.98 | 0.352 | 2.65 (0.10) | 2.64 (0.11) | 0.86 | 0.97 | 0.97 | <0.001 | -0.93 | 0.378 |
| Inferior Temporal | 2.96 (0.14) | 2.95 (0.11) | 1.27 | 0.92 | 0.92 | <0.001 | -0.60 | 0.564 | 2.97 (0.09) | 2.97 (0.09) | 0.84 | 0.95 | 0.95 | <0.001 | -0.90 | 0.390 |
| Insula | 3.21 (0.13) | 3.21 (0.13) | 1.01 | 0.95 | 0.95 | <0.001 | -0.31 | 0.760 | 3.21 (0.15) | 3.20 (0.12) | 1.92 | 0.89 | 0.89 | <0.001 | -0.42 | 0.685 |
| Isthmus Cingulate Gyrus | 2.58 (0.17) | 2.58 (0.20) | 1.49 | 0.95 | 0.95 | <0.001 | -0.23 | 0.821 | 2.48 (0.21) | 2.47 (0.22) | 1.91 | 0.96 | 0.96 | <0.001 | -0.53 | 0.606 |
| Lateral Frontal Opercularis | 2.83 (0.09) | 2.82 (0.09) | 0.99 | 0.92 | 0.93 | <0.001 | -1.20 | 0.259 | 2.70 (0.04) | 2.68 (0.08) | 1.53 | 0.72 | 0.72 | 0.006 | -0.86 | 0.410 |
| Lateral Frontal Orbitalis | 2.83 (0.18) | 2.81 (0.17) | 1.14 | 0.98 | 0.98 | <0.001 | -1.20 | 0.262 | 2.82 (0.09) | 2.81 (0.06) | 1.90 | 0.67 | 0.67 | 0.011 | -0.53 | 0.612 |
| Lateral Frontal Triangularis | 2.59 (0.14) | 2.59 (0.13) | 1.30 | 0.95 | 0.95 | <0.001 | 0.20 | 0.848 | 2.52 (0.11) | 2.52 (0.14) | 1.63 | 0.92 | 0.92 | <0.001 | -0.08 | 0.940 |
| Lateral Orbitofrontal | 2.84 (0.12) | 2.80 (0.11) | 2.04 | 0.82 | 0.87 | <0.001 | -2.36 | 0.043 | 2.79 (0.15) | 2.78 (0.13) | 1.23 | 0.96 | 0.96 | <0.001 | -0.93 | 0.378 |
| Lingual Gyrus | 2.14 (0.11) | 2.16 (0.11) | 1.58 | 0.91 | 0.92 | <0.001 | 1.21 | 0.257 | 2.15 (0.13) | 2.14 (0.12) | 1.25 | 0.97 | 0.97 | <0.001 | -0.23 | 0.823 |
| Medial Orbitofrontal | 2.48 (0.17) | 2.48 (0.12) | 3.22 | 0.70 | 0.70 | 0.008 | -0.07 | 0.949 | 2.40 (0.24) | 2.40 (0.21) | 4.96 | 0.82 | 0.82 | 0.001 | 0.00 | 0.997 |
| Middle Temporal | 2.93 (0.11) | 2.93 (0.10) | 1.24 | 0.92 | 0.92 | <0.001 | -0.41 | 0.689 | 2.99 (0.10) | 2.97 (0.10) | 0.72 | 0.96 | 0.98 | <0.001 | -2.51 | 0.033 |
| Paracentral Gyrus | 2.61 (0.18) | 2.62 (0.19) | 1.26 | 0.98 | 0.98 | <0.001 | 0.25 | 0.806 | 2.61 (0.10) | 2.63 (0.11) | 1.38 | 0.88 | 0.89 | <0.001 | 1.35 | 0.209 |
| Parahippocampal | 2.86 (0.19) | 2.87 (0.19) | 1.01 | 0.97 | 0.97 | <0.001 | 0.54 | 0.604 | 2.86 (0.22) | 2.85 (0.25) | 1.59 | 0.98 | 0.98 | <0.001 | -0.40 | 0.697 |
| Pericalcarine | 1.74 (0.15) | 1.77 (0.12) | 2.44 | 0.92 | 0.95 | <0.001 | 2.78 | 0.022 | 1.83 (0.12) | 1.85 (0.09) | 2.20 | 0.91 | 0.92 | <0.001 | 1.36 | 0.207 |
| Postcentral Gyrus | 2.21 (0.15) | 2.20 (0.15) | 1.27 | 0.98 | 0.98 | <0.001 | -0.51 | 0.625 | 2.20 (0.13) | 2.19 (0.14) | 1.37 | 0.96 | 0.96 | <0.001 | -1.00 | 0.344 |
| Posterior Cingulate | 2.56 (0.10) | 2.57 (0.10) | 1.07 | 0.95 | 0.96 | <0.001 | 1.84 | 0.099 | 2.55 (0.12) | 2.57 (0.11) | 1.66 | 0.89 | 0.89 | <0.001 | 1.21 | 0.256 |
| Precentral Gyrus | 2.82 (0.11) | 2.82 (0.11) | 0.82 | 0.97 | 0.97 | <0.001 | -0.67 | 0.518 | 2.79 (0.09) | 2.79 (0.10) | 0.79 | 0.96 | 0.96 | <0.001 | -0.63 | 0.546 |
| Precuneus | 2.49 (0.10) | 2.50 (0.11) | 0.65 | 0.98 | 0.98 | <0.001 | 0.35 | 0.738 | 2.52 (0.05) | 2.54 (0.06) | 1.20 | 0.75 | 0.76 | 0.003 | 1.33 | 0.215 |
| Rostral Anterior Cingulate | 2.92 (0.18) | 2.89 (0.17) | 1.56 | 0.95 | 0.96 | <0.001 | -1.62 | 0.140 | 2.80 (0.12) | 2.78 (0.13) | 3.26 | 0.64 | 0.64 | 0.018 | -0.80 | 0.445 |
| Rostral Middle Frontal | 2.60 (0.11) | 2.58 (0.12) | 1.05 | 0.96 | 0.96 | <0.001 | -1.85 | 0.098 | 2.43 (0.09) | 2.42 (0.12) | 1.67 | 0.88 | 0.88 | <0.001 | -0.89 | 0.398 |
| Superior Frontal Gyrus | 2.82 (0.12) | 2.82 (0.13) | 0.86 | 0.97 | 0.97 | <0.001 | 0.46 | 0.659 | 2.72 (0.11) | 2.70 (0.13) | 1.39 | 0.92 | 0.92 | <0.001 | -0.85 | 0.415 |
| Superior Parietal | 2.32 (0.09) | 2.32 (0.09) | 1.09 | 0.94 | 0.94 | <0.001 | 0.05 | 0.961 | 2.33 (0.10) | 2.33 (0.10) | 0.87 | 0.96 | 0.96 | <0.001 | -0.26 | 0.799 |
| Superior Temporal | 3.04 (0.14) | 3.03 (0.10) | 1.13 | 0.93 | 0.93 | <0.001 | -0.65 | 0.534 | 3.06 (0.12) | 3.04 (0.12) | 1.08 | 0.93 | 0.95 | <0.001 | -2.33 | 0.045 |
| Supramarginal Gyrus | 2.71 (0.17) | 2.70 (0.17) | 1.00 | 0.97 | 0.97 | <0.001 | -1.14 | 0.284 | 2.69 (0.15) | 2.68 (0.17) | 1.28 | 0.97 | 0.97 | <0.001 | -0.97 | 0.357 |
| Transverse Temporal | 2.57 (0.21) | 2.57 (0.17) | 1.86 | 0.95 | 0.95 | <0.001 | 0.23 | 0.826 | 2.61 (0.19) | 2.56 (0.13) | 3.37 | 0.79 | 0.82 | 0.001 | -1.66 | 0.130 |

APD, absolute percentage difference; ICC, intraclass correlation coefficient; lmer, linear mixed-effects model; ROI, region of interest; SD, standard deviation. Note: positive t-values represent cortical thickness increases.

**Supplementary Table 4.** Trio-Prisma reliability measures for cortical thickness estimated by the FreeSurfer pipeline.

| ROI | FreeSurfer |  |  |  |  |  |  |  |  |  |  |  |  |  |  |  |
| --- | --- | --- | --- | --- | --- | --- | --- | --- | --- | --- | --- | --- | --- | --- | --- | --- |
|  | Left Hemisphere |  |  |  |  |  |  |  | Right Hemisphere |  |  |  |  |  |  |  |
|  | Timepoint 1<br>Mean (SD) | Timepoint 2<br>Mean (SD) | APD | ICC (2,1) | ICC (3,1) | p-value (ICC) | t-value (lmer) | p-value (lmer) | Timepoint 1<br>Mean (SD) | Timepoint 2<br>Mean (SD) | APD | ICC (2,1) | ICC (3,1) | p-value (ICC) | t-value (lmer) | p-value (lmer) |
| Caudal Anterior Cingulate | 2.79 (0.16) | 2.88 (0.14) | 3.30 | 0.78 | 0.91 | <0.001 | 5.27 | <0.001* | 2.54 (0.11) | 2.61 (0.15) | 3.67 | 0.68 | 0.76 | <0.001 | 2.70 | 0.019 |
| Caudal Middle Frontal | 2.74 (0.08) | 2.77 (0.08) | 1.37 | 0.83 | 0.88 | <0.001 | 2.78 | 0.017 | 2.65 (0.12) | 2.65 (0.13) | 2.73 | 0.73 | 0.73 | 0.001 | 0.12 | 0.907 |
| Cuneus | 2.00 (0.05) | 1.97 (0.07) | 2.43 | 0.59 | 0.62 | 0.009 | -1.66 | 0.124 | 2.02 (0.09) | 1.98 (0.09) | 2.49 | 0.76 | 0.85 | <0.001 | -3.31 | 0.006 |
| Entorhinal Cortex | 3.54 (0.24) | 3.51 (0.17) | 3.32 | 0.72 | 0.72 | 0.002 | -0.84 | 0.415 | 3.73 (0.29) | 3.66 (0.30) | 3.57 | 0.83 | 0.84 | <0.001 | -1.40 | 0.188 |
| Fusiform Gyrus | 2.84 (0.13) | 2.82 (0.15) | 1.43 | 0.92 | 0.93 | <0.001 | -1.16 | 0.270 | 2.85 (0.12) | 2.86 (0.12) | 1.85 | 0.87 | 0.87 | <0.001 | 0.69 | 0.505 |
| Inferior Occipital Cortex | 2.24 (0.10) | 2.27 (0.11) | 1.96 | 0.87 | 0.91 | <0.001 | 2.58 | 0.024 | 2.35 (0.06) | 2.37 (0.07) | 2.36 | 0.55 | 0.57 | 0.017 | 1.40 | 0.187 |
| Inferior Parietal | 2.57 (0.11) | 2.53 (0.13) | 1.68 | 0.89 | 0.94 | <0.001 | -3.75 | 0.003 | 2.61 (0.11) | 2.63 (0.11) | 1.30 | 0.94 | 0.94 | <0.001 | 1.51 | 0.157 |
| Inferior Temporal | 2.95 (0.10) | 2.98 (0.11) | 1.61 | 0.87 | 0.90 | <0.001 | 2.44 | 0.031 | 2.95 (0.09) | 2.99 (0.10) | 1.53 | 0.81 | 0.87 | <0.001 | 2.77 | 0.017 |
| Insula | 3.21 (0.12) | 3.26 (0.14) | 2.01 | 0.85 | 0.90 | <0.001 | 3.08 | 0.010 | 3.20 (0.10) | 3.27 (0.14) | 2.41 | 0.73 | 0.88 | <0.001 | 4.74 | <0.001* |
| Isthmus Cingulate Gyrus | 2.60 (0.18) | 2.63 (0.14) | 2.82 | 0.82 | 0.82 | <0.001 | 0.97 | 0.352 | 2.47 (0.20) | 2.51 (0.21) | 2.97 | 0.91 | 0.93 | <0.001 | 2.04 | 0.064 |
| Lateral Frontal Opercularis | 2.79 (0.10) | 2.81 (0.10) | 1.28 | 0.91 | 0.94 | <0.001 | 2.79 | 0.016 | 2.67 (0.08) | 2.69 (0.07) | 1.88 | 0.62 | 0.63 | 0.008 | 1.06 | 0.310 |
| Lateral Frontal Orbitalis | 2.80 (0.15) | 2.79 (0.17) | 1.34 | 0.96 | 0.96 | <0.001 | -1.12 | 0.284 | 2.78 (0.08) | 2.81 (0.09) | 2.00 | 0.69 | 0.71 | 0.002 | 1.54 | 0.151 |
| Lateral Frontal Triangularis | 2.58 (0.12) | 2.59 (0.13) | 1.66 | 0.92 | 0.92 | <0.001 | 0.99 | 0.339 | 2.52 (0.12) | 2.52 (0.12) | 2.68 | 0.81 | 0.81 | <0.001 | 0.03 | 0.978 |
| Lateral Orbitofrontal | 2.77 (0.11) | 2.83 (0.14) | 3.02 | 0.73 | 0.80 | <0.001 | 2.71 | 0.019 | 2.74 (0.14) | 2.77 (0.15) | 3.10 | 0.75 | 0.75 | <0.001 | 0.98 | 0.345 |
| Lingual Gyrus | 2.15 (0.09) | 2.14 (0.09) | 1.95 | 0.86 | 0.86 | <0.001 | -1.21 | 0.251 | 2.14 (0.10) | 2.11 (0.09) | 2.15 | 0.86 | 0.89 | <0.001 | -2.18 | 0.050 |
| Medial Orbitofrontal | 2.45 (0.13) | 2.55 (0.13) | 5.36 | 0.47 | 0.61 | 0.011 | 3.34 | 0.006 | 2.39 (0.19) | 2.46 (0.16) | 5.00 | 0.62 | 0.67 | 0.005 | 1.90 | 0.081 |
| Middle Temporal | 2.92 (0.11) | 2.90 (0.15) | 1.82 | 0.89 | 0.89 | <0.001 | -1.01 | 0.332 | 2.95 (0.11) | 2.97 (0.13) | 1.57 | 0.91 | 0.92 | <0.001 | 1.85 | 0.089 |
| Paracentral Gyrus | 2.61 (0.17) | 2.59 (0.16) | 2.02 | 0.92 | 0.93 | <0.001 | -1.11 | 0.289 | 2.63 (0.10) | 2.61 (0.12) | 1.95 | 0.81 | 0.81 | <0.001 | -1.04 | 0.320 |
| Parahippocampal | 2.80 (0.29) | 2.77 (0.30) | 2.92 | 0.94 | 0.94 | <0.001 | -0.98 | 0.348 | 2.81 (0.29) | 2.78 (0.29) | 2.56 | 0.96 | 0.96 | <0.001 | -1.66 | 0.124 |
| Pericalcarine | 1.78 (0.13) | 1.71 (0.13) | 4.10 | 0.78 | 0.90 | <0.001 | -4.47 | <0.001* | 1.84 (0.08) | 1.73 (0.08) | 6.25 | 0.35 | 0.65 | 0.006 | -5.84 | <0.001* |
| Postcentral Gyrus | 2.18 (0.14) | 2.16 (0.14) | 1.27 | 0.97 | 0.98 | <0.001 | -2.08 | 0.060 | 2.18 (0.13) | 2.15 (0.12) | 1.50 | 0.95 | 0.96 | <0.001 | -2.66 | 0.021 |
| Posterior Cingulate | 2.57 (0.11) | 2.60 (0.13) | 2.05 | 0.85 | 0.86 | <0.001 | 1.55 | 0.148 | 2.56 (0.14) | 2.56 (0.16) | 1.53 | 0.94 | 0.94 | <0.001 | 0.07 | 0.941 |
| Precentral Gyrus | 2.80 (0.10) | 2.81 (0.11) | 1.04 | 0.95 | 0.95 | <0.001 | 0.50 | 0.624 | 2.76 (0.11) | 2.76 (0.11) | 1.40 | 0.92 | 0.92 | <0.001 | 0.05 | 0.962 |
| Precuneus | 2.49 (0.10) | 2.45 (0.10) | 1.83 | 0.85 | 0.92 | <0.001 | -3.62 | 0.004 | 2.54 (0.06) | 2.50 (0.07) | 1.94 | 0.58 | 0.66 | 0.005 | -2.49 | 0.029 |
| Rostral Anterior Cingulate | 2.86 (0.17) | 2.93 (0.18) | 3.38 | 0.82 | 0.87 | <0.001 | 2.79 | 0.016 | 2.78 (0.13) | 2.86 (0.14) | 4.22 | 0.43 | 0.49 | 0.038 | 2.08 | 0.060 |
| Rostral Middle Frontal | 2.55 (0.13) | 2.57 (0.13) | 1.53 | 0.94 | 0.95 | <0.001 | 2.24 | 0.045 | 2.40 (0.12) | 2.41 (0.10) | 3.32 | 0.64 | 0.64 | 0.007 | 0.26 | 0.802 |
| Superior Frontal Gyrus | 2.79 (0.12) | 2.82 (0.13) | 1.21 | 0.96 | 0.97 | <0.001 | 2.89 | 0.014 | 2.68 (0.12) | 2.70 (0.13) | 2.07 | 0.85 | 0.85 | <0.001 | 1.15 | 0.273 |
| Superior Parietal | 2.31 (0.10) | 2.25 (0.12) | 2.79 | 0.80 | 0.93 | <0.001 | -5.36 | <0.001* | 2.31 (0.10) | 2.26 (0.12) | 3.23 | 0.74 | 0.82 | <0.001 | -2.99 | 0.011 |
| Superior Temporal | 3.02 (0.10) | 3.01 (0.13) | 1.26 | 0.91 | 0.91 | <0.001 | -0.14 | 0.889 | 3.03 (0.10) | 3.04 (0.11) | 1.53 | 0.87 | 0.87 | <0.001 | 0.59 | 0.564 |
| Supramarginal Gyrus | 2.68 (0.16) | 2.68 (0.17) | 1.07 | 0.97 | 0.97 | <0.001 | -0.02 | 0.984 | 2.66 (0.16) | 2.66 (0.17) | 2.03 | 0.93 | 0.93 | <0.001 | -0.51 | 0.618 |
| Transverse Temporal | 2.54 (0.18) | 2.43 (0.21) | 6.07 | 0.53 | 0.59 | 0.013 | -2.20 | 0.048 | 2.54 (0.12) | 2.49 (0.19) | 3.67 | 0.73 | 0.75 | <0.001 | -1.64 | 0.126 |

APD, absolute percentage difference; ICC, intraclass correlation coefficient; lmer, linear mixed-effects model; ROI, region of interest; SD, standard deviation. Note: positive t-values represent cortical thickness increases.

**Supplementary Table 5.** Prisma-Prisma reliability measures for cortical thickness estimated by the CIVET pipeline.

| ROI | CIVET |  |  |  |  |  |  |  |  |  |  |  |  |  |  |  |
| --- | --- | --- | --- | --- | --- | --- | --- | --- | --- | --- | --- | --- | --- | --- | --- | --- |
|  | Left Hemisphere |  |  |  |  |  |  |  | Right Hemisphere |  |  |  |  |  |  |  |
|  | Timepoint 1<br>Mean (SD) | Timepoint 2<br>Mean (SD) | APD | ICC (2,1) | ICC (3,1) | p-value (ICC) | t-value (lmer) | p-value (lmer) | Timepoint 1<br>Mean (SD) | Timepoint 2<br>Mean (SD) | APD | ICC (2,1) | ICC (3,1) | p-value (ICC) | t-value (lmer) | p-value (lmer) |
| Caudal Anterior Cingulate | 2.88 (0.13) | 2.90 (0.12) | 2.12 | 0.79 | 0.80 | <0.001 | 0.77 | 0.452 | 2.75 (0.14) | 2.77 (0.13) | 2.00 | 0.81 | 0.81 | <0.001 | -0.27 | 0.787 |
| Caudal Middle Frontal | 3.08 (0.08) | 3.08 (0.09) | 0.80 | 0.91 | 0.91 | <0.001 | 0.82 | 0.426 | 3.04 (0.11) | 3.04 (0.10) | 0.61 | 0.96 | 0.96 | <0.001 | 0.68 | 0.507 |
| Cuneus | 2.53 (0.11) | 2.54 (0.12) | 0.58 | 0.98 | 0.98 | <0.001 | 1.27 | 0.225 | 2.59 (0.12) | 2.59 (0.12) | 0.95 | 0.97 | 0.97 | <0.001 | 0.02 | 0.988 |
| Entorhinal Cortex | 3.89 (0.25) | 3.92 (0.29) | 2.67 | 0.85 | 0.85 | <0.001 | 0.12 | 0.903 | 4.00 (0.23) | 4.03 (0.27) | 2.69 | 0.87 | 0.87 | <0.001 | 0.93 | 0.369 |
| Fusiform Gyrus | 3.14 (0.11) | 3.14 (0.11) | 0.98 | 0.94 | 0.94 | <0.001 | 1.16 | 0.264 | 3.19 (0.13) | 3.19 (0.14) | 1.33 | 0.93 | 0.93 | <0.001 | 0.02 | 0.982 |
| Inferior Occipital Cortex | 2.66 (0.09) | 2.68 (0.08) | 1.07 | 0.93 | 0.94 | <0.001 | 2.77 | 0.014 | 2.81 (0.08) | 2.81 (0.08) | 1.08 | 0.91 | 0.91 | <0.001 | 1.80 | 0.092 |
| Inferior Parietal | 2.93 (0.11) | 2.95 (0.10) | 1.00 | 0.95 | 0.95 | <0.001 | 0.97 | 0.348 | 3.06 (0.11) | 3.05 (0.11) | 0.92 | 0.95 | 0.95 | <0.001 | 0.96 | 0.354 |
| Inferior Temporal | 3.19 (0.12) | 3.20 (0.12) | 0.86 | 0.96 | 0.96 | <0.001 | 1.47 | 0.161 | 3.30 (0.13) | 3.30 (0.14) | 1.48 | 0.90 | 0.90 | <0.001 | 1.56 | 0.139 |
| Insula | 3.63 (0.19) | 3.70 (0.18) | 2.64 | 0.80 | 0.85 | <0.001 | 2.60 | 0.020 | 3.64 (0.19) | 3.68 (0.21) | 1.81 | 0.90 | 0.91 | <0.001 | -0.27 | 0.789 |
| Isthmus Cingulate Gyrus | 3.15 (0.20) | 3.16 (0.22) | 2.49 | 0.81 | 0.81 | <0.001 | -0.29 | 0.778 | 3.15 (0.17) | 3.13 (0.20) | 3.14 | 0.80 | 0.80 | <0.001 | -0.37 | 0.719 |
| Lateral Frontal Opercularis | 3.05 (0.10) | 3.06 (0.08) | 0.88 | 0.93 | 0.93 | <0.001 | 1.42 | 0.177 | 3.05 (0.11) | 3.04 (0.11) | 1.16 | 0.93 | 0.93 | <0.001 | 1.18 | 0.256 |
| Lateral Frontal Orbitalis | 3.14 (0.09) | 3.15 (0.10) | 1.20 | 0.89 | 0.90 | <0.001 | 0.36 | 0.726 | 3.17 (0.12) | 3.19 (0.13) | 1.03 | 0.94 | 0.94 | <0.001 | -1.24 | 0.234 |
| Lateral Frontal Triangularis | 2.98 (0.11) | 2.98 (0.11) | 1.16 | 0.91 | 0.91 | <0.001 | 0.51 | 0.617 | 2.95 (0.10) | 2.96 (0.10) | 1.05 | 0.93 | 0.93 | <0.001 | 0.58 | 0.572 |
| Lateral Orbitofrontal | 3.00 (0.09) | 3.01 (0.09) | 1.55 | 0.84 | 0.84 | <0.001 | 1.67 | 0.117 | 2.94 (0.10) | 2.95 (0.09) | 1.22 | 0.89 | 0.89 | <0.001 | 0.79 | 0.443 |
| Lingual Gyrus | 2.62 (0.09) | 2.63 (0.10) | 1.74 | 0.82 | 0.82 | <0.001 | 1.67 | 0.115 | 2.68 (0.10) | 2.65 (0.10) | 1.98 | 0.73 | 0.75 | <0.001 | 0.17 | 0.871 |
| Medial Orbitofrontal | 2.73 (0.12) | 2.74 (0.10) | 2.35 | 0.73 | 0.73 | <0.001 | 1.99 | 0.065 | 2.72 (0.10) | 2.73 (0.12) | 1.74 | 0.84 | 0.84 | <0.001 | -0.48 | 0.637 |
| Middle Temporal | 3.28 (0.11) | 3.30 (0.11) | 0.86 | 0.94 | 0.96 | <0.001 | 2.51 | 0.024 | 3.37 (0.12) | 3.36 (0.12) | 0.81 | 0.96 | 0.96 | <0.001 | 1.76 | 0.099 |
| Paracentral Gyrus | 2.95 (0.14) | 2.96 (0.13) | 1.01 | 0.96 | 0.96 | <0.001 | 0.37 | 0.714 | 3.00 (0.11) | 2.99 (0.10) | 1.21 | 0.92 | 0.92 | <0.001 | -0.80 | 0.438 |
| Parahippocampal | 2.80 (0.13) | 2.83 (0.13) | 2.07 | 0.78 | 0.79 | <0.001 | 1.84 | 0.085 | 2.79 (0.12) | 2.82 (0.15) | 2.66 | 0.75 | 0.76 | <0.001 | -0.47 | 0.645 |
| Pericalcarine | 2.40 (0.11) | 2.40 (0.11) | 1.31 | 0.93 | 0.93 | <0.001 | 0.65 | 0.523 | 2.49 (0.10) | 2.49 (0.10) | 2.13 | 0.81 | 0.81 | <0.001 | -1.71 | 0.108 |
| Postcentral Gyrus | 2.65 (0.13) | 2.66 (0.13) | 0.88 | 0.98 | 0.98 | <0.001 | 0.86 | 0.404 | 2.68 (0.10) | 2.68 (0.11) | 0.69 | 0.96 | 0.96 | <0.001 | -1.31 | 0.209 |
| Posterior Cingulate | 2.94 (0.12) | 3.00 (0.14) | 2.59 | 0.75 | 0.80 | <0.001 | 0.74 | 0.469 | 2.95 (0.07) | 2.96 (0.09) | 1.76 | 0.73 | 0.73 | <0.001 | -1.22 | 0.241 |
| Precentral Gyrus | 3.06 (0.09) | 3.07 (0.10) | 1.00 | 0.91 | 0.91 | <0.001 | 0.59 | 0.562 | 3.06 (0.11) | 3.06 (0.09) | 0.79 | 0.96 | 0.96 | <0.001 | 0.31 | 0.758 |
| Precuneus | 2.91 (0.08) | 2.93 (0.08) | 0.95 | 0.91 | 0.93 | <0.001 | 1.51 | 0.152 | 3.00 (0.1) | 3.01 (0.09) | 0.93 | 0.94 | 0.94 | <0.001 | 0.95 | 0.355 |
| Rostral Anterior Cingulate | 3.04 (0.19) | 3.11 (0.19) | 3.15 | 0.82 | 0.86 | <0.001 | 1.46 | 0.164 | 2.94 (0.14) | 2.98 (0.15) | 2.60 | 0.78 | 0.80 | <0.001 | 1.10 | 0.287 |
| Rostral Middle Frontal | 3.00 (0.11) | 3.01 (0.12) | 0.96 | 0.96 | 0.96 | <0.001 | 2.38 | 0.031 | 2.93 (0.11) | 2.92 (0.12) | 0.91 | 0.97 | 0.97 | <0.001 | 0.34 | 0.735 |
| Superior Frontal Gyrus | 3.14 (0.13) | 3.15 (0.13) | 1.01 | 0.94 | 0.94 | <0.001 | -0.38 | 0.706 | 3.07 (0.12) | 3.08 (0.13) | 0.93 | 0.95 | 0.95 | <0.001 | -0.62 | 0.543 |
| Superior Parietal | 2.71 (0.09) | 2.71 (0.10) | 0.83 | 0.96 | 0.96 | <0.001 | 0.90 | 0.380 | 2.75 (0.10) | 2.74 (0.10) | 0.55 | 0.98 | 0.98 | <0.001 | -0.78 | 0.446 |
| Superior Temporal | 3.30 (0.12) | 3.30 (0.11) | 0.94 | 0.95 | 0.95 | <0.001 | 2.14 | 0.049 | 3.34 (0.10) | 3.34 (0.10) | 0.92 | 0.94 | 0.94 | <0.001 | -0.41 | 0.689 |
| Supramarginal Gyrus | 3.03 (0.14) | 3.04 (0.13) | 0.98 | 0.96 | 0.96 | <0.001 | 2.52 | 0.024 | 3.05 (0.14) | 3.05 (0.14) | 0.66 | 0.98 | 0.98 | <0.001 | 0.79 | 0.444 |
| Transverse Temporal | 3.02 (0.14) | 3.01 (0.14) | 1.35 | 0.94 | 0.94 | <0.001 | 0.38 | 0.706 | 3.05 (0.17) | 3.04 (0.15) | 1.33 | 0.95 | 0.95 | <0.001 | 0.07 | 0.948 |

APD, absolute percentage difference; ICC, intraclass correlation coefficient; lmer, linear mixed-effects model; ROI, region of interest; SD, standard deviation. Note: positive t-values represent cortical thickness increases.

**Supplementary Table 6.** Prisma-Prisma reliability measures for cortical thickness estimated by the FreeSurfer pipeline.

|  | FreeSurfer |  |  |  |  |  |  |  |  |  |  |  |  |  |  |  |
| --- | --- | --- | --- | --- | --- | --- | --- | --- | --- | --- | --- | --- | --- | --- | --- | --- |
| ROI | Left Hemisphere |  |  |  |  |  |  |  | Right Hemisphere |  |  |  |  |  |  |  |
|  | Timepoint 1 Mean (SD) | Timepoint 2 Mean (SD) | APD | ICC (2,1) | ICC (3,1) | p-value (ICC) | t-value (lmer) | p-value (lmer) | Timepoint 1 Mean (SD) | Timepoint 2 Mean (SD) | APD | ICC (2,1) | ICC (3,1) | p-value (ICC) | t-value (lmer) | p-value (lmer) |
| Caudal Anterior Cingulate | 2.82 (0.15) | 2.82 (0.16) | 1.68 | 0.92 | 0.92 | <0.001 | -0.10 | 0.925 | 2.59 (0.18) | 2.60 (0.21) | 3.13 | 0.76 | 0.76 | <0.001 | 0.33 | 0.743 |
| Caudal Middle Frontal | 2.76 (0.09) | 2.77 (0.11) | 1.28 | 0.90 | 0.90 | <0.001 | 0.51 | 0.614 | 2.68 (0.13) | 2.67 (0.14) | 1.15 | 0.97 | 0.97 | <0.001 | -0.91 | 0.378 |
| Cuneus | 1.99 (0.06) | 2.01 (0.07) | 1.93 | 0.74 | 0.74 | <0.001 | 1.01 | 0.327 | 1.98 (0.13) | 1.98 (0.12) | 2.29 | 0.90 | 0.90 | <0.001 | -0.46 | 0.653 |
| Entorhinal Cortex | 3.54 (0.15) | 3.56 (0.23) | 3.70 | 0.67 | 0.67 | 0.001 | 0.39 | 0.704 | 3.69 (0.27) | 3.67 (0.24) | 3.25 | 0.82 | 0.82 | <0.001 | -0.44 | 0.668 |
| Fusiform Gyrus | 2.81 (0.12) | 2.81 (0.13) | 0.94 | 0.96 | 0.96 | <0.001 | -0.89 | 0.387 | 2.87 (0.11) | 2.86 (0.11) | 0.85 | 0.97 | 0.97 | <0.001 | -1.19 | 0.251 |
| Inferior Occipital Cortex | 2.26 (0.09) | 2.26 (0.09) | 1.55 | 0.87 | 0.87 | <0.001 | 0.30 | 0.772 | 2.38 (0.08) | 2.37 (0.09) | 1.31 | 0.89 | 0.89 | <0.001 | -0.84 | 0.415 |
| Inferior Parietal | 2.52 (0.13) | 2.51 (0.12) | 1.02 | 0.96 | 0.97 | <0.001 | -1.59 | 0.133 | 2.64 (0.10) | 2.64 (0.10) | 0.93 | 0.95 | 0.95 | <0.001 | 0.12 | 0.908 |
| Inferior Temporal | 2.95 (0.12) | 2.95 (0.14) | 1.55 | 0.85 | 0.85 | <0.001 | 0.39 | 0.702 | 2.97 (0.09) | 2.96 (0.10) | 1.24 | 0.90 | 0.90 | <0.001 | -0.57 | 0.575 |
| Insula | 3.25 (0.13) | 3.25 (0.11) | 1.44 | 0.88 | 0.88 | <0.001 | 0.23 | 0.825 | 3.26 (0.13) | 3.24 (0.13) | 1.07 | 0.95 | 0.95 | <0.001 | -1.21 | 0.247 |
| Isthmus Cingulate Gyrus | 2.60 (0.12) | 2.59 (0.16) | 2.08 | 0.89 | 0.89 | <0.001 | -0.89 | 0.385 | 2.54 (0.21) | 2.55 (0.18) | 3.02 | 0.85 | 0.85 | <0.001 | 0.16 | 0.872 |
| Lateral Frontal Opercularis | 2.81 (0.10) | 2.80 (0.10) | 1.18 | 0.92 | 0.93 | <0.001 | -1.03 | 0.319 | 2.68 (0.12) | 2.67 (0.12) | 1.38 | 0.93 | 0.93 | <0.001 | -0.77 | 0.451 |
| Lateral Frontal Orbitalis | 2.82 (0.17) | 2.85 (0.14) | 2.15 | 0.86 | 0.86 | <0.001 | 1.26 | 0.228 | 2.81 (0.11) | 2.82 (0.09) | 1.94 | 0.74 | 0.74 | <0.001 | 0.41 | 0.686 |
| Lateral Frontal Triangularis | 2.59 (0.12) | 2.57 (0.11) | 1.47 | 0.91 | 0.91 | <0.001 | -1.12 | 0.281 | 2.50 (0.12) | 2.49 (0.13) | 1.38 | 0.93 | 0.93 | <0.001 | -0.51 | 0.620 |
| Lateral Orbitofrontal | 2.86 (0.12) | 2.85 (0.10) | 1.81 | 0.84 | 0.84 | <0.001 | -0.79 | 0.440 | 2.78 (0.14) | 2.78 (0.15) | 1.63 | 0.93 | 0.93 | <0.001 | -0.05 | 0.961 |
| Lingual Gyrus | 2.14 (0.08) | 2.14 (0.08) | 2.02 | 0.80 | 0.80 | <0.001 | 0.06 | 0.950 | 2.12 (0.09) | 2.11 (0.10) | 1.76 | 0.88 | 0.88 | <0.001 | -0.72 | 0.486 |
| Medial Orbitofrontal | 2.58 (0.14) | 2.53 (0.19) | 4.30 | 0.71 | 0.74 | <0.001 | -1.79 | 0.094 | 2.49 (0.15) | 2.51 (0.15) | 4.43 | 0.61 | 0.61 | 0.005 | 0.59 | 0.563 |
| Middle Temporal | 2.91 (0.13) | 2.91 (0.14) | 0.79 | 0.98 | 0.98 | <0.001 | -0.06 | 0.955 | 3.00 (0.11) | 3.00 (0.13) | 1.08 | 0.95 | 0.95 | <0.001 | -0.33 | 0.746 |
| Paracentral Gyrus | 2.59 (0.19) | 2.59 (0.18) | 0.85 | 0.99 | 0.99 | <0.001 | -0.57 | 0.574 | 2.62 (0.13) | 2.60 (0.14) | 1.44 | 0.94 | 0.95 | <0.001 | -1.79 | 0.094 |
| Parahippocampal | 2.82 (0.27) | 2.82 (0.26) | 2.34 | 0.95 | 0.95 | <0.001 | 0.04 | 0.970 | 2.78 (0.27) | 2.75 (0.28) | 1.52 | 0.98 | 0.99 | <0.001 | -2.58 | 0.021 |
| Pericalcarine | 1.72 (0.14) | 1.74 (0.14) | 2.17 | 0.94 | 0.95 | <0.001 | 1.19 | 0.252 | 1.74 (0.11) | 1.75 (0.12) | 3.93 | 0.74 | 0.74 | <0.001 | 0.37 | 0.719 |
| Postcentral Gyrus | 2.19 (0.15) | 2.19 (0.15) | 1.11 | 0.98 | 0.98 | <0.001 | 0.03 | 0.974 | 2.17 (0.11) | 2.17 (0.12) | 1.00 | 0.98 | 0.98 | <0.001 | -0.55 | 0.591 |
| Posterior Cingulate | 2.58 (0.11) | 2.54 (0.12) | 2.19 | 0.82 | 0.87 | <0.001 | -2.80 | 0.013 | 2.59 (0.14) | 2.57 (0.13) | 2.03 | 0.87 | 0.87 | <0.001 | -1.27 | 0.223 |
| Precentral Gyrus | 2.80 (0.11) | 2.81 (0.13) | 0.84 | 0.97 | 0.97 | <0.001 | 0.97 | 0.347 | 2.76 (0.10) | 2.77 (0.11) | 1.00 | 0.95 | 0.95 | <0.001 | 0.10 | 0.924 |
| Precuneus | 2.45 (0.11) | 2.45 (0.09) | 1.00 | 0.95 | 0.95 | <0.001 | -0.44 | 0.669 | 2.50 (0.07) | 2.49 (0.07) | 1.07 | 0.90 | 0.90 | <0.001 | -0.52 | 0.607 |
| Rostral Anterior Cingulate | 2.91 (0.18) | 2.90 (0.18) | 2.03 | 0.90 | 0.90 | <0.001 | -0.69 | 0.499 | 2.86 (0.15) | 2.92 (0.13) | 3.55 | 0.67 | 0.70 | <0.001 | 2.02 | 0.061 |
| Rostral Middle Frontal | 2.58 (0.11) | 2.58 (0.13) | 1.14 | 0.95 | 0.95 | <0.001 | -0.34 | 0.736 | 2.44 (0.10) | 2.44 (0.11) | 1.79 | 0.87 | 0.87 | <0.001 | 0.18 | 0.863 |
| Superior Frontal Gyrus | 2.84 (0.12) | 2.84 (0.13) | 0.77 | 0.97 | 0.97 | <0.001 | -0.16 | 0.872 | 2.72 (0.12) | 2.72 (0.13) | 1.21 | 0.95 | 0.95 | <0.001 | 0.17 | 0.868 |
| Superior Parietal | 2.27 (0.12) | 2.27 (0.12) | 0.92 | 0.98 | 0.98 | <0.001 | 0.12 | 0.909 | 2.29 (0.13) | 2.29 (0.12) | 1.24 | 0.97 | 0.97 | <0.001 | 0.07 | 0.947 |
| Superior Temporal | 3.01 (0.14) | 3.01 (0.14) | 0.82 | 0.97 | 0.97 | <0.001 | -0.64 | 0.532 | 3.04 (0.10) | 3.04 (0.11) | 1.12 | 0.92 | 0.92 | <0.001 | -0.76 | 0.460 |
| Supramarginal Gyrus | 2.67 (0.16) | 2.66 (0.17) | 1.15 | 0.98 | 0.98 | <0.001 | -1.20 | 0.250 | 2.68 (0.15) | 2.67 (0.16) | 1.38 | 0.95 | 0.95 | <0.001 | -0.13 | 0.901 |
| Transverse Temporal | 2.46 (0.20) | 2.49 (0.23) | 2.67 | 0.92 | 0.93 | <0.001 | 1.74 | 0.102 | 2.52 (0.19) | 2.58 (0.17) | 3.48 | 0.81 | 0.85 | <0.001 | 2.56 | 0.022 |

APD, absolute percentage difference; ICC, intraclass correlation coefficient; lmer, linear mixed-effects model; ROI, region of interest; SD, standard deviation. Note: positive t-values represent cortical thickness increases.

**Supplementary Table 7.**
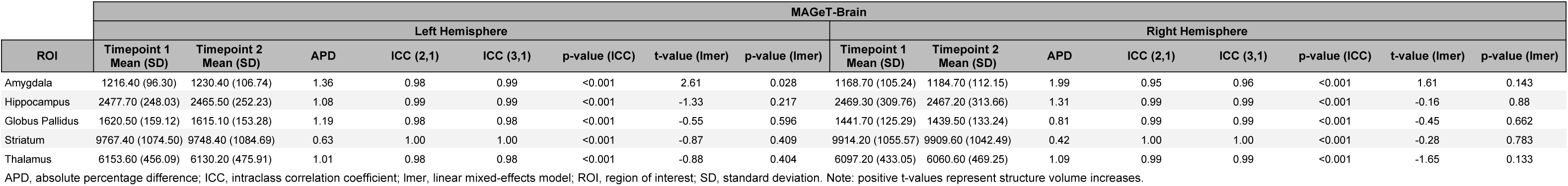
Trio-Trio reliability measures for structure volume estimated by the MAGeT-Brain pipeline.

**Supplementary Table 8.** Trio-Trio reliability measures for structure volume estimated by the FreeSurfer pipeline.

| ROI | FreeSurfer |  |  |  |  |  |  |  |  |  |  |  |  |  |  |  |
| --- | --- | --- | --- | --- | --- | --- | --- | --- | --- | --- | --- | --- | --- | --- | --- | --- |
|  | Left Hemisphere |  |  |  |  |  |  |  | Right Hemisphere |  |  |  |  |  |  |  |
|  | Timepoint 1<br>Mean (SD) | Timepoint 2<br>Mean (SD) | APD | ICC (2,1) | ICC (3,1) | p-value (ICC) | t-value (lmer) | p-value (lmer) | Timepoint 1<br>Mean (SD) | Timepoint 2<br>Mean (SD) | APD | ICC (2,1) | ICC (3,1) | p-value (ICC) | t-value (lmer) | p-value (lmer) |
| Amygdala | 1697.63 (210.87) | 1735.46 (241.32) | 3.39 | 0.95 | 0.96 | <0.001 | 1.84 | 0.100 | 1784.16 (219.38) | 1789.66 (255.86) | 3.04 | 0.97 | 0.97 | <0.001 | 0.27 | 0.795 |
| Hippocampus | 4249.92 (327.84) | 4239.38 (369.27) | 2.91 | 0.89 | 0.89 | <0.001 | -0.20 | 0.847 | 4346.86 (329.99) | 4385.21 (427.81) | 2.80 | 0.92 | 0.92 | <0.001 | 0.79 | 0.451 |
| Globus Pallidus | 2140.80 (256.85) | 2205.75 (211.29) | 4.22 | 0.90 | 0.93 | <0.001 | 2.29 | 0.048 | 1929.34 (186.41) | 2004.70 (192.24) | 5.09 | 0.80 | 0.85 | <0.001 | 2.31 | 0.046 |
| Striatum | 8503.25 (1120.80) | 8539.35 (1122.70) | 1.09 | 1.00 | 1.00 | <0.001 | 1.09 | 0.305 | 8640.18 (934.63) | 8637.54 (928.72) | 0.88 | 1.00 | 1.00 | <0.001 | -0.09 | 0.927 |
| Thalamus | 7567.50 (545.53) | 7606.80 (654.15) | 3.13 | 0.91 | 0.91 | <0.001 | 0.46 | 0.655 | 7480.53 (633.75) | 7509.11 (702.89) | 2.18 | 0.97 | 0.97 | <0.001 | 0.50 | 0.630 |

APD, absolute percentage difference; ICC, intraclass correlation coefficient; lmer, linear mixed-effects model; ROI, region of interest; SD, standard deviation. Note: positive t-values represent structure volume increases.

**Supplementary Table 9.** Trio-Prisma reliability measures for structure volume estimated by FreeSurfer.

| ROI | FreeSurfer |  |  |  |  |  |  |  |  |  |  |  |  |  |  |  |
| --- | --- | --- | --- | --- | --- | --- | --- | --- | --- | --- | --- | --- | --- | --- | --- | --- |
|  | Left Hemisphere |  |  |  |  |  |  |  | Right Hemisphere |  |  |  |  |  |  |  |
|  | Timepoint 1<br>Mean (SD) | Timepoint 2<br>Mean (SD) | APD | ICC (2,1) | ICC (3,1) | p-value (ICC) | t-value (lmer) | p-value (lmer) | Timepoint 1<br>Mean (SD) | Timepoint 2<br>Mean (SD) | APD | ICC (2,1) | ICC (3,1) | p-value (ICC) | t-value (lmer) | p-value (lmer) |
| Amygdala | 1718.35 (230.18) | 1477.37 (138.14) | 16.00 | 0.23 | 0.40 | 0.076 | -4.19 | 0.001* | 1774.99 (254.02) | 1625.55 (244.12) | 9.33 | 0.72 | 0.85 | <0.001 | -3.90 | 0.002* |
| Hippocampus | 4203.95 (329.73) | 4506.44 (653.09) | 9.31 | 0.20 | 0.22 | 0.226 | 1.69 | 0.118 | 4410.23 (399.21) | 4363.69 (355.78) | 4.47 | 0.75 | 0.75 | 0.001 | -0.61 | 0.553 |
| Globus Pallidus | 2193.55 (208.05) | 1608.02 (301.64) | 31.60 | 0.10 | 0.34 | 0.120 | -7.07 | <0.001* | 2001.1 (196.34) | 1653.24 (297.44) | 21.90 | 0.15 | 0.29 | 0.161 | -4.16 | 0.001* |
| Striatum | 8554.95 (1025.10) | 7505.94 (989.56) | 14.40 | 0.42 | 0.64 | 0.007 | -4.43 | <0.001* | 8716.21 (910.20) | 7586.56 (1006.60) | 14.70 | 0.42 | 0.70 | 0.003 | -5.48 | <0.001* |
| Thalamus | 7658.48 (589.68) | 8612.5 (1655.50) | 12.70 | 0.26 | 0.32 | 0.131 | 2.38 | 0.035 | 7569.81 (682.49) | 8354.95 (1634.80) | 11.40 | 0.46 | 0.53 | 0.025 | 2.34 | 0.038 |

APD, absolute percentage difference; ICC, intraclass correlation coefficient; lmer, linear mixed-effects model; ROI, region of interest; SD, standard deviation. Note: positive t-values represent structure volume increases. Asterisk indicates significance after Bonferroni correction.

**Supplementary Table 10.**
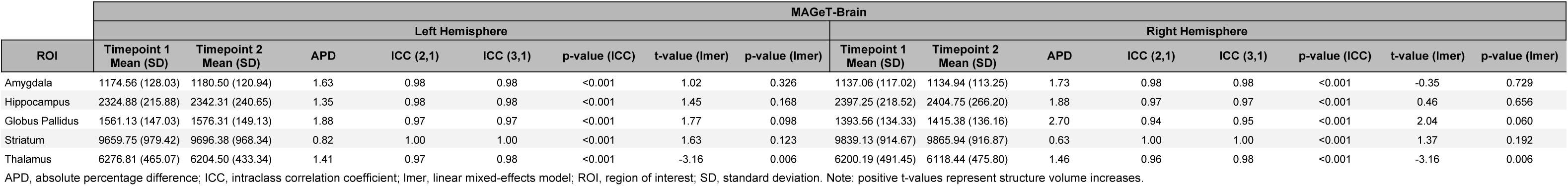
Prisma-Prisma reliability measures for structure volume estimated by the MAGeT-Brain pipeline.

**Supplementary Table 11.** Prisma-Prisma reliability measures for structure volume estimated by the FreeSurfer pipeline.

| ROI | FreeSurfer |  |  |  |  |  |  |  |  |  |  |  |  |  |  |  |
| --- | --- | --- | --- | --- | --- | --- | --- | --- | --- | --- | --- | --- | --- | --- | --- | --- |
|  | Left Hemisphere |  |  |  |  |  |  |  | Right Hemisphere |  |  |  |  |  |  |  |
|  | Timepoint 1<br>Mean (SD) | Timepoint 2<br>Mean (SD) | APD | ICC (2,1) | ICC (3,1) | p-value (ICC) | t-value (lmer) | p-value (lmer) | Timepoint 1<br>Mean (SD) | Timepoint 2<br>Mean (SD) | APD | ICC (2,1) | ICC (3,1) | p-value (ICC) | t-value (lmer) | p-value (lmer) |
| Amygdala | 1494.55 (146.39) | 1593.99 (182.17) | 7.42 | 0.56 | 0.64 | 0.003 | 2.85 | 0.012 | 1608.64 (240.83) | 1701.61 (238.54) | 7.66 | 0.74 | 0.78 | <0.001 | 2.36 | 0.032 |
| Hippocampus | 4249.69 (296.07) | 4248.63 (337.21) | 3.12 | 0.89 | 0.89 | <0.001 | -0.03 | 0.979 | 4287.03 (304.21) | 4334.63 (341.38) | 2.78 | 0.81 | 0.81 | <0.001 | 0.96 | 0.351 |
| Globus Pallidus | 1732.16 (327.95) | 1942.96 (315.51) | 12.30 | 0.45 | 0.53 | 0.014 | 2.71 | 0.016 | 1667.43 (236.73) | 1795.68 (214.92) | 11.10 | 0.34 | 0.39 | 0.063 | 2.05 | 0.059 |
| Striatum | 8035.04 (1077.1) | 8396.39 (1099.8) | 5.64 | 0.82 | 0.86 | <0.001 | 2.53 | 0.023 | 8066.73 (1157.8) | 8536.78 (938.11) | 6.77 | 0.73 | 0.79 | <0.001 | 2.75 | 0.015 |
| Thalamus | 7768.07 (732.64) | 7596.22 (809.57) | 5.15 | 0.77 | 0.78 | <0.001 | -1.33 | 0.202 | 7608.48 (817.59) | 7471.39 (708.92) | 3.95 | 0.80 | 0.80 | <0.001 | -1.14 | 0.273 |

APD, absolute percentage difference; ICC, intraclass correlation coefficient; lmer, linear mixed-effects model; ROI, region of interest; SD, standard deviation. Note: positive t-values represent structure volume increases.

**Supplementary Table 12.** Trio-Trio reliability measures for 1H-MRS indices.

| Measure | Timepoint 1<br>Mean (SD) | Timepoint 2<br>Mean (SD) | APD | ICC (2,1) | ICC (3,1) | p-value (ICC) | t-value (lmer) | p-value (lmer) |
| --- | --- | --- | --- | --- | --- | --- | --- | --- |
| Cho | 0.21 (0.02) | 0.21 (0.02) | 5.99 | 0.69 | 0.69 | 0.014 | 0.16 | 0.880 |
| GABA | 0.22 (0.05) | 0.22 (0.04) | 20.90 | 0.18 | 0.18 | 0.310 | 0.20 | 0.848 |
| Gln | 0.21 (0.03) | 0.24 (0.04) | 12.70 | 0.59 | 0.72 | 0.009 | 2.80 | 0.023 |
| Glu | 1.08 (0.14) | 1.07 (0.10) | 4.80 | 0.88 | 0.88 | <0.001 | -0.58 | 0.575 |
| Glx | 1.29 (0.15) | 1.30 (0.10) | 4.91 | 0.81 | 0.81 | 0.002 | 0.46 | 0.655 |
| GSH | 0.19 (0.01) | 0.19 (0.02) | 6.16 | 0.64 | 0.69 | 0.013 | 1.80 | 0.109 |
| Ins | 0.81 (0.06) | 0.79 (0.06) | 4.07 | 0.81 | 0.85 | 0.001 | -2.03 | 0.077 |
| Lac | 0.08 (0.02) | 0.07 (0.02) | 25.70 | 0.45 | 0.45 | 0.097 | -1.00 | 0.347 |
| NAA | 1.32 (0.13) | 1.33 (0.11) | 5.23 | 0.75 | 0.75 | 0.007 | 0.21 | 0.840 |
| NAAG | 0.12 (0.04) | 0.11 (0.03) | 26.70 | 0.07 | 0.07 | 0.421 | -0.80 | 0.447 |
| tNAA | 1.44 (0.13) | 1.44 (0.11) | 2.87 | 0.90 | 0.90 | <0.001 | -0.32 | 0.754 |
| FWHM | 7.26 (0.55) | 7.47 (0.90) | 11.40 | 0.10 | 0.10 | 0.395 | 0.60 | 0.564 |
| SNR | 332.8 (34.17) | 318.34 (61.19) | 12.40 | 0.57 | 0.57 | 0.043 | -0.94 | 0.375 |

APD, absolute percentage difference; ICC, intraclass correlation coefficient; lmer, linear mixed-effects model; ROI, region of interest; SD, standard deviation.
Note: positive t-values represent increases.

**Supplementary Table 13.** Prisma-Prisma reliability measures for 1H-MRS indices.

| Measure | Timepoint 1<br>Mean (SD) | Timepoint 2<br>Mean (SD) | APD | ICC (2,1) | ICC (3,1) | p-value (ICC) | t-value (lmer) | p-value (lmer) |
| --- | --- | --- | --- | --- | --- | --- | --- | --- |
| Cho | 0.22 (0.03) | 0.23 (0.02) | 5.59 | 0.80 | 0.80 | <0.001 | 1.00 | 0.335 |
| GABA | 0.25 (0.03) | 0.23 (0.06) | 14.40 | 0.54 | 0.55 | 0.022 | -1.09 | 0.296 |
| Gln | 0.23 (0.03) | 0.22 (0.04) | 10.10 | 0.75 | 0.75 | 0.001 | -0.71 | 0.492 |
| Glu | 1.13 (0.11) | 1.08 (0.12) | 6.18 | 0.79 | 0.85 | <0.001 | -2.63 | 0.022 |
| Glx | 1.35 (0.13) | 1.30 (0.15) | 6.09 | 0.78 | 0.82 | <0.001 | -2.30 | 0.040 |
| GSH | 0.20 (0.01) | 0.21 (0.01) | 5.64 | 0.29 | 0.32 | 0.136 | 1.60 | 0.135 |
| Ins | 0.86 (0.09) | 0.87 (0.09) | 5.57 | 0.80 | 0.80 | <0.001 | 0.42 | 0.684 |
| Lac | 0.07 (0.02) | 0.08 (0.03) | 27.20 | 0.35 | 0.35 | 0.108 | 1.01 | 0.331 |
| NAA | 1.35 (0.10) | 1.35 (0.11) | 3.79 | 0.81 | 0.81 | <0.001 | 0.12 | 0.908 |
| NAAG | 0.13 (0.03) | 0.14 (0.02) | 17.70 | 0.44 | 0.44 | 0.057 | 0.57 | 0.577 |
| tNAA | 1.48 (0.10) | 1.49 (0.12) | 2.74 | 0.85 | 0.85 | <0.001 | 0.39 | 0.701 |
| FWHM | 6.80 (0.39) | 6.85 (0.48) | 3.65 | 0.77 | 0.77 | <0.001 | 0.56 | 0.584 |
| SNR | 316.27 (50.28) | 317.52 (40.78) | 11.80 | 0.48 | 0.48 | 0.042 | 0.09 | 0.927 |

APD, absolute percentage difference; ICC, intraclass correlation coefficient; lmer, linear mixed-effects model; ROI, region of interest; SD, standard deviation.
Note: positive t-values represent increases.

**Supplementary Table 14.** MPRAGE reliability measures for cortical thickness estimated by the CIVET pipeline.

| ROI | CIVET |  |  |  |  |  |  |  |  |  |  |  |  |  |  |  |
| --- | --- | --- | --- | --- | --- | --- | --- | --- | --- | --- | --- | --- | --- | --- | --- | --- |
|  | Left Hemisphere |  |  |  |  |  |  |  | Right Hemisphere |  |  |  |  |  |  |  |
|  | Timepoint 1<br>Mean (SD) | Timepoint 2<br>Mean (SD) | APD | ICC (2,1) | ICC (3,1) | p-value (ICC) | t-value (lmer) | p-value (lmer) | Timepoint 1<br>Mean (SD) | Timepoint 2<br>Mean (SD) | APD | ICC (2,1) | ICC (3,1) | p-value (ICC) | t-value (lmer) | p-value (lmer) |
| Caudal Anterior Cingulate | 2.86 (0.14) | 2.86 (0.13) | 2.05 | 0.83 | 0.83 | <0.001 | -0.01 | 0.995 | 2.78 (0.13) | 2.78 (0.13) | 2.09 | 0.85 | 0.85 | <0.001 | 0.19 | 0.856 |
| Caudal Middle Frontal | 3.09 (0.07) | 3.09 (0.08) | 0.72 | 0.94 | 0.94 | <0.001 | -0.06 | 0.954 | 3.07 (0.09) | 3.06 (0.09) | 0.52 | 0.96 | 0.96 | <0.001 | -1.13 | 0.280 |
| Cuneus | 2.55 (0.10) | 2.56 (0.10) | 0.55 | 0.98 | 0.98 | <0.001 | 1.17 | 0.265 | 2.59 (0.12) | 2.60 (0.12) | 0.75 | 0.98 | 0.98 | <0.001 | 0.59 | 0.568 |
| Entorhinal Cortex | 4.00 (0.29) | 3.95 (0.24) | 2.49 | 0.86 | 0.87 | <0.001 | -1.42 | 0.182 | 4.07 (0.22) | 4.07 (0.25) | 2.09 | 0.89 | 0.89 | <0.001 | 0.03 | 0.973 |
| Fusiform Gyrus | 3.14 (0.09) | 3.14 (0.08) | 1.19 | 0.87 | 0.87 | <0.001 | 0.06 | 0.956 | 3.19 (0.11) | 3.19 (0.12) | 0.88 | 0.96 | 0.96 | <0.001 | 0.75 | 0.467 |
| Inferior Occipital Cortex | 2.69 (0.07) | 2.69 (0.07) | 0.87 | 0.93 | 0.93 | <0.001 | 0.72 | 0.485 | 2.81 (0.09) | 2.82 (0.08) | 0.74 | 0.95 | 0.95 | <0.001 | 1.44 | 0.176 |
| Inferior Parietal | 2.95 (0.10) | 2.96 (0.10) | 0.94 | 0.95 | 0.95 | <0.001 | 0.94 | 0.368 | 3.07 (0.10) | 3.07 (0.10) | 0.73 | 0.96 | 0.96 | <0.001 | 0.65 | 0.529 |
| Inferior Temporal | 3.19 (0.12) | 3.19 (0.12) | 1.01 | 0.93 | 0.93 | <0.001 | 0.11 | 0.917 | 3.28 (0.12) | 3.30 (0.12) | 1.10 | 0.93 | 0.94 | <0.001 | 1.84 | 0.090 |
| Insula | 3.67 (0.15) | 3.72 (0.16) | 2.12 | 0.82 | 0.83 | <0.001 | 1.66 | 0.123 | 3.67 (0.17) | 3.69 (0.17) | 1.72 | 0.90 | 0.91 | <0.001 | 1.11 | 0.288 |
| Isthmus Cingulate Gyrus | 3.16 (0.18) | 3.17 (0.21) | 2.07 | 0.90 | 0.90 | <0.001 | 0.46 | 0.656 | 3.10 (0.13) | 3.13 (0.13) | 2.43 | 0.74 | 0.74 | 0.001 | 0.91 | 0.380 |
| Lateral Frontal Opercularis | 3.07 (0.10) | 3.07 (0.09) | 0.91 | 0.93 | 0.93 | <0.001 | -0.36 | 0.724 | 3.06 (0.11) | 3.05 (0.13) | 1.19 | 0.94 | 0.94 | <0.001 | -0.53 | 0.603 |
| Lateral Frontal Orbitalis | 3.15 (0.08) | 3.16 (0.08) | 1.34 | 0.80 | 0.80 | <0.001 | 0.14 | 0.895 | 3.16 (0.14) | 3.17 (0.12) | 1.75 | 0.86 | 0.86 | <0.001 | 0.53 | 0.605 |
| Lateral Frontal Triangularis | 2.97 (0.08) | 2.97 (0.08) | 0.88 | 0.91 | 0.91 | <0.001 | 0.01 | 0.990 | 2.97 (0.10) | 2.97 (0.08) | 1.27 | 0.89 | 0.89 | <0.001 | 0.39 | 0.706 |
| Lateral Orbitofrontal | 3.02 (0.07) | 2.99 (0.06) | 2.05 | 0.48 | 0.53 | 0.025 | -2.09 | 0.058 | 2.95 (0.08) | 2.94 (0.08) | 1.45 | 0.76 | 0.76 | <0.001 | -1.04 | 0.317 |
| Lingual Gyrus | 2.63 (0.09) | 2.63 (0.07) | 1.96 | 0.70 | 0.70 | 0.003 | -0.08 | 0.938 | 2.66 (0.09) | 2.66 (0.09) | 0.97 | 0.93 | 0.93 | <0.001 | 0.37 | 0.717 |
| Medial Orbitofrontal | 2.73 (0.12) | 2.74 (0.10) | 1.88 | 0.85 | 0.85 | <0.001 | 0.62 | 0.546 | 2.72 (0.11) | 2.71 (0.10) | 1.98 | 0.84 | 0.84 | <0.001 | -0.13 | 0.898 |
| Middle Temporal | 3.28 (0.11) | 3.29 (0.10) | 1.02 | 0.92 | 0.92 | <0.001 | 0.59 | 0.566 | 3.37 (0.12) | 3.37 (0.12) | 0.66 | 0.98 | 0.98 | <0.001 | 0.56 | 0.589 |
| Paracentral Gyrus | 2.96 (0.13) | 2.97 (0.15) | 0.80 | 0.96 | 0.96 | <0.001 | 0.73 | 0.478 | 3.00 (0.10) | 2.99 (0.10) | 1.08 | 0.93 | 0.93 | <0.001 | -0.91 | 0.383 |
| Parahippocampal | 2.83 (0.11) | 2.83 (0.10) | 1.85 | 0.83 | 0.83 | <0.001 | -0.43 | 0.677 | 2.83 (0.15) | 2.81 (0.15) | 1.48 | 0.93 | 0.93 | <0.001 | -0.99 | 0.344 |
| Pericalcarine | 2.40 (0.12) | 2.42 (0.11) | 1.22 | 0.94 | 0.95 | <0.001 | 2.58 | 0.024 | 2.49 (0.10) | 2.49 (0.09) | 1.47 | 0.87 | 0.87 | <0.001 | 0.07 | 0.943 |
| Postcentral Gyrus | 2.69 (0.11) | 2.69 (0.12) | 0.80 | 0.97 | 0.97 | <0.001 | 0.78 | 0.452 | 2.69 (0.08) | 2.70 (0.10) | 0.54 | 0.98 | 0.98 | <0.001 | 1.33 | 0.207 |
| Posterior Cingulate | 2.96 (0.12) | 2.98 (0.14) | 2.33 | 0.82 | 0.82 | <0.001 | 0.76 | 0.461 | 2.96 (0.07) | 2.96 (0.08) | 1.76 | 0.64 | 0.64 | 0.007 | -0.39 | 0.704 |
| Precentral Gyrus | 3.08 (0.08) | 3.07 (0.09) | 0.80 | 0.94 | 0.94 | <0.001 | -0.29 | 0.778 | 3.09 (0.08) | 3.09 (0.08) | 0.61 | 0.95 | 0.95 | <0.001 | 1.02 | 0.326 |
| Precuneus | 2.93 (0.07) | 2.94 (0.08) | 0.99 | 0.89 | 0.89 | <0.001 | 1.05 | 0.316 | 3.02 (0.07) | 3.02 (0.09) | 0.79 | 0.93 | 0.93 | <0.001 | -0.18 | 0.856 |
| Rostral Anterior Cingulate | 3.06 (0.23) | 3.09 (0.16) | 3.55 | 0.80 | 0.80 | <0.001 | 0.74 | 0.471 | 2.95 (0.15) | 2.95 (0.15) | 1.90 | 0.89 | 0.89 | <0.001 | 0.23 | 0.819 |
| Rostral Middle Frontal | 3.02 (0.10) | 3.02 (0.09) | 0.70 | 0.94 | 0.94 | <0.001 | -0.35 | 0.733 | 2.95 (0.09) | 2.94 (0.09) | 0.75 | 0.96 | 0.96 | <0.001 | -1.54 | 0.149 |
| Superior Frontal Gyrus | 3.16 (0.11) | 3.16 (0.12) | 0.70 | 0.97 | 0.97 | <0.001 | -0.11 | 0.912 | 3.09 (0.12) | 3.08 (0.11) | 0.72 | 0.98 | 0.98 | <0.001 | -1.75 | 0.106 |
| Superior Parietal | 2.72 (0.09) | 2.72 (0.09) | 0.88 | 0.95 | 0.95 | <0.001 | -1.04 | 0.318 | 2.77 (0.08) | 2.77 (0.08) | 0.68 | 0.96 | 0.96 | <0.001 | -0.07 | 0.944 |
| Superior Temporal | 3.30 (0.11) | 3.30 (0.12) | 0.75 | 0.96 | 0.96 | <0.001 | 0.52 | 0.611 | 3.35 (0.10) | 3.35 (0.10) | 0.95 | 0.91 | 0.91 | <0.001 | -0.04 | 0.967 |
| Supramarginal Gyrus | 3.06 (0.11) | 3.05 (0.11) | 0.89 | 0.95 | 0.95 | <0.001 | -0.27 | 0.793 | 3.09 (0.12) | 3.08 (0.11) | 0.59 | 0.98 | 0.98 | <0.001 | -0.48 | 0.642 |
| Transverse Temporal | 3.00 (0.15) | 3.03 (0.16) | 1.40 | 0.95 | 0.96 | <0.001 | 2.47 | 0.029 | 3.06 (0.15) | 3.08 (0.16) | 1.22 | 0.95 | 0.96 | <0.001 | 1.32 | 0.211 |

APD, absolute percentage difference; ICC, intraclass correlation coefficient; lmer, linear mixed-effects model; ROI, region of interest; SD, standard deviation. Note: positive t-values represent cortical thickness increases.

**Supplementary Table 15.** MPRAGE reliability measures for cortical thickness estimated by the FreeSurfer pipeline.

| ROI | FreeSurfer |  |  |  |  |  |  |  |  |  |  |  |  |  |  |  |
| --- | --- | --- | --- | --- | --- | --- | --- | --- | --- | --- | --- | --- | --- | --- | --- | --- |
|  | Left Hemisphere |  |  |  |  |  |  |  | Right Hemisphere |  |  |  |  |  |  |  |
|  | Timepoint 1<br>Mean (SD) | Timepoint 2<br>Mean (SD) | APD | ICC (2,1) | ICC (3,1) | p-value (ICC) | t-value (lmer) | p-value (lmer) | Timepoint 1<br>Mean (SD) | Timepoint 2<br>Mean (SD) | APD | ICC (2,1) | ICC (3,1) | p-value (ICC) | t-value (lmer) | p-value (lmer) |
| Caudal Anterior Cingulate | 2.76 (0.12) | 2.78 (0.15) | 2.30 | 0.82 | 0.82 | <0.001 | 1.16 | 0.267 | 2.58 (0.17) | 2.60 (0.20) | 2.18 | 0.91 | 0.91 | <0.001 | 0.64 | 0.533 |
| Caudal Middle Frontal | 2.75 (0.08) | 2.76 (0.11) | 1.14 | 0.90 | 0.90 | <0.001 | 0.75 | 0.468 | 2.69 (0.11) | 2.70 (0.13) | 1.25 | 0.95 | 0.95 | <0.001 | 0.56 | 0.588 |
| Cuneus | 2.00 (0.06) | 2.02 (0.07) | 2.11 | 0.74 | 0.75 | <0.001 | 1.44 | 0.177 | 2.00 (0.13) | 1.98 (0.12) | 1.98 | 0.92 | 0.93 | <0.001 | -1.52 | 0.154 |
| Entorhinal Cortex | 3.55 (0.26) | 3.57 (0.21) | 3.68 | 0.73 | 0.73 | 0.002 | 0.40 | 0.694 | 3.68 (0.20) | 3.74 (0.21) | 3.53 | 0.75 | 0.76 | <0.001 | 1.36 | 0.199 |
| Fusiform Gyrus | 2.82 (0.11) | 2.82 (0.11) | 1.22 | 0.91 | 0.91 | <0.001 | 0.47 | 0.650 | 2.88 (0.12) | 2.87 (0.12) | 0.96 | 0.97 | 0.97 | <0.001 | -1.22 | 0.247 |
| Inferior Occipital Cortex | 2.26 (0.07) | 2.28 (0.08) | 1.53 | 0.85 | 0.87 | <0.001 | 1.70 | 0.116 | 2.38 (0.08) | 2.39 (0.09) | 1.23 | 0.88 | 0.88 | <0.001 | 0.32 | 0.758 |
| Inferior Parietal | 2.52 (0.10) | 2.52 (0.12) | 1.35 | 0.93 | 0.93 | <0.001 | 0.18 | 0.858 | 2.66 (0.10) | 2.67 (0.11) | 1.17 | 0.94 | 0.94 | <0.001 | 1.20 | 0.252 |
| Inferior Temporal | 2.94 (0.12) | 2.94 (0.13) | 1.26 | 0.94 | 0.94 | <0.001 | 0.43 | 0.677 | 2.94 (0.09) | 2.95 (0.08) | 1.40 | 0.82 | 0.82 | <0.001 | 0.85 | 0.412 |
| Insula | 3.25 (0.08) | 3.26 (0.08) | 1.00 | 0.89 | 0.90 | <0.001 | 1.46 | 0.169 | 3.25 (0.11) | 3.24 (0.09) | 1.21 | 0.90 | 0.90 | <0.001 | -0.45 | 0.659 |
| Isthmus Cingulate Gyrus | 2.60 (0.14) | 2.60 (0.15) | 2.01 | 0.90 | 0.90 | <0.001 | -0.10 | 0.919 | 2.57 (0.18) | 2.56 (0.20) | 3.06 | 0.85 | 0.85 | <0.001 | -0.14 | 0.892 |
| Lateral Frontal Opercularis | 2.80 (0.09) | 2.80 (0.10) | 1.01 | 0.94 | 0.94 | <0.001 | -0.78 | 0.451 | 2.68 (0.13) | 2.69 (0.12) | 1.66 | 0.91 | 0.91 | <0.001 | 0.35 | 0.734 |
| Lateral Frontal Orbitalis | 2.87 (0.14) | 2.85 (0.13) | 1.92 | 0.86 | 0.86 | <0.001 | -0.98 | 0.347 | 2.80 (0.12) | 2.83 (0.11) | 2.57 | 0.72 | 0.72 | 0.002 | 1.19 | 0.257 |
| Lateral Frontal Triangularis | 2.56 (0.10) | 2.56 (0.10) | 1.23 | 0.92 | 0.92 | <0.001 | -0.07 | 0.944 | 2.49 (0.11) | 2.50 (0.11) | 2.05 | 0.84 | 0.84 | <0.001 | 0.77 | 0.455 |
| Lateral Orbitofrontal | 2.86 (0.11) | 2.84 (0.10) | 1.99 | 0.77 | 0.78 | <0.001 | -1.20 | 0.253 | 2.76 (0.12) | 2.79 (0.10) | 2.01 | 0.82 | 0.83 | <0.001 | 1.37 | 0.196 |
| Lingual Gyrus | 2.14 (0.09) | 2.14 (0.08) | 1.32 | 0.93 | 0.93 | <0.001 | 0.17 | 0.870 | 2.12 (0.10) | 2.12 (0.10) | 1.61 | 0.89 | 0.89 | <0.001 | 0.20 | 0.848 |
| Medial Orbitofrontal | 2.49 (0.21) | 2.51 (0.19) | 4.94 | 0.72 | 0.72 | 0.002 | 0.46 | 0.652 | 2.40 (0.18) | 2.40 (0.18) | 3.53 | 0.78 | 0.78 | <0.001 | 0.15 | 0.881 |
| Middle Temporal | 2.90 (0.12) | 2.91 (0.11) | 1.08 | 0.95 | 0.95 | <0.001 | 0.50 | 0.628 | 2.99 (0.12) | 2.99 (0.12) | 1.29 | 0.93 | 0.93 | <0.001 | 0.42 | 0.681 |
| Paracentral Gyrus | 2.63 (0.19) | 2.61 (0.20) | 0.96 | 0.99 | 0.99 | <0.001 | -1.26 | 0.233 | 2.62 (0.13) | 2.61 (0.14) | 1.12 | 0.96 | 0.97 | <0.001 | -1.62 | 0.130 |
| Parahippocampal | 2.84 (0.23) | 2.82 (0.26) | 1.57 | 0.98 | 0.98 | <0.001 | -1.40 | 0.188 | 2.77 (0.28) | 2.74 (0.27) | 1.47 | 0.98 | 0.98 | <0.001 | -1.82 | 0.094 |
| Pericalcarine | 1.73 (0.12) | 1.73 (0.15) | 2.09 | 0.94 | 0.94 | <0.001 | 0.25 | 0.803 | 1.76 (0.11) | 1.76 (0.11) | 2.57 | 0.84 | 0.84 | <0.001 | -0.40 | 0.700 |
| Postcentral Gyrus | 2.22 (0.13) | 2.22 (0.14) | 1.22 | 0.97 | 0.97 | <0.001 | -0.04 | 0.966 | 2.20 (0.10) | 2.20 (0.12) | 1.16 | 0.95 | 0.95 | <0.001 | 0.10 | 0.919 |
| Posterior Cingulate | 2.57 (0.13) | 2.56 (0.14) | 1.73 | 0.91 | 0.91 | <0.001 | -1.03 | 0.324 | 2.59 (0.14) | 2.56 (0.11) | 2.57 | 0.78 | 0.79 | <0.001 | -1.34 | 0.206 |
| Precentral Gyrus | 2.81 (0.11) | 2.81 (0.12) | 0.54 | 0.99 | 0.99 | <0.001 | 0.03 | 0.977 | 2.78 (0.10) | 2.78 (0.11) | 0.98 | 0.94 | 0.94 | <0.001 | -0.48 | 0.640 |
| Precuneus | 2.44 (0.10) | 2.46 (0.10) | 1.03 | 0.93 | 0.95 | <0.001 | 2.30 | 0.040 | 2.50 (0.08) | 2.50 (0.08) | 1.24 | 0.90 | 0.90 | <0.001 | 0.17 | 0.867 |
| Rostral Anterior Cingulate | 2.88 (0.17) | 2.87 (0.16) | 2.65 | 0.78 | 0.78 | <0.001 | -0.06 | 0.952 | 2.87 (0.11) | 2.89 (0.13) | 1.95 | 0.85 | 0.85 | <0.001 | 1.04 | 0.318 |
| Rostral Middle Frontal | 2.60 (0.09) | 2.59 (0.08) | 0.65 | 0.97 | 0.98 | <0.001 | -1.43 | 0.179 | 2.44 (0.09) | 2.45 (0.09) | 1.08 | 0.94 | 0.94 | <0.001 | 0.82 | 0.430 |
| Superior Frontal Gyrus | 2.85 (0.10) | 2.84 (0.11) | 0.91 | 0.95 | 0.95 | <0.001 | -0.70 | 0.496 | 2.72 (0.11) | 2.72 (0.12) | 1.08 | 0.94 | 0.94 | <0.001 | -0.34 | 0.738 |
| Superior Parietal | 2.27 (0.11) | 2.27 (0.12) | 1.13 | 0.97 | 0.97 | <0.001 | 0.55 | 0.594 | 2.31 (0.13) | 2.31 (0.13) | 0.92 | 0.98 | 0.98 | <0.001 | 1.29 | 0.221 |
| Superior Temporal | 3.00 (0.13) | 3.01 (0.13) | 0.90 | 0.97 | 0.97 | <0.001 | 0.31 | 0.766 | 3.05 (0.10) | 3.03 (0.10) | 1.26 | 0.90 | 0.90 | <0.001 | -1.50 | 0.161 |
| Supramarginal Gyrus | 2.69 (0.15) | 2.68 (0.15) | 0.82 | 0.99 | 0.99 | <0.001 | -0.87 | 0.400 | 2.70 (0.13) | 2.71 (0.15) | 1.23 | 0.96 | 0.96 | <0.001 | 0.35 | 0.730 |
| Transverse Temporal | 2.45 (0.23) | 2.49 (0.26) | 2.16 | 0.96 | 0.97 | <0.001 | 2.92 | 0.013 | 2.56 (0.22) | 2.57 (0.19) | 2.29 | 0.94 | 0.94 | <0.001 | 0.60 | 0.559 |

APD, absolute percentage difference; ICC, intraclass correlation coefficient; lmer, linear mixed-effects model; ROI, region of interest; SD, standard deviation. Note: positive t-values represent cortical thickness increases.

**Supplementary Table 16.** vNav-MPRAGE reliability measures for cortical thickness estimated by the CIVET pipeline.

| ROI | CIVET |  |  |  |  |  |  |  |  |  |  |  |  |  |  |  |
| --- | --- | --- | --- | --- | --- | --- | --- | --- | --- | --- | --- | --- | --- | --- | --- | --- |
|  | Left Hemisphere |  |  |  |  |  |  |  | Right Hemisphere |  |  |  |  |  |  |  |
|  | Timepoint 1<br>Mean (SD) | Timepoint 2<br>Mean (SD) | APD | ICC (2,1) | ICC (3,1) | p-value (ICC) | t-value (lmer) | p-value (lmer) | Timepoint 1<br>Mean (SD) | Timepoint 2<br>Mean (SD) | APD | ICC (2,1) | ICC (3,1) | p-value (ICC) | t-value (lmer) | p-value (lmer) |
| Caudal Anterior Cingulate | 2.80 (0.11) | 2.79 (0.12) | 2.60 | 0.75 | 0.75 | 0.001 | -0.48 | 0.640 | 2.74 (0.14) | 2.74 (0.15) | 1.85 | 0.91 | 0.91 | <0.001 | 0.41 | 0.690 |
| Caudal Middle Frontal | 3.06 (0.07) | 3.06 (0.09) | 1.07 | 0.89 | 0.89 | <0.001 | 0.01 | 0.989 | 3.05 (0.08) | 3.05 (0.09) | 0.72 | 0.95 | 0.95 | <0.001 | 1.20 | 0.252 |
| Cuneus | 2.55 (0.11) | 2.57 (0.09) | 1.32 | 0.91 | 0.92 | <0.001 | 1.69 | 0.116 | 2.58 (0.10) | 2.57 (0.10) | 1.33 | 0.89 | 0.89 | <0.001 | -0.26 | 0.800 |
| Entorhinal Cortex | 3.83 (0.27) | 3.90 (0.26) | 3.09 | 0.84 | 0.86 | <0.001 | 1.70 | 0.114 | 3.98 (0.19) | 4.04 (0.16) | 2.69 | 0.62 | 0.64 | 0.007 | 1.50 | 0.159 |
| Fusiform Gyrus | 3.11 (0.08) | 3.11 (0.11) | 1.28 | 0.85 | 0.85 | <0.001 | 0.07 | 0.944 | 3.14 (0.10) | 3.15 (0.11) | 1.15 | 0.92 | 0.92 | <0.001 | 0.96 | 0.356 |
| Inferior Occipital Cortex | 2.68 (0.07) | 2.68 (0.07) | 1.36 | 0.79 | 0.79 | <0.001 | -0.02 | 0.986 | 2.78 (0.08) | 2.79 (0.09) | 1.32 | 0.89 | 0.89 | <0.001 | 0.97 | 0.350 |
| Inferior Parietal | 2.96 (0.09) | 2.96 (0.11) | 0.90 | 0.95 | 0.95 | <0.001 | 0.20 | 0.846 | 3.02 (0.09) | 3.03 (0.10) | 0.92 | 0.94 | 0.94 | <0.001 | 1.44 | 0.176 |
| Inferior Temporal | 3.20 (0.10) | 3.20 (0.11) | 1.15 | 0.87 | 0.87 | <0.001 | -0.22 | 0.833 | 3.23 (0.08) | 3.26 (0.11) | 1.68 | 0.77 | 0.79 | <0.001 | 1.69 | 0.117 |
| Insula | 3.61 (0.15) | 3.61 (0.14) | 1.84 | 0.84 | 0.84 | <0.001 | 0.05 | 0.965 | 3.60 (0.13) | 3.55 (0.18) | 2.50 | 0.72 | 0.74 | 0.001 | -1.67 | 0.120 |
| Isthmus Cingulate Gyrus | 3.04 (0.23) | 3.05 (0.20) | 2.80 | 0.86 | 0.86 | <0.001 | 0.19 | 0.851 | 2.97 (0.17) | 2.99 (0.17) | 4.44 | 0.48 | 0.48 | 0.042 | 0.52 | 0.611 |
| Lateral Frontal Opercularis | 3.06 (0.08) | 3.07 (0.08) | 0.60 | 0.95 | 0.95 | <0.001 | 0.19 | 0.850 | 3.04 (0.10) | 3.07 (0.11) | 0.91 | 0.94 | 0.97 | <0.001 | 3.38 | 0.005 |
| Lateral Frontal Orbitalis | 3.13 (0.09) | 3.11 (0.05) | 1.68 | 0.52 | 0.54 | 0.023 | -1.38 | 0.192 | 3.12 (0.12) | 3.12 (0.13) | 1.54 | 0.91 | 0.91 | <0.001 | -0.08 | 0.939 |
| Lateral Frontal Triangularis | 2.97 (0.08) | 2.96 (0.10) | 0.92 | 0.94 | 0.94 | <0.001 | -0.99 | 0.344 | 2.96 (0.09) | 2.97 (0.10) | 1.04 | 0.94 | 0.94 | <0.001 | 0.53 | 0.607 |
| Lateral Orbitofrontal | 2.98 (0.07) | 3.01 (0.06) | 1.68 | 0.59 | 0.62 | 0.009 | 1.64 | 0.126 | 2.92 (0.09) | 2.93 (0.08) | 1.95 | 0.66 | 0.66 | 0.005 | 0.35 | 0.730 |
| Lingual Gyrus | 2.59 (0.08) | 2.61 (0.07) | 1.50 | 0.79 | 0.81 | <0.001 | 1.59 | 0.137 | 2.61 (0.06) | 2.61 (0.09) | 1.89 | 0.75 | 0.75 | <0.001 | 0.23 | 0.823 |
| Medial Orbitofrontal | 2.78 (0.11) | 2.80 (0.10) | 2.21 | 0.71 | 0.72 | 0.002 | 1.31 | 0.214 | 2.77 (0.11) | 2.78 (0.12) | 1.43 | 0.88 | 0.88 | <0.001 | 0.16 | 0.876 |
| Middle Temporal | 3.26 (0.10) | 3.26 (0.11) | 1.28 | 0.88 | 0.88 | <0.001 | -0.30 | 0.768 | 3.32 (0.11) | 3.32 (0.13) | 0.75 | 0.96 | 0.96 | <0.001 | 0.77 | 0.458 |
| Paracentral Gyrus | 2.96 (0.14) | 2.96 (0.14) | 1.34 | 0.91 | 0.91 | <0.001 | -0.09 | 0.929 | 2.93 (0.10) | 2.95 (0.12) | 1.48 | 0.88 | 0.89 | <0.001 | 1.46 | 0.171 |
| Parahippocampal | 2.72 (0.14) | 2.72 (0.12) | 1.60 | 0.92 | 0.92 | <0.001 | -0.13 | 0.900 | 2.71 (0.12) | 2.73 (0.18) | 2.35 | 0.88 | 0.88 | <0.001 | 1.09 | 0.298 |
| Pericalcarine | 2.42 (0.11) | 2.42 (0.13) | 1.69 | 0.91 | 0.91 | <0.001 | 0.20 | 0.849 | 2.48 (0.08) | 2.48 (0.10) | 2.93 | 0.53 | 0.53 | 0.027 | -0.03 | 0.978 |
| Postcentral Gyrus | 2.71 (0.11) | 2.72 (0.11) | 0.79 | 0.97 | 0.97 | <0.001 | 0.58 | 0.572 | 2.71 (0.09) | 2.71 (0.10) | 0.79 | 0.97 | 0.97 | <0.001 | 1.07 | 0.308 |
| Posterior Cingulate | 2.94 (0.11) | 2.94 (0.10) | 0.98 | 0.95 | 0.95 | <0.001 | -0.32 | 0.754 | 2.89 (0.06) | 2.90 (0.05) | 1.40 | 0.51 | 0.51 | 0.030 | 0.70 | 0.499 |
| Precentral Gyrus | 3.05 (0.07) | 3.05 (0.09) | 0.85 | 0.92 | 0.92 | <0.001 | -0.08 | 0.935 | 3.05 (0.08) | 3.05 (0.09) | 0.82 | 0.95 | 0.95 | <0.001 | 0.04 | 0.971 |
| Precuneus | 2.92 (0.07) | 2.92 (0.08) | 0.69 | 0.94 | 0.94 | <0.001 | 0.03 | 0.979 | 2.96 (0.06) | 2.97 (0.07) | 0.96 | 0.82 | 0.82 | <0.001 | 0.99 | 0.340 |
| Rostral Anterior Cingulate | 3.05 (0.17) | 3.04 (0.19) | 3.03 | 0.82 | 0.82 | <0.001 | -0.17 | 0.865 | 2.91 (0.15) | 2.93 (0.16) | 2.55 | 0.83 | 0.83 | <0.001 | 0.78 | 0.452 |
| Rostral Middle Frontal | 3.01 (0.10) | 3.01 (0.10) | 0.73 | 0.95 | 0.95 | <0.001 | -0.19 | 0.849 | 2.97 (0.08) | 2.98 (0.09) | 0.88 | 0.93 | 0.93 | <0.001 | 0.93 | 0.368 |
| Superior Frontal Gyrus | 3.17 (0.11) | 3.16 (0.12) | 0.94 | 0.95 | 0.95 | <0.001 | -0.77 | 0.456 | 3.12 (0.10) | 3.12 (0.11) | 0.53 | 0.98 | 0.98 | <0.001 | 0.56 | 0.589 |
| Superior Parietal | 2.72 (0.08) | 2.73 (0.09) | 0.68 | 0.96 | 0.96 | <0.001 | 0.88 | 0.398 | 2.76 (0.07) | 2.76 (0.08) | 0.69 | 0.96 | 0.96 | <0.001 | 1.07 | 0.307 |
| Superior Temporal | 3.26 (0.11) | 3.26 (0.12) | 1.10 | 0.92 | 0.92 | <0.001 | -0.35 | 0.734 | 3.30 (0.09) | 3.31 (0.11) | 0.93 | 0.91 | 0.91 | <0.001 | 0.78 | 0.451 |
| Supramarginal Gyrus | 3.04 (0.11) | 3.04 (0.12) | 1.39 | 0.88 | 0.88 | <0.001 | -0.32 | 0.758 | 3.05 (0.10) | 3.06 (0.10) | 0.78 | 0.96 | 0.96 | <0.001 | 0.49 | 0.635 |
| Transverse Temporal | 2.99 (0.13) | 2.98 (0.14) | 1.21 | 0.94 | 0.94 | <0.001 | -1.06 | 0.310 | 3.05 (0.13) | 3.04 (0.12) | 1.54 | 0.91 | 0.91 | <0.001 | -0.93 | 0.372 |

APD, absolute percentage difference; ICC, intraclass correlation coefficient; lmer, linear mixed-effects model; ROI, region of interest; SD, standard deviation. Note: positive t-values represent cortical thickness increases.

**Supplementary Table 17.** vNav-MPRAGE reliability measures for cortical thickness estimated by the FreeSurfer pipeline.

| ROI | FreeSurfer |  |  |  |  |  |  |  |  |  |  |  |  |  |  |  |
| --- | --- | --- | --- | --- | --- | --- | --- | --- | --- | --- | --- | --- | --- | --- | --- | --- |
|  | Left Hemisphere |  |  |  |  |  |  |  | Right Hemisphere |  |  |  |  |  |  |  |
|  | Timepoint 1<br>Mean (SD) | Timepoint 2<br>Mean (SD) | APD | ICC (2,1) | ICC (3,1) | p-value (ICC) | t-value (lmer) | p-value (lmer) | Timepoint 1<br>Mean (SD) | Timepoint 2<br>Mean (SD) | APD | ICC (2,1) | ICC (3,1) | p-value (ICC) | t-value (lmer) | p-value (lmer) |
| Caudal Anterior Cingulate | 2.73 (0.11) | 2.69 (0.14) | 1.99 | 0.82 | 0.84 | <0.001 | -1.76 | 0.103 | 2.59 (0.20) | 2.57 (0.19) | 3.57 | 0.75 | 0.75 | 0.001 | -0.53 | 0.605 |
| Caudal Middle Frontal | 2.75 (0.07) | 2.75 (0.09) | 1.15 | 0.89 | 0.89 | <0.001 | 0.02 | 0.983 | 2.74 (0.10) | 2.73 (0.10) | 1.18 | 0.91 | 0.91 | <0.001 | -1.00 | 0.339 |
| Cuneus | 2.03 (0.06) | 2.04 (0.06) | 1.68 | 0.74 | 0.74 | 0.001 | 0.32 | 0.756 | 2.05 (0.13) | 2.04 (0.11) | 1.86 | 0.92 | 0.92 | <0.001 | -0.51 | 0.620 |
| Entorhinal Cortex | 3.49 (0.33) | 3.51 (0.24) | 5.27 | 0.74 | 0.74 | 0.001 | 0.24 | 0.813 | 3.74 (0.16) | 3.74 (0.17) | 3.02 | 0.61 | 0.61 | 0.010 | -0.10 | 0.922 |
| Fusiform Gyrus | 2.82 (0.09) | 2.81 (0.10) | 1.19 | 0.91 | 0.92 | <0.001 | -1.65 | 0.125 | 2.86 (0.11) | 2.86 (0.10) | 1.57 | 0.87 | 0.87 | <0.001 | -0.23 | 0.823 |
| Inferior Occipital Cortex | 2.24 (0.09) | 2.24 (0.08) | 1.58 | 0.84 | 0.84 | <0.001 | -0.06 | 0.954 | 2.33 (0.11) | 2.34 (0.09) | 1.76 | 0.88 | 0.88 | <0.001 | 0.69 | 0.502 |
| Inferior Parietal | 2.57 (0.10) | 2.58 (0.11) | 1.39 | 0.92 | 0.92 | <0.001 | 0.80 | 0.440 | 2.65 (0.10) | 2.66 (0.09) | 1.39 | 0.86 | 0.86 | <0.001 | 0.43 | 0.676 |
| Inferior Temporal | 2.96 (0.09) | 2.97 (0.08) | 1.26 | 0.86 | 0.86 | <0.001 | 0.13 | 0.896 | 2.96 (0.10) | 2.95 (0.07) | 1.65 | 0.75 | 0.75 | <0.001 | -0.52 | 0.614 |
| Insula | 3.21 (0.09) | 3.21 (0.10) | 1.41 | 0.83 | 0.83 | <0.001 | -0.15 | 0.885 | 3.23 (0.12) | 3.20 (0.08) | 1.70 | 0.77 | 0.79 | <0.001 | -1.56 | 0.146 |
| Isthmus Cingulate Gyrus | 2.55 (0.15) | 2.53 (0.16) | 3.09 | 0.82 | 0.82 | <0.001 | -0.80 | 0.441 | 2.50 (0.17) | 2.50 (0.19) | 3.72 | 0.82 | 0.82 | <0.001 | 0.00 | 0.996 |
| Lateral Frontal Opercularis | 2.81 (0.10) | 2.80 (0.10) | 1.33 | 0.88 | 0.88 | <0.001 | -0.25 | 0.803 | 2.72 (0.12) | 2.72 (0.11) | 1.39 | 0.92 | 0.92 | <0.001 | 0.01 | 0.991 |
| Lateral Frontal Orbitalis | 2.88 (0.14) | 2.87 (0.12) | 2.05 | 0.83 | 0.83 | <0.001 | -0.69 | 0.502 | 2.83 (0.11) | 2.83 (0.10) | 1.57 | 0.86 | 0.86 | <0.001 | -0.15 | 0.884 |
| Lateral Frontal Triangularis | 2.60 (0.10) | 2.60 (0.11) | 1.56 | 0.88 | 0.88 | <0.001 | 0.17 | 0.865 | 2.59 (0.10) | 2.58 (0.09) | 1.36 | 0.88 | 0.88 | <0.001 | -0.61 | 0.552 |
| Lateral Orbitofrontal | 2.87 (0.09) | 2.89 (0.11) | 2.21 | 0.66 | 0.67 | 0.005 | 1.05 | 0.316 | 2.81 (0.11) | 2.82 (0.10) | 2.45 | 0.68 | 0.68 | 0.004 | 0.51 | 0.621 |
| Lingual Gyrus | 2.18 (0.08) | 2.16 (0.09) | 1.54 | 0.90 | 0.91 | <0.001 | -1.53 | 0.152 | 2.17 (0.10) | 2.17 (0.11) | 1.28 | 0.93 | 0.93 | <0.001 | -0.23 | 0.821 |
| Medial Orbitofrontal | 2.60 (0.13) | 2.61 (0.11) | 3.13 | 0.65 | 0.65 | 0.006 | 0.38 | 0.711 | 2.54 (0.15) | 2.56 (0.19) | 3.40 | 0.78 | 0.78 | <0.001 | 0.66 | 0.521 |
| Middle Temporal | 2.91 (0.12) | 2.92 (0.11) | 1.07 | 0.94 | 0.94 | <0.001 | 0.10 | 0.919 | 3.00 (0.11) | 2.99 (0.12) | 1.21 | 0.93 | 0.93 | <0.001 | -0.62 | 0.548 |
| Paracentral Gyrus | 2.63 (0.18) | 2.63 (0.18) | 1.58 | 0.97 | 0.97 | <0.001 | -0.16 | 0.879 | 2.60 (0.12) | 2.60 (0.14) | 1.36 | 0.93 | 0.93 | <0.001 | -0.23 | 0.819 |
| Parahippocampal | 2.83 (0.23) | 2.81 (0.22) | 1.35 | 0.97 | 0.97 | <0.001 | -1.22 | 0.244 | 2.76 (0.26) | 2.78 (0.27) | 1.53 | 0.98 | 0.98 | <0.001 | 1.39 | 0.191 |
| Pericalcarine | 1.81 (0.15) | 1.79 (0.18) | 3.72 | 0.88 | 0.88 | <0.001 | -0.91 | 0.380 | 1.84 (0.13) | 1.82 (0.12) | 2.78 | 0.88 | 0.89 | <0.001 | -1.46 | 0.171 |
| Postcentral Gyrus | 2.26 (0.13) | 2.26 (0.14) | 1.19 | 0.97 | 0.97 | <0.001 | 0.43 | 0.672 | 2.24 (0.11) | 2.23 (0.11) | 1.43 | 0.94 | 0.94 | <0.001 | -0.96 | 0.357 |
| Posterior Cingulate | 2.57 (0.13) | 2.55 (0.11) | 2.34 | 0.71 | 0.71 | 0.002 | -0.94 | 0.368 | 2.54 (0.09) | 2.54 (0.09) | 1.55 | 0.86 | 0.86 | <0.001 | -0.30 | 0.773 |
| Precentral Gyrus | 2.79 (0.10) | 2.79 (0.12) | 0.90 | 0.96 | 0.96 | <0.001 | 0.02 | 0.987 | 2.78 (0.10) | 2.77 (0.10) | 1.31 | 0.86 | 0.86 | <0.001 | -1.09 | 0.297 |
| Precuneus | 2.47 (0.09) | 2.49 (0.10) | 1.10 | 0.92 | 0.93 | <0.001 | 1.52 | 0.154 | 2.50 (0.06) | 2.49 (0.05) | 1.36 | 0.77 | 0.77 | <0.001 | -0.54 | 0.600 |
| Rostral Anterior Cingulate | 2.88 (0.15) | 2.86 (0.18) | 2.88 | 0.80 | 0.80 | <0.001 | -0.56 | 0.587 | 2.85 (0.14) | 2.84 (0.18) | 3.98 | 0.41 | 0.41 | 0.075 | -0.17 | 0.869 |
| Rostral Middle Frontal | 2.64 (0.09) | 2.64 (0.09) | 1.35 | 0.90 | 0.90 | <0.001 | -0.05 | 0.965 | 2.56 (0.07) | 2.56 (0.07) | 1.53 | 0.75 | 0.75 | <0.001 | -0.11 | 0.916 |
| Superior Frontal Gyrus | 2.86 (0.10) | 2.87 (0.12) | 0.86 | 0.96 | 0.96 | <0.001 | 0.43 | 0.676 | 2.78 (0.10) | 2.77 (0.10) | 1.12 | 0.94 | 0.95 | <0.001 | -1.73 | 0.110 |
| Superior Parietal | 2.33 (0.10) | 2.33 (0.10) | 1.18 | 0.95 | 0.95 | <0.001 | 0.30 | 0.771 | 2.34 (0.11) | 2.35 (0.10) | 1.16 | 0.95 | 0.95 | <0.001 | 0.35 | 0.733 |
| Superior Temporal | 3.00 (0.14) | 3.00 (0.12) | 0.75 | 0.97 | 0.97 | <0.001 | 0.14 | 0.893 | 3.03 (0.11) | 3.03 (0.09) | 1.20 | 0.91 | 0.91 | <0.001 | -0.10 | 0.921 |
| Supramarginal Gyrus | 2.72 (0.13) | 2.72 (0.15) | 0.91 | 0.98 | 0.98 | <0.001 | 0.57 | 0.582 | 2.71 (0.12) | 2.71 (0.13) | 1.62 | 0.91 | 0.91 | <0.001 | -0.18 | 0.859 |
| Transverse Temporal | 2.59 (0.21) | 2.56 (0.18) | 2.03 | 0.94 | 0.94 | <0.001 | -1.56 | 0.145 | 2.61 (0.18) | 2.63 (0.14) | 2.67 | 0.85 | 0.85 | <0.001 | 0.65 | 0.526 |

APD, absolute percentage difference; ICC, intraclass correlation coefficient; lmer, linear mixed-effects model; ROI, region of interest; SD, standard deviation. Note: positive t-values represent cortical thickness increases.

**Supplementary Table 18.** MPRAGE and vNav-MPRAGE similarity measures for cortical thickness estimated by the FreeSurfer pipeline.

| ROI | FreeSurfer |  |  |  |  |  |  |  |  |  |  |  |  |  |  |  |
| --- | --- | --- | --- | --- | --- | --- | --- | --- | --- | --- | --- | --- | --- | --- | --- | --- |
|  | Left Hemisphere |  |  |  |  |  |  |  | Right Hemisphere |  |  |  |  |  |  |  |
|  | MPRAGE Mean (SD) | vNav-MPRAGE Mean (SD) | APD | ICC (2,1) | ICC (3,1) | p-value (ICC) | t-value (lmer) | p-value (lmer) | MPRAGE Mean (SD) | vNav-MPRAGE Mean (SD) | APD | ICC (2,1) | ICC (3,1) | p-value (ICC) | t-value (lmer) | p-value (lmer) |
| Caudal Anterior Cingulate | 2.76 (0.12) | 2.73 (0.11) | 2.06 | 0.83 | 0.85 | <0.001 | -1.53 | 0.152 | 2.58 (0.17) | 2.59 (0.20) | 4.09 | 0.75 | 0.75 | <0.001 | 0.09 | 0.932 |
| Caudal Middle Frontal | 2.75 (0.08) | 2.75 (0.07) | 0.95 | 0.92 | 0.92 | <0.001 | 0.22 | 0.827 | 2.69 (0.11) | 2.74 (0.10) | 2.01 | 0.81 | 0.90 | <0.001 | 3.80 | 0.003 |
| Cuneus | 2.00 (0.06) | 2.03 (0.06) | 2.36 | 0.64 | 0.73 | 0.002 | 2.70 | 0.019 | 2.00 (0.13) | 2.05 (0.13) | 2.77 | 0.83 | 0.89 | <0.001 | 3.02 | 0.011 |
| Entorhinal Cortex | 3.55 (0.26) | 3.49 (0.33) | 3.40 | 0.84 | 0.85 | <0.001 | -1.34 | 0.205 | 3.68 (0.20) | 3.74 (0.16) | 4.43 | 0.28 | 0.28 | 0.163 | 0.92 | 0.376 |
| Fusiform Gyrus | 2.82 (0.11) | 2.82 (0.09) | 1.57 | 0.86 | 0.86 | <0.001 | 0.32 | 0.756 | 2.88 (0.12) | 2.86 (0.11) | 1.37 | 0.92 | 0.92 | <0.001 | -1.04 | 0.321 |
| Inferior Occipital Cortex | 2.26 (0.07) | 2.24 (0.09) | 1.91 | 0.83 | 0.83 | <0.001 | -1.14 | 0.276 | 2.38 (0.08) | 2.33 (0.11) | 2.47 | 0.77 | 0.91 | <0.001 | -5.12 | <0.001* |
| Inferior Parietal | 2.52 (0.10) | 2.57 (0.10) | 2.23 | 0.80 | 0.91 | <0.001 | 4.58 | <0.001* | 2.66 (0.10) | 2.65 (0.10) | 1.04 | 0.93 | 0.93 | <0.001 | -0.85 | 0.410 |
| Inferior Temporal | 2.94 (0.12) | 2.96 (0.09) | 1.93 | 0.78 | 0.78 | <0.001 | 1.27 | 0.230 | 2.94 (0.09) | 2.96 (0.10) | 1.21 | 0.88 | 0.89 | <0.001 | 1.71 | 0.112 |
| Insula | 3.25 (0.08) | 3.21 (0.09) | 1.61 | 0.76 | 0.83 | <0.001 | -2.86 | 0.014 | 3.25 (0.11) | 3.23 (0.12) | 1.26 | 0.92 | 0.93 | <0.001 | -1.58 | 0.139 |
| Isthmus Cingulate Gyrus | 2.60 (0.14) | 2.55 (0.15) | 2.86 | 0.77 | 0.81 | <0.001 | -1.92 | 0.079 | 2.57 (0.18) | 2.50 (0.17) | 3.35 | 0.82 | 0.87 | <0.001 | -2.57 | 0.025 |
| Lateral Frontal Opercularis | 2.80 (0.09) | 2.81 (0.10) | 1.53 | 0.87 | 0.87 | <0.001 | 0.36 | 0.724 | 2.68 (0.13) | 2.72 (0.12) | 1.84 | 0.89 | 0.93 | <0.001 | 3.34 | 0.006 |
| Lateral Frontal Orbitalis | 2.87 (0.14) | 2.88 (0.14) | 1.37 | 0.94 | 0.94 | <0.001 | 1.03 | 0.324 | 2.80 (0.12) | 2.83 (0.11) | 2.08 | 0.84 | 0.85 | <0.001 | 1.47 | 0.168 |
| Lateral Frontal Triangularis | 2.56 (0.10) | 2.60 (0.10) | 2.10 | 0.80 | 0.88 | <0.001 | 3.21 | 0.008 | 2.49 (0.11) | 2.59 (0.10) | 3.97 | 0.61 | 0.88 | <0.001 | 6.91 | <0.001* |
| Lateral Orbitofrontal | 2.86 (0.11) | 2.87 (0.09) | 2.25 | 0.68 | 0.68 | 0.004 | 0.30 | 0.769 | 2.76 (0.12) | 2.81 (0.11) | 2.18 | 0.78 | 0.84 | <0.001 | 2.65 | 0.021 |
| Lingual Gyrus | 2.14 (0.09) | 2.18 (0.08) | 1.87 | 0.82 | 0.88 | <0.001 | 2.93 | 0.013 | 2.12 (0.10) | 2.17 (0.10) | 2.55 | 0.81 | 0.92 | <0.001 | 5.13 | <0.001* |
| Medial Orbitofrontal | 2.49 (0.21) | 2.60 (0.13) | 4.90 | 0.64 | 0.77 | <0.001 | 3.56 | 0.004 | 2.40 (0.18) | 2.54 (0.15) | 7.02 | 0.47 | 0.63 | 0.008 | 3.61 | 0.004 |
| Middle Temporal | 2.90 (0.12) | 2.91 (0.12) | 0.87 | 0.96 | 0.96 | <0.001 | 1.07 | 0.305 | 2.99 (0.12) | 3.00 (0.11) | 1.25 | 0.93 | 0.93 | <0.001 | 0.57 | 0.580 |
| Paracentral Gyrus | 2.63 (0.19) | 2.63 (0.18) | 0.92 | 0.99 | 0.99 | <0.001 | 0.50 | 0.624 | 2.62 (0.13) | 2.6 (0.12) | 1.61 | 0.90 | 0.90 | <0.001 | -1.57 | 0.142 |
| Parahippocampal | 2.84 (0.23) | 2.83 (0.23) | 2.39 | 0.94 | 0.94 | <0.001 | -0.72 | 0.487 | 2.77 (0.28) | 2.76 (0.26) | 1.61 | 0.98 | 0.98 | <0.001 | -0.31 | 0.765 |
| Pericalcarine | 1.73 (0.12) | 1.81 (0.15) | 4.64 | 0.74 | 0.86 | <0.001 | 4.14 | 0.001 | 1.76 (0.11) | 1.84 (0.13) | 4.21 | 0.78 | 0.93 | <0.001 | 6.21 | <0.001* |
| Postcentral Gyrus | 2.22 (0.13) | 2.26 (0.13) | 1.56 | 0.94 | 0.98 | <0.001 | 4.61 | <0.001* | 2.20 (0.10) | 2.24 (0.11) | 1.90 | 0.90 | 0.97 | <0.001 | 5.44 | <0.001* |
| Posterior Cingulate | 2.57 (0.13) | 2.57 (0.13) | 1.72 | 0.89 | 0.89 | <0.001 | -0.13 | 0.897 | 2.59 (0.14) | 2.54 (0.09) | 3.09 | 0.62 | 0.66 | 0.005 | -1.93 | 0.078 |
| Precentral Gyrus | 2.81 (0.11) | 2.79 (0.10) | 1.17 | 0.94 | 0.95 | <0.001 | -2.24 | 0.045 | 2.78 (0.10) | 2.78 (0.10) | 1.12 | 0.93 | 0.93 | <0.001 | 0.07 | 0.942 |
| Precuneus | 2.44 (0.10) | 2.47 (0.09) | 1.64 | 0.86 | 0.90 | <0.001 | 2.54 | 0.026 | 2.50 (0.08) | 2.50 (0.06) | 1.46 | 0.81 | 0.81 | <0.001 | 0.34 | 0.738 |
| Rostral Anterior Cingulate | 2.88 (0.17) | 2.88 (0.15) | 2.43 | 0.83 | 0.83 | <0.001 | 0.00 | 0.998 | 2.87 (0.11) | 2.85 (0.14) | 3.21 | 0.69 | 0.69 | 0.003 | -0.67 | 0.513 |
| Rostral Middle Frontal | 2.60 (0.09) | 2.64 (0.09) | 1.71 | 0.85 | 0.92 | <0.001 | 3.85 | 0.002 | 2.44 (0.09) | 2.56 (0.07) | 4.62 | 0.42 | 0.80 | <0.001 | 7.79 | <0.001* |
| Superior Frontal Gyrus | 2.85 (0.10) | 2.86 (0.10) | 1.05 | 0.94 | 0.95 | <0.001 | 2.03 | 0.066 | 2.72 (0.11) | 2.78 (0.10) | 2.19 | 0.81 | 0.93 | <0.001 | 5.17 | <0.001* |
| Superior Parietal | 2.27 (0.11) | 2.33 (0.10) | 2.69 | 0.80 | 0.93 | <0.001 | 5.68 | <0.001* | 2.31 (0.13) | 2.34 (0.11) | 2.19 | 0.89 | 0.93 | <0.001 | 3.08 | 0.010 |
| Superior Temporal | 3.00 (0.13) | 3.00 (0.14) | 0.77 | 0.98 | 0.98 | <0.001 | -0.72 | 0.486 | 3.05 (0.10) | 3.03 (0.11) | 1.36 | 0.87 | 0.87 | <0.001 | -1.37 | 0.194 |
| Supramarginal Gyrus | 2.69 (0.15) | 2.72 (0.13) | 1.31 | 0.96 | 0.97 | <0.001 | 3.15 | 0.008 | 2.70 (0.13) | 2.71 (0.12) | 1.16 | 0.95 | 0.95 | <0.001 | 0.46 | 0.651 |
| Transverse Temporal | 2.45 (0.23) | 2.59 (0.21) | 5.82 | 0.80 | 0.98 | <0.001 | 10.65 | <0.001* | 2.56 (0.22) | 2.61 (0.18) | 2.36 | 0.90 | 0.92 | <0.001 | 2.14 | 0.054 |

APD, absolute percentage difference; ICC, intraclass correlation coefficient; lmer, linear mixed-effects model; ROI, region of interest; SD, standard deviation. Note: positive t-values represent cortical thickness increases in vNav-MPRAGE. Asterisk indicates significance after Bonferroni correction.

**Supplementary Table 19.** MPRAGE reliability measures for structure volume estimated by the MAGeT-Brain pipeline.

| ROI | MAGeT-Brain |  |  |  |  |  |  |  |  |  |  |  |  |  |  |  |
| --- | --- | --- | --- | --- | --- | --- | --- | --- | --- | --- | --- | --- | --- | --- | --- | --- |
|  | Left Hemisphere |  |  |  |  |  |  |  | Right Hemisphere |  |  |  |  |  |  |  |
|  | Timepoint 1<br>Mean (SD) | Timepoint 2<br>Mean (SD) | APD | ICC (2,1) | ICC (3,1) | p-value (ICC) | t-value (lmer) | p-value (lmer) | Timepoint 1<br>Mean (SD) | Timepoint 2<br>Mean (SD) | APD | ICC (2,1) | ICC (3,1) | p-value (ICC) | t-value (lmer) | p-value (lmer) |
| Amygdala | 1222.85 (136.32) | 1223.39 (135.61) | 1.74 | 0.98 | 0.98 | <0.001 | 0.07 | 0.943 | 1161.00 (131.63) | 1166.46 (121.84) | 2.08 | 0.98 | 0.98 | <0.001 | 0.71 | 0.494 |
| Hippocampus | 2350.08 (233.80) | 2364.31 (231.38) | 0.79 | 1.00 | 1.00 | <0.001 | 2.84 | 0.015 | 2329.92 (303.82) | 2350.77 (249.94) | 2.22 | 0.97 | 0.97 | <0.001 | 1.13 | 0.282 |
| Globus Pallidus | 1610.85 (126.33) | 1608.15 (125.48) | 1.15 | 0.98 | 0.98 | <0.001 | -0.36 | 0.727 | 1436.92 (104.92) | 1435.77 (106.04) | 1.18 | 0.98 | 0.98 | <0.001 | -0.20 | 0.841 |
| Striatum | 9809.77 (936.77) | 9823.69 (932.11) | 0.47 | 1.00 | 1.00 | <0.001 | 0.91 | 0.380 | 9988.92 (857.92) | 9999.77 (846.35) | 0.46 | 1.00 | 1.00 | <0.001 | 0.71 | 0.494 |
| Thalamus | 6294.77 (383.12) | 6275.62 (384.89) | 0.66 | 0.99 | 0.99 | <0.001 | -1.40 | 0.186 | 6124.23 (421.55) | 6125.92 (416.50) | 0.64 | 0.99 | 0.99 | <0.001 | 0.13 | 0.898 |

APD, absolute percentage difference; ICC, intraclass correlation coefficient; lmer, linear mixed-effects model; ROI, region of interest; SD, standard deviation. Note: positive t-values represent structure volume increases.

**Supplementary Table 20.**
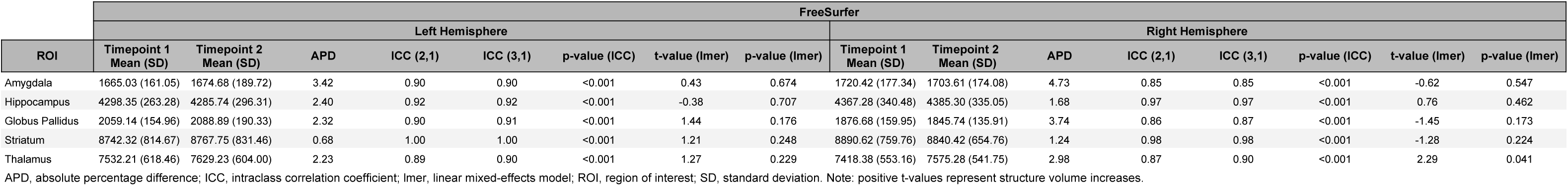
MPRAGE reliability measures for structure volume estimated by the FreeSurfer pipeline.

**Supplementary Table 21.** vNav-MPRAGE reliability measures for structure volume estimated by the MAGeT-Brain pipeline.

| ROI | MAGEt-Brain |  |  |  |  |  |  |  |  |  |  |  |  |  |  |  |
| --- | --- | --- | --- | --- | --- | --- | --- | --- | --- | --- | --- | --- | --- | --- | --- | --- |
|  | Left Hemisphere |  |  |  |  |  |  |  | Right Hemisphere |  |  |  |  |  |  |  |
|  | Timepoint 1<br>Mean (SD) | Timepoint 2<br>Mean (SD) | APD | ICC (2,1) | ICC (3,1) | p-value (ICC) | t-value (lmer) | p-value (lmer) | Timepoint 1<br>Mean (SD) | Timepoint 2<br>Mean (SD) | APD | ICC (2,1) | ICC (3,1) | p-value (ICC) | t-value (lmer) | p-value (lmer) |
| Amygdala | 1216.46 (139.91) | 1226.92 (126.80) | 3.33 | 0.93 | 0.93 | <0.001 | 0.75 | 0.468 | 1159.23 (126.28) | 1160.23 (114.89) | 3.09 | 0.94 | 0.94 | <0.001 | 0.08 | 0.936 |
| Hippocampus | 2371.00 (256.56) | 2379.08 (259.55) | 1.16 | 0.99 | 0.99 | <0.001 | 0.79 | 0.447 | 2353.69 (331.67) | 2359.54 (339.74) | 0.91 | 1.00 | 1.00 | <0.001 | 0.74 | 0.474 |
| Globus Pallidus | 1631.77 (120.63) | 1634.69 (117.85) | 1.17 | 0.98 | 0.98 | <0.001 | 0.39 | 0.703 | 1454.23 (101.37) | 1459.39 (108.23) | 1.86 | 0.94 | 0.94 | <0.001 | 0.50 | 0.628 |
| Striatum | 9801.85 (960.31) | 9820.23 (942.25) | 0.49 | 1.00 | 1.00 | <0.001 | 1.08 | 0.299 | 10001.85 (861.24) | 9984.08 (880.28) | 0.51 | 1.00 | 1.00 | <0.001 | -1.10 | 0.294 |
| Thalamus | 6283.08 (400.22) | 6266.00 (405.69) | 0.84 | 0.99 | 0.99 | <0.001 | -1.03 | 0.322 | 6107.08 (444.83) | 6102.62 (428.25) | 0.68 | 0.99 | 0.99 | <0.001 | -0.29 | 0.775 |

APD, absolute percentage difference; ICC, intraclass correlation coefficient; lmer, linear mixed-effects model; ROI, region of interest; SD, standard deviation. Note: positive t-values represent structure volume increases.

**Supplementary Table 22.** vNav-MPRAGE reliability measures for structure volume estimated by the FreeSurfer pipeline.

| ROI | FreeSurfer |  |  |  |  |  |  |  |  |  |  |  |  |  |  |  |
| --- | --- | --- | --- | --- | --- | --- | --- | --- | --- | --- | --- | --- | --- | --- | --- | --- |
|  | Left Hemisphere |  |  |  |  |  |  |  | Right Hemisphere |  |  |  |  |  |  |  |
|  | Timepoint 1<br>Mean (SD) | Timepoint 2<br>Mean (SD) | APD | ICC (2,1) | ICC (3,1) | p-value (ICC) | t-value (lmer) | p-value (lmer) | Timepoint 1<br>Mean (SD) | Timepoint 2<br>Mean (SD) | APD | ICC (2,1) | ICC (3,1) | p-value (ICC) | t-value (lmer) | p-value (lmer) |
| Amygdala | 1654.08 (174.95) | 1616.63 (160.55) | 4.70 | 0.87 | 0.88 | <0.001 | -1.64 | 0.128 | 1746.14 (201.62) | 1684.74 (172.65) | 4.17 | 0.90 | 0.94 | <0.001 | -3.46 | 0.005* |
| Hippocampus | 4242.25 (279.35) | 4234.42 (255.99) | 1.88 | 0.92 | 0.92 | <0.001 | -0.26 | 0.801 | 4310.73 (343.31) | 4293.28 (326.36) | 1.47 | 0.97 | 0.97 | <0.001 | -0.79 | 0.443 |
| Globus Pallidus | 2075.32 (172.61) | 2094.14 (162.96) | 3.42 | 0.82 | 0.82 | <0.001 | 0.66 | 0.521 | 1915.75 (133.19) | 1923.96 (134.92) | 4.31 | 0.73 | 0.73 | 0.002 | 0.29 | 0.776 |
| Striatum | 8635.66 (868.91) | 8632.25 (859.65) | 1.45 | 0.98 | 0.98 | <0.001 | -0.06 | 0.950 | 8702.45 (843.11) | 8754.83 (726.22) | 2.19 | 0.90 | 0.90 | <0.001 | 0.53 | 0.606 |
| Thalamus | 7800.26 (583.18) | 7871.18 (691.86) | 2.04 | 0.92 | 0.92 | <0.001 | 0.97 | 0.351 | 7655.42 (564.99) | 7690.08 (649.06) | 2.36 | 0.93 | 0.93 | <0.001 | 0.55 | 0.595 |

APD, absolute percentage difference; ICC, intraclass correlation coefficient; lmer, linear mixed-effects model; ROI, region of interest; SD, standard deviation. Note: positive t-values represent structure volume increases. Asterisk indicates significance after Bonferroni correction.

**Supplementary Table 23.**
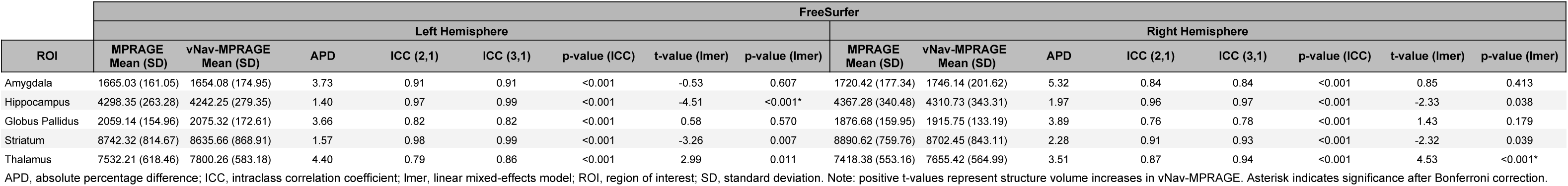
MPRAGE and vNav-MPRAGE similarity measures for structure volume estimated by the FreeSurfer pipeline.

**Supplementary Table 24.** Relationships between signal-to-noise ratio and cortical thickness, as estimated by the CIVET and FreeSurfer pipelines.

| ROI | CIVET |  |  |  | FreeSurfer |  |  |  |
| --- | --- | --- | --- | --- | --- | --- | --- | --- |
|  | Left Hemisphere |  | Right Hemisphere |  | Left Hemisphere |  | Right Hemisphere |  |
|  | t-value (lmer) | p-value (lmer) | t-value (lmer) | p-value (lmer) | t-value (lmer) | p-value (lmer) | t-value (lmer) | p-value (lmer) |
| Caudal Anterior Cingulate | 4.65 | <0.001* | 3.20 | 0.007 | 6.95 | <0.001* | 3.51 | 0.004 |
| Caudal Middle Frontal | 1.73 | 0.109 | 2.14 | 0.053 | 2.77 | 0.017 | -0.03 | 0.980 |
| Cuneus | -1.06 | 0.311 | -0.28 | 0.783 | -1.22 | 0.245 | -2.42 | 0.032 |
| Entorhinal Cortex | -0.05 | 0.962 | -0.90 | 0.384 | -0.80 | 0.439 | -1.33 | 0.207 |
| Fusiform Gyrus | 1.65 | 0.125 | 2.85 | 0.014 | -0.92 | 0.375 | 0.82 | 0.429 |
| Inferior Occipital Cortex | -2.88 | 0.014 | 1.04 | 0.320 | 2.50 | 0.028 | 1.49 | 0.161 |
| Inferior Parietal | -1.52 | 0.153 | 4.71 | <0.001* | -3.73 | 0.003 | 1.61 | 0.132 |
| Inferior Temporal | 4.35 | 0.001 | 5.55 | <0.001* | 2.38 | 0.034 | 2.85 | 0.014 |
| Insula | 1.94 | 0.076 | 5.17 | <0.001* | 3.22 | 0.007 | 4.68 | <0.001* |
| Isthmus Cingulate Gyrus | 1.42 | 0.180 | 3.85 | 0.002 | 1.10 | 0.294 | 2.54 | 0.026 |
| Lateral Frontal Opercularis | 2.10 | 0.057 | 3.49 | 0.004 | 2.92 | 0.013 | 1.20 | 0.253 |
| Lateral Frontal Orbitalis | 2.51 | 0.027 | 4.08 | 0.001 | -1.19 | 0.256 | 1.37 | 0.194 |
| Lateral Frontal Triangularis | 3.07 | 0.010 | 1.60 | 0.134 | 0.90 | 0.384 | 0.05 | 0.964 |
| Lateral Orbitofrontal | 5.39 | <0.001* | 6.55 | <0.001* | 2.06 | 0.061 | 0.97 | 0.352 |
| Lingual Gyrus | 2.44 | 0.030 | 2.03 | 0.063 | -0.79 | 0.442 | -1.79 | 0.098 |
| Medial Orbitofrontal | 6.60 | <0.001* | 6.22 | <0.001* | 2.87 | 0.013 | 1.78 | 0.099 |
| Middle Temporal | 0.76 | 0.461 | 6.54 | <0.001* | -1.07 | 0.304 | 2.21 | 0.047 |
| Paracentral Gyrus | 0.00 | 0.997 | -0.44 | 0.665 | -1.40 | 0.186 | -1.07 | 0.303 |
| Parahippocampal | 1.22 | 0.242 | 0.31 | 0.764 | -0.77 | 0.456 | -1.64 | 0.126 |
| Pericalcarine | -1.25 | 0.233 | 0.29 | 0.776 | -4.14 | 0.001 | -3.65 | 0.003 |
| Postcentral Gyrus | -3.56 | 0.004 | -1.56 | 0.144 | -1.65 | 0.125 | -2.92 | 0.013 |
| Posterior Cingulate | -0.64 | 0.533 | 1.08 | 0.302 | 2.13 | 0.055 | 0.28 | 0.783 |
| Precentral Gyrus | -1.56 | 0.143 | 1.09 | 0.296 | 0.52 | 0.613 | -0.18 | 0.860 |
| Precuneus | -2.29 | 0.040 | 1.38 | 0.191 | -3.19 | 0.008 | -2.09 | 0.057 |
| Rostral Anterior Cingulate | 4.80 | <0.001* | 4.59 | <0.001* | 2.19 | 0.048 | 1.46 | 0.168 |
| Rostral Middle Frontal | 3.52 | 0.004 | 3.92 | 0.002 | 2.29 | 0.041 | -0.17 | 0.871 |
| Superior Frontal Gyrus | 2.26 | 0.043 | 2.64 | 0.021 | 2.56 | 0.025 | 0.71 | 0.491 |
| Superior Parietal | -3.98 | 0.002 | -1.12 | 0.286 | -5.12 | <0.001* | -3.03 | 0.010 |
| Superior Temporal | 3.18 | 0.008 | 6.08 | <0.001* | 0.06 | 0.954 | 0.50 | 0.625 |
| Supramarginal Gyrus | 0.10 | 0.922 | 0.80 | 0.437 | 0.02 | 0.982 | -0.56 | 0.586 |
| Transverse Temporal | 3.87 | 0.002 | 3.54 | 0.004 | -1.79 | 0.098 | -1.35 | 0.201 |

lmer, linear mixed-effects model; ROI, region of interest. Note: positive t-values represent cortical thickness increases. Asterisk indicates significance after Bonferroni correction.

**Supplementary Table 25.** Relationships between signal-to-noise ratio and structure volume, as estimated by the MAGeT-Brain and FreeSurfer pipelines.

| ROI | MAGeT-Brain |  |  |  | FreeSurfer |  |  |  |
| --- | --- | --- | --- | --- | --- | --- | --- | --- |
|  | Left Hemisphere |  | Right Hemisphere |  | Left Hemisphere |  | Right Hemisphere |  |
|  | t-value (lmer) | p-value (lmer) | t-value (lmer) | p-value (lmer) | t-value (lmer) | p-value (lmer) | t-value (lmer) | p-value (lmer) |
| Amygdala | -4.47 | <0.001* | -1.67 | 0.121 | -5.87 | <0.001* | -4.58 | <0.001* |
| Hippocampus | -2.68 | 0.020 | -3.40 | 0.005 | 1.95 | 0.071 | -0.52 | 0.611 |
| Globus Pallidus | -5.13 | <0.001* | -3.47 | 0.005* | -7.03 | <0.001* | -4.64 | <0.001* |
| Striatum | 1.69 | 0.117 | -1.47 | 0.166 | -5.49 | <0.001* | -7.43 | <0.001* |
| Thalamus | 9.42 | <0.001* | 11.03 | <0.001* | 2.28 | 0.039 | 2.82 | 0.014 |

lmer, linear mixed-effects model; ROI, region of interest. Note: positive t-values represent cortical thickness increases. Asterisk indicates significance after Bonferroni correction.

## References

Ai, L., Craddock, R.C., Tottenham, N., Dyke, J.P., Colcombe, S., Milham, M., Franco, A.R., 2019. Is it Time to Switch Your T1W Sequence? Assessing the Impact of Prospective Motion Correction on the Reliability and Quality of Structural Imaging. bioRxiv 666289. https://doi.org/10.1101/666289

Alexander-Bloch, A., Clasen, L., Stockman, M., Ronan, L., Lalonde, F., Giedd, J., Raznahan, A., 2016. Subtle in-scanner motion biases automated measurement of brain anatomy from in vivo MRI. Hum. Brain Mapp. 37, 2385–2397.

Amaral, R.S.C., Park, M.T.M., Devenyi, G.A., Lynn, V., Pipitone, J., Winterburn, J., Chavez, S., Schira, M., Lobaugh, N.J., Voineskos, A.N., Pruessner, J.C., Chakravarty, M.M., Alzheimer’s Disease Neuroimaging Initiative, 2018. Manual segmentation of the fornix, fimbria, and alveus on high-resolution 3T MRI: Application via fully-automated mapping of the human memory circuit white and grey matter in healthy and pathological aging. Neuroimage 170, 132–150.

Andersen, M., Björkman-Burtscher, I.M., Marsman, A., Petersen, E.T., Boer, V.O., 2019. Improvement in diagnostic quality of structural and angiographic MRI of the brain using motion correction with interleaved, volumetric navigators. PLoS One 14, e0217145.

Avants, B.B., Epstein, C.L., Grossman, M., Gee, J.C., 2008. Symmetric diffeomorphic image registration with cross-correlation: evaluating automated labeling of elderly and neurodegenerative brain. Med. Image Anal. 12, 26–41.

Bedford, S.A., Park, M.T.M., Devenyi, G.A., Tullo, S., Germann, J., Patel, R., Anagnostou, E., Baron-Cohen, S., Bullmore, E.T., Chura, L.R., Craig, M.C., Ecker, C., Floris, D.L., Holt, R.J., Lenroot, R., Lerch, J.P., Lombardo, M.V., Murphy, D.G.M., Raznahan, A., Ruigrok, A.N.V., Smith, E., Spencer, M.D., Suckling, J., Taylor, M.J., Thurm, A., MRC AIMS Consortium, Lai, M.-C., Chakravarty, M.M., 2020. Large-scale analyses of the relationship between sex, age and intelligence quotient heterogeneity and cortical morphometry in autism spectrum disorder. Mol. Psychiatry 25, 614–628.

Cannon, T.D., Chung, Y., He, G., Sun, D., Jacobson, A., van Erp, T.G.M., McEwen, S., Addington, J., Bearden, C.E., Cadenhead, K., Cornblatt, B., Mathalon, D.H., McGlashan, T., Perkins, D., Jeffries, C., Seidman, L.J., Tsuang, M., Walker, E., Woods, S.W., Heinssen, R., North American Prodrome Longitudinal Study Consortium, 2015. Progressive reduction in cortical thickness as psychosis develops: a multisite longitudinal neuroimaging study of youth at elevated clinical risk. Biol. Psychiatry 77, 147–157.

Casey, B.J., Cannonier, T., Conley, M.I., Cohen, A.O., Barch, D.M., Heitzeg, M.M., Soules, M.E., Teslovich, T., Dellarco, D.V., Garavan, H., Orr, C.A., Wager, T.D., Banich, M.T., Speer, N.K., Sutherland, M.T., Riedel, M.C., Dick, A.S., Bjork, J.M., Thomas, K.M., Chaarani, B., Mejia, M.H., Hagler, D.J., Jr, Daniela Cornejo, M., Sicat, C.S., Harms, M.P., Dosenbach, N.U.F., Rosenberg, M., Earl, E., Bartsch, H., Watts, R., Polimeni, J.R., Kuperman, J.M., Fair, D.A., Dale, A.M., ABCD Imaging Acquisition Workgroup, 2018. The Adolescent Brain Cognitive Development (ABCD) study: Imaging acquisition across 21 sites. Dev. Cogn. Neurosci. 32, 43–54.

Chakravarty, M.M., Bertrand, G., Hodge, C.P., Sadikot, A.F., Collins, D.L., 2006. The creation of a brain atlas for image guided neurosurgery using serial histological data. Neuroimage 30, 359–376.

Chakravarty, M.M., Steadman, P., van Eede, M.C., Calcott, R.D., Gu, V., Shaw, P., Raznahan, A., Collins, D.L., Lerch, J.P., 2013. Performing label-fusion-based segmentation using multiple automatically generated templates. Hum. Brain Mapp. 34, 2635–2654.

Chincarini, A., Sensi, F., Rei, L., Gemme, G., Squarcia, S., Longo, R., Brun, F., Tangaro, S., Bellotti, R., Amoroso, N., Bocchetta, M., Redolfi, A., Bosco, P., Boccardi, M., Frisoni, G.B., Nobili, F., Alzheimer’s Disease Neuroimaging Initiative, 2016. Integrating longitudinal information in hippocampal volume measurements for the early detection of Alzheimer’s disease. Neuroimage 125, 834–847.

Cicchetti, D.V., 1994. Guidelines, criteria, and rules of thumb for evaluating normed and standardized assessment instruments in psychology. Psychol. Assess. 6, 284–290.

Collins, D.L., Neelin, P., Peters, T.M., Evans, A.C., 1994. Automatic 3D intersubject registration of MR volumetric data in standardized Talairach space. J. Comput. Assist. Tomogr. 18, 192–205.

Coupé, P., Manjón, J.V., Gedamu, E., Arnold, D., Robles, M., Collins, D.L., 2009. An object-based method for Rician noise estimation in MR images. Med. Image Comput. Comput. Assist. Interv. 12, 601–608.

Dale, A.M., Fischl, B., Sereno, M.I., 1999. Cortical surface-based analysis. I. Segmentation and surface reconstruction. Neuroimage 9, 179–194.

de Graaf, R.A., 2019. In Vivo NMR Spectroscopy: Principles and Techniques. John Wiley & Sons.

Fischl, B., Salat, D.H., Busa, E., Albert, M., Dieterich, M., Haselgrove, C., van der Kouwe, A., Killiany, R., Kennedy, D., Klaveness, S., Montillo, A., Makris, N., Rosen, B., Dale, A.M., 2002. Whole brain segmentation: automated labeling of neuroanatomical structures in the human brain. Neuron 33, 341–355.

Fischl, B., Sereno, M.I., Dale, A.M., 1999. Cortical surface-based analysis. II: Inflation, flattening, and a surface-based coordinate system. Neuroimage 9, 195–207.

Fjell, A.M., Grydeland, H., Krogsrud, S.K., Amlien, I., Rohani, D.A., Ferschmann, L., Storsve, A.B., Tamnes, C.K., Sala-Llonch, R., Due-Tønnessen, P., Bjørnerud, A., Sølsnes, A.E., Håberg, A.K., Skranes, J., Bartsch, H., Chen, C.-H., Thompson, W.K., Panizzon, M.S., Kremen, W.S., Dale, A.M., Walhovd, K.B., 2015. Development and aging of cortical thickness correspond to genetic organization patterns. Proc. Natl. Acad. Sci. U. S. A. 112, 15462–15467.

Gruetter, R., Tkác, I., 2000. Field mapping without reference scan using asymmetric echo-planar techniques. Magn. Reson. Med. 43, 319–323.

Jack, C.R., Knopman, D.S., Jagust, W.J., Petersen, R.C., Weiner, M.W., Aisen, P.S., Shaw, L.M., Vemuri, P., Wiste, H.J., Weigand, S.D., Lesnick, T.G., Pankratz, V.S., Donohue, M.C., Trojanowski, J.Q., 2013. Tracking pathophysiological processes in Alzheimer’s disease: an updated hypothetical model of dynamic biomarkers. The Lancet Neurology 12, 207–216.

Kim, J.S., Singh, V., Lee, J.K., Lerch, J., Ad-Dab’bagh, Y., MacDonald, D., Lee, J.M., Kim, S.I., Evans, A.C., 2005. Automated 3-D extraction and evaluation of the inner and outer cortical surfaces using a Laplacian map and partial volume effect classification. Neuroimage 27, 210–221.

Klein, A., Tourville, J., 2012. 101 Labeled Brain Images and a Consistent Human Cortical Labeling Protocol. Frontiers in Neuroscience 6, 1–12.

Lee, H., Nakamura, K., Narayanan, S., Brown, R.A., Arnold, D.L., Alzheimer’s Disease Neuroimaging Initiative, 2019. Estimating and accounting for the effect of MRI scanner changes on longitudinal whole-brain volume change measurements. Neuroimage 184, 555–565.

Lerch, J.P., Evans, A.C., 2005. Cortical thickness analysis examined through power analysis and a population simulation. Neuroimage 24, 163–173.

Makowski, C., Béland, S., Kostopoulos, P., Bhagwat, N., Devenyi, G.A., Malla, A.K., Joober, R., Lepage, M., Chakravarty, M.M., 2018. Evaluating accuracy of striatal, pallidal, and thalamic segmentation methods: Comparing automated approaches to manual delineation. Neuroimage 170, 182–198.

Mekle, R., Mlynárik, V., Gambarota, G., Hergt, M., Krueger, G., Gruetter, R., 2009. MR spectroscopy of the human brain with enhanced signal intensity at ultrashort echo times on a clinical platform at 3T and 7T. Magn. Reson. Med. 61, 1279–1285.

Pardoe, H.R., Kucharsky Hiess, R., Kuzniecky, R., 2016. Motion and morphometry in clinical and nonclinical populations. Neuroimage 135, 177–185.

Pipitone, J., Park, M.T.M., Winterburn, J., Lett, T.A., Lerch, J.P., Pruessner, J.C., Lepage, M., Voineskos, A.N., Chakravarty, M.M., Alzheimer’s Disease Neuroimaging Initiative, 2014. Multi-atlas segmentation of the whole hippocampus and subfields using multiple automatically generated templates. Neuroimage 101, 494–512.

Potvin, O., Khademi, A., Chouinard, I., Farokhian, F., Dieumegarde, L., Leppert, I., Hoge, R., Rajah, M.N., Bellec, P., Duchesne, S., CIMA-Q group, CCNA group, 2019. Measurement Variability Following MRI System Upgrade. Front. Neurol. 10, 726.

Provencher, S.W., 2001. Automatic quantitation of localized in vivo 1H spectra with LCModel. NMR Biomed. 14, 260–264.

Raznahan, A., Shaw, P.W., Lerch, J.P., Clasen, L.S., Greenstein, D., Berman, R., Pipitone, J., Chakravarty, M.M., Giedd, J.N., 2014. Longitudinal four-dimensional mapping of subcortical anatomy in human development. Proc. Natl. Acad. Sci. U. S. A. 111, 1592–1597.

Reardon, P.K., Seidlitz, J., Vandekar, S., Liu, S., Patel, R., Park, M.T.M., Alexander-Bloch, A., Clasen, L.S., Blumenthal, J.D., Lalonde, F.M., Giedd, J.N., Gur, R., Gur, R., Lerch, J.P., Chakravarty, M.M., Satterthwaite, T., Shinohara, R.T., Raznahan, A., 2018. Normative brain size variation and brain shape diversity in humans. Science 360, 1222–1227.

Reuter, M., Tisdall, M.D., Qureshi, A., Buckner, R.L., van der Kouwe, A.J.W., Fischl, B., 2015. Head motion during MRI acquisition reduces gray matter volume and thickness estimates. Neuroimage 107, 107–115.

Schmaal, L., Veltman, D.J., van Erp, T.G.M., Sämann, P.G., Frodl, T., Jahanshad, N., Loehrer, E., Tiemeier, H., Hofman, A., Niessen, W.J., Vernooij, M.W., Ikram, M.A., Wittfeld, K., Grabe, H.J., Block, A., Hegenscheid, K., Völzke, H., Hoehn, D., Czisch, M., Lagopoulos, J., Hatton, S.N., Hickie, I.B., Goya-Maldonado, R., Krämer, B., Gruber, O., Couvy-Duchesne, B., Rentería, M.E., Strike, L.T., Mills, N.T., de Zubicaray, G.I., McMahon, K.L., Medland, S.E., Martin, N.G., Gillespie, N.A., Wright, M.J., Hall, G.B., MacQueen, G.M., Frey, E.M., Carballedo, A., van Velzen, L.S., van Tol, M.J., van der Wee, N.J., Veer, I.M., Walter, H., Schnell, K., Schramm, E., Normann, C., Schoepf, D., Konrad, C., Zurowski, B., Nickson, T., McIntosh, A.M., Papmeyer, M., Whalley, H.C., Sussmann, J.E., Godlewska, B.R., Cowen, P.J., Fischer, F.H., Rose, M., Penninx, B.W.J.H., Thompson, P.M., Hibar, D.P., 2016. Subcortical brain alterations in major depressive disorder: findings from the ENIGMA Major Depressive Disorder working group. Mol. Psychiatry 21, 806–812.

Shuter, B., Yeh, I.B., Graham, S., Au, C., Wang, S.-C., 2008. Reproducibility of brain tissue volumes in longitudinal studies: effects of changes in signal-to-noise ratio and scanner software. Neuroimage 41, 371–379.

Siemens, 2020. MAGNETOM Trio Upgrade [WWW Document]. URL https://www.siemens-healthineers.com/magnetic-resonance-imaging/options-and-upgrades/upgrades/magnetom-trio-upgrade/features

Simpson, R., Devenyi, G.A., Jezzard, P., Hennessy, T.J., Near, J., 2017. Advanced processing and simulation of MRS data using the FID appliance (FID-A)-An open source, MATLAB-based toolkit. Magn. Reson. Med. 77, 23–33.

Sowell, E.R., Peterson, B.S., Kan, E., Woods, R.P., Yoshii, J., Bansal, R., Xu, D., Zhu, H., Thompson, P.M., Toga, A.W., 2007. Sex differences in cortical thickness mapped in 176 healthy individuals between 7 and 87 years of age. Cereb. Cortex 17, 1550–1560.

Takao, H., Hayashi, N., Ohtomo, K., 2013. Effects of the use of multiple scanners and of scanner upgrade in longitudinal voxel-based morphometry studies. J. Magn. Reson. Imaging 38, 1283–1291.

Tisdall, M.D., Dylan Tisdall, M., Hess, A.T., Reuter, M., Meintjes, E.M., Fischl, B., van der Kouwe, A.J.W., 2012. Volumetric navigators for prospective motion correction and selective reacquisition in neuroanatomical MRI. Magnetic Resonance in Medicine 68, 389–399.

Tisdall, M.D., Reuter, M., Qureshi, A., Buckner, R.L., Fischl, B., van der Kouwe, A.J.W., 2016. Prospective motion correction with volumetric navigators (vNavs) reduces the bias and variance in brain morphometry induced by subject motion. Neuroimage 127, 11–22.

Tkác, I., Starcuk, Z., Y. Choi, I., Gruetter, R., 1999. In vivo 1H NMR spectroscopy of rat brain at 1 ms echo time. Magnetic Resonance in Medicine 41, 649–656.

Treadway, M.T., Waskom, M.L., Dillon, D.G., Holmes, A.J., Park, M.T.M., Chakravarty, M.M., Dutra, S.J., Polli, F.E., Iosifescu, D.V., Fava, M., Gabrieli, J.D.E., Pizzagalli, D.A., 2015. Illness progression, recent stress, and morphometry of hippocampal subfields and medial prefrontal cortex in major depression. Biol. Psychiatry 77, 285–294.

Tullo, S., Devenyi, G.A., Patel, R., Park, M.T.M., Collins, D.L., Chakravarty, M.M., 2018. Warping an atlas derived from serial histology to 5 high-resolution MRIs. Sci Data 5, 180107.

Tullo, S., Patel, R., Devenyi, G.A., Salaciak, A., Bedford, S.A., Farzin, S., Wlodarski, N., Tardif, C.L., Breitner, J.C.S., Mallar Chakravarty, M., the PREVENT-AD Research Group, 2019. MR-based age-related effects on the striatum, globus pallidus, and thalamus in healthy individuals across the adult lifespan. Human Brain Mapping 40, 5269–5288.

Tustison, N.J., Avants, B.B., Cook, P.A., Zheng, Y., Egan, A., Yushkevich, P.A., Gee, J.C., 2010. N4ITK: improved N3 bias correction. IEEE Trans. Med. Imaging 29, 1310–1320.

van Erp, T.G.M., Hibar, D.P., Rasmussen, J.M., Glahn, D.C., Pearlson, G.D., Andreassen, O.A., Agartz, I., Westlye, L.T., Haukvik, U.K., Dale, A.M., Melle, I., Hartberg, C.B., Gruber, O., Kraemer, B., Zilles, D., Donohoe, G., Kelly, S., McDonald, C., Morris, D.W., Cannon, D.M., Corvin, A., Machielsen, M.W.J., Koenders, L., de Haan, L., Veltman, D.J., Satterthwaite, T.D., Wolf, D.H., Gur, R.C., Gur, R.E., Potkin, S.G., Mathalon, D.H., Mueller, B.A., Preda, A., Macciardi, F., Ehrlich, S., Walton, E., Hass, J., Calhoun, V.D., Bockholt, H.J., Sponheim, S.R., Shoemaker, J.M., van Haren, N.E.M., Pol, H.E.H., Ophoff, R.A., Kahn, R.S., Roiz-Santiañez, R., Crespo-Facorro, B., Wang, L., Alpert, K.I., Jönsson, E.G., Dimitrova, R., Bois, C., Whalley, H.C., McIntosh, A.M., Lawrie, S.M., Hashimoto, R., Thompson, P.M., Turner, J.A., 2016. Subcortical brain volume abnormalities in 2028 individuals with schizophrenia and 2540 healthy controls via the ENIGMA consortium. Mol. Psychiatry 21, 585.

van Erp, T.G.M., Walton, E., Hibar, D.P., Schmaal, L., Jiang, W., Glahn, D.C., Pearlson, G.D., Yao, N., Fukunaga, M., Hashimoto, R., Okada, N., Yamamori, H., Bustillo, J.R., Clark, V.P., Agartz, I., Mueller, B.A., Cahn, W., de Zwarte, S.M.C., Hulshoff Pol, H.E., Kahn, R.S., Ophoff, R.A., van Haren, N.E.M., Andreassen, O.A., Dale, A.M., Doan, N.T., Gurholt, T.P., Hartberg, C.B., Haukvik, U.K., Jørgensen, K.N., Lagerberg, T.V., Melle, I., Westlye, L.T., Gruber, O., Kraemer, B., Richter, A., Zilles, D., Calhoun, V.D., Crespo-Facorro, B., Roiz-Santiañez, R., Tordesillas-Gutiérrez, D., Loughland, C., Carr, V.J., Catts, S., Cropley, V.L., Fullerton, J.M., Green, M.J., Henskens, F.A., Jablensky, A., Lenroot, R.K., Mowry, B.J., Michie, P.T., Pantelis, C., Quidé, Y., Schall, U., Scott, R.J., Cairns, M.J., Seal, M., Tooney, P.A., Rasser, P.E., Cooper, G., Shannon Weickert, C., Weickert, T.W., Morris, D.W., Hong, E., Kochunov, P., Beard, L.M., Gur, R.E., Gur, R.C., Satterthwaite, T.D., Wolf, D.H., Belger, A., Brown, G.G., Ford, J.M., Macciardi, F., Mathalon, D.H., O’Leary, D.S., Potkin, S.G., Preda, A., Voyvodic, J., Lim, K.O., McEwen, S., Yang, F., Tan, Y., Tan, S., Wang, Z., Fan, F., Chen, J., Xiang, H., Tang, S., Guo, H., Wan, P., Wei, D., Bockholt, H.J., Ehrlich, S., Wolthusen, R.P.F., King, M.D., Shoemaker, J.M., Sponheim, S.R., De Haan, L., Koenders, L., Machielsen, M.W., van Amelsvoort, T., Veltman, D.J., Assogna, F., Banaj, N., de Rossi, P., Iorio, M., Piras, F., Spalletta, G., McKenna, P.J., Pomarol-Clotet, E., Salvador, R., Corvin, A., Donohoe, G., Kelly, S., Whelan, C.D., Dickie, E.W., Rotenberg, D., Voineskos, A.N., Ciufolini, S., Radua, J., Dazzan, P., Murray, R., Reis Marques, T., Simmons, A., Borgwardt, S., Egloff, L., Harrisberger, F., Riecher-Rössler, A., Smieskova, R., Alpert, K.I., Wang, L., Jönsson, E.G., Koops, S., Sommer, I.E.C., Bertolino, A., Bonvino, A., Di Giorgio, A., Neilson, E., Mayer, A.R., Stephen, J.M., Kwon, J.S., Yun, J.-Y., Cannon, D.M., McDonald, C., Lebedeva, I., Tomyshev, A.S., Akhadov, T., Kaleda, V., Fatouros-Bergman, H., Flyckt, L., Karolinska Schizophrenia Project, Busatto, G.F., Rosa, P.G.P., Serpa, M.H., Zanetti, M.V., Hoschl, C., Skoch, A., Spaniel, F., Tomecek, D., Hagenaars, S.P., McIntosh, A.M., Whalley, H.C., Lawrie, S.M., Knöchel, C., Oertel-Knöchel, V., Stäblein, M., Howells, F.M., Stein, D.J., Temmingh, H.S., Uhlmann, A., Lopez-Jaramillo, C., Dima, D., McMahon, A., Faskowitz, J.I., Gutman, B.A., Jahanshad, N., Thompson, P.M., Turner, J.A., 2018. Cortical Brain Abnormalities in 4474 Individuals With Schizophrenia and 5098 Control Subjects via the Enhancing Neuro Imaging Genetics Through Meta Analysis (ENIGMA) Consortium. Biol. Psychiatry 84, 644–654.

van Haren, N.E.M., Schnack, H.G., Cahn, W., van den Heuvel, M.P., Lepage, C., Collins, L., Evans, A.C., Hulshoff Pol, H.E., Kahn, R.S., 2011. Changes in cortical thickness during the course of illness in schizophrenia. Arch. Gen. Psychiatry 68, 871–880.

van Rooij, D., Anagnostou, E., Arango, C., Auzias, G., Behrmann, M., Busatto, G.F., Calderoni, S., Daly, E., Deruelle, C., Di Martino, A., Dinstein, I., Duran, F.L.S., Durston, S., Ecker, C., Fair, D., Fedor, J., Fitzgerald, J., Freitag, C.M., Gallagher, L., Gori, I., Haar, S., Hoekstra, L., Jahanshad, N., Jalbrzikowski, M., Janssen, J., Lerch, J., Luna, B., Martinho, M.M., McGrath, J., Muratori, F., Murphy, C.M., Murphy, D.G.M., O’Hearn, K., Oranje, B., Parellada, M., Retico, A., Rosa, P., Rubia, K., Shook, D., Taylor, M., Thompson, P.M., Tosetti, M., Wallace, G.L., Zhou, F., Buitelaar, J.K., 2018. Cortical and Subcortical Brain Morphometry Differences Between Patients With Autism Spectrum Disorder and Healthy Individuals Across the Lifespan: Results From the ENIGMA ASD Working Group. American Journal of Psychiatry 175, 359–369.

Voineskos, A.N., Winterburn, J.L., Felsky, D., Pipitone, J., Rajji, T.K., Mulsant, B.H., Chakravarty, M.M., 2015. Hippocampal (subfield) volume and shape in relation to cognitive performance across the adult lifespan. Hum. Brain Mapp. 36, 3020–3037.

Wijtenburg, S.A., Yang, S., Fischer, B.A., Rowland, L.M., 2015. In vivo assessment of neurotransmitters and modulators with magnetic resonance spectroscopy: application to schizophrenia. Neurosci. Biobehav. Rev. 51, 276–295.

Winterburn, J.L., Pruessner, J.C., Chavez, S., Schira, M.M., Lobaugh, N.J., Voineskos, A.N., Chakravarty, M.M., 2013. A novel in vivo atlas of human hippocampal subfields using high-resolution 3 T magnetic resonance imaging. Neuroimage 74, 254–265.

Zijdenbos, A.P., Forghani, R., Evans, A.C., 2002. Automatic “pipeline” analysis of 3-D MRI data for clinical trials: application to multiple sclerosis. IEEE Trans. Med. Imaging 21, 1280– 1291.

